# Species-specific loss of genetic diversity and inbreeding following agricultural intensification

**DOI:** 10.1101/2024.10.07.616612

**Authors:** Zachary J. Nolen, Patrycja Jamelska, Ana Sofia Torres Lara, Niklas Wahlberg, Anna Runemark

## Abstract

Land-use change from agricultural intensification is a major driver of global insect declines. We investigated the genetic consequences of grassland decline in Sweden by sequencing museum and modern specimens of three focal and five additional Polyommatinae butterfly species. From 59 historical and 90 contemporary genomes, we find a 6% decline in genetic diversity and increased isolation and inbreeding in the grassland specialist *Cyaniris semiargus* over the past 70 years. In contrast, generalist *Polyommatus icarus* and heathland specialist *Plebejus argus* maintained genetic diversity and connectivity. Although currently genomic erosion is mild, we infer it lags considerably larger declines in effective population sizes. Using simulations, we demonstrate that only a small portion of genetic diversity is lost in early stages of decline, creating a genetic extinction debt. Two of five additional species show similar reductions, underscoring that restoration of grassland habitat is necessary to restore gene flow and halt further genomic erosion in grassland insects.

## Introduction

Insect decline has received increased attention in recent years as reports uncover alarming declines in insect assemblages and biomass (*1*). Efforts to quantify this decline estimate that terrestrial insects are declining at an average rate of 11% per decade globally, with this trend being primarily driven by even stronger regional declines in Europe and North America (*2*). This stark decline is the result of a variety of anthropogenic pressures, primarily climate change and land use changes such as agricultural intensification and urbanization (*1*, *3–7*). In Europe, formerly biodiverse semi-natural grasslands have been abandoned or converted into intensively cultivated land (*8*), placing a strong pressure on grassland insect species (*6*).

While insect declines have alarming implications for the natural communities and ecosystem services they play key roles in, declines also lead to genomic erosion within insect populations themselves. As populations decline, the relative strength of drift increases (*9*), eroding genetic diversity, thereby reducing adaptive potential (*10*, *11*). Additionally, in small, isolated populations, deleterious mutations are more likely to accumulate through drift and become unmasked through inbreeding (*12–17*), further reinforcing population decline (*18*, *19*). Mitigation of genomic erosion that threatens population persistence has been made a part of Goal A of the Kunming-Montréal Global Biodiversity Framework, which stipulates maintenance of genetic diversity to preserve the adaptive potential of wild populations (*20*).

Monitoring approaches to assess genomic erosion are under development. While plans to monitor genetic diversity in the wild have been implemented in several countries (*21–24*), interpreting measures of genetic diversity remains a challenge. First, genomic erosion lags disturbance events, and current estimates of genetic diversity will therefore not capture the genetic extinction debt of populations (*25–30*). Therefore, identifying measures that enable early detection of genomic erosion is a priority. Second, absolute levels of genetic diversity are influenced by a multitude of factors not related to threat level and vary considerably across taxa and populations (*31–34*). To control for variation in baseline levels of genetic diversity, programs in several countries supplement estimates of contemporary effective population sizes with measures of genetic diversity metrics across multiple time points (*22*, *35*).

Temporal estimates of genetic diversity enable direct quantification of genomic erosion but are limited to detecting changes after the first sampled time point. To overcome this, genomic resources from historic samples preserved in natural history museums provide an opportunity to establish baseline estimates of genetic diversity from before major anthropogenically driven insect declines (*24*, *36*). Using comparisons to museum specimens, populations already undergoing genomic erosion can be identified with only one contemporary sampling point, reducing time to detection of populations at risk.

Here, we use museomics to address to what extent genetic diversity and connectivity has been lost in grassland associated butterflies during the second wave of agricultural modernization in Sweden (*37*, *38*). Starting in the 1940s, small farms were abandoned, and their associated semi-natural grasslands began to decline considerably (*38*). We examine three focal butterfly species with Swedish Red List statuses of ‘least concern’ (*39*) in this landscape. These species vary in habitat specialization, ranging from grassland generalist to specialist, enabling us to examine if patterns of genetic diversity decline are stronger in more specialized species, in line with expectations from greater demographic declines in specialist insect abundance compared to generalists (*40–42*). We assess genetic change by comparing museum and contemporary estimates of genetic diversity and use simulations to assess if the lag in loss of diversity is expected given the scale of demographic change. We further examine if changes in genetic diversity are common across grassland butterfly species by extending the dataset to include historical and modern individuals of five additional related butterfly species with Swedish Red List statuses ranging from least concern to endangered (*39*). In the face of ongoing mass extinction, identifying species and populations at risk at an early stage of decline is paramount. Understanding early signals of genomic erosion is key for intervention before substantial diversity is irreversibly lost. Here, by combining a large set of historical and contemporary specimens, we uncover a case of ongoing genomic erosion in response to recent anthropogenic change and highlight metrics that respond quickly to population decline, providing insights into genomic erosion at its early stages when intervention is most effective.

## Results

We analyzed changes in genetic diversity, differentiation, and proxies for genetic load in three focal Polyommatinae (Lepidoptera: Lycaenidae) butterfly species: *Polyommatus icarus,* a grassland generalist, *Plebejus argus*, a heathland specialist, and *Cyaniris semiargus*, a grassland specialist (Figure 1A). We generated whole genome resequencing data from pinned museum specimens from the entomological collections of the Biological Museum at Lund University (*43*) collected from 1917-1956 (Table S1; *N* = 28 - *Po. icarus*; 12 - *Pl. argus*; 14 - *Cy. semiargus*) in southern Sweden. These individuals were sampled as populations by grouping specimens generally collected at the same locality during the same year and compared with WGS data from samples collected at corresponding localities in 2020-2023 by Nolen et al (*44*) (Figure 1B; Table S1; *N* = 27 - *Po. icarus*; 29 - *Pl. argus*; 29 - *Cy. semiargus*). As DNA extracted from historical specimens is fragmented and subject to post-mortem DNA damage (*45*, *46*) (Figure S1), we removed transitions from all analyses. As the shorter fragment size of historical samples can result in poor mapping of reads containing non-reference alleles, we constructed variation graphs for each reference genome containing high quality variants called in the modern samples. Mapping both sample types to these variation graphs reduced reference bias compared to aligning to the linear reference genome, as well as reduced the disparity in reference bias between historical and modern samples, especially when calling genotypes (Figure S2&3; Supplementary Methods). The reduction in reference bias and its influence on downstream analyses is clearly visible in population structure analyses, where using the linear reference alignments resulted in samples clustering primarily by sample type (historical or modern), while using the variation graph alignments resulted in samples clustering primarily by sampling locality (Figure S4). While the variation graph alignment method reduced reference bias in historical samples, it was not eliminated, and results should be considered with this in mind, especially for *Po. icarus* which exhibits the largest residual bias (Figure S2&4). Median sequencing depth was 6.4x (range = 2.2 to 16.7x) for historical samples and 19.3x (range = 7.0 to 56.7x; Table S1) for modern samples. To account for variation in sequencing depth, all analyses were performed with data subsampled to between 6 and 8x sequencing depth.

**Figure 1.**
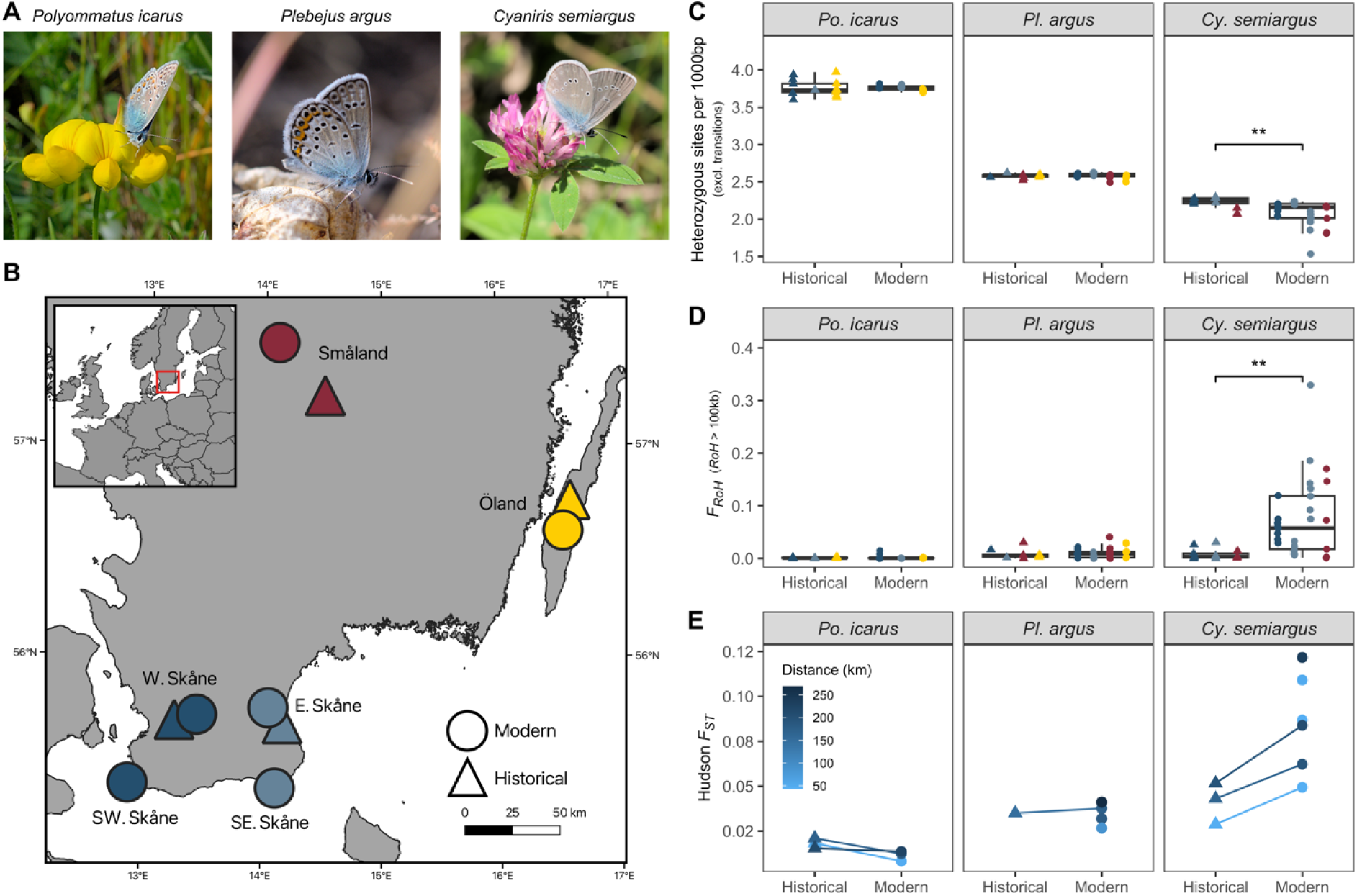
Sampling design and changes in genetic diversity, differentiation, and inbreeding in the focal species. (A) Images of the three focal species. (B) Map of the sampling area in southern Sweden, sampling locations are approximate. Circles represent areas with modern samples in at least one of the species and triangles represent areas with historical samples, see Table S1 for complete sampling information. (C) Change in individual heterozygosity over time, measured as heterozygous sites per 1000bp, excluding transitions. Each point represents an individual sample, grouped into historical and modern samples per species. Coloring refers to sampling region and corresponds to the map in panel B. (D) Change in inbreeding coefficients over time, estimated using runs of homozygosity larger than 100kb (F_RoH_). (E) Change in pairwise population genetic differentiation, _ST_ over time between equivalent comparisons in each time period. Color of the points and line refer to the geographic distance between populations. Populations with no connecting line refer to pairs of modern sample populations that have no comparable historic pair. In C & D, results of linear mixed models assessing the effect of time period on the response are shown using the following scale, where panels without * are not significant (P > 0.05): * P <= 0.05, ** P <= 0.01, *** P <= 0.001.

### Reduced genetic diversity and effective population size in Cy. semiargus

To assess if agricultural intensification and the associated reduction in semi-natural grassland habitat has resulted in reduced genetic diversity, we compared individual genome-wide heterozygosity of historical and modern samples within each species. Individual heterozygosity is significantly reduced in the specialist *Cy. semiargus* (percent difference: −6%; LMM: *P* = 0.008; Figure 1C; Table S2&3), but not in *Po. icarus* or *Pl. argus*. Population specific changes in genetic diversity estimated both from mean heterozygosity and nucleotide diversity (Table S4) suggest that genetic diversity has mostly been reduced in populations in Skåne, where arable land cover is greatest, with only a 1% change in nucleotide diversity in Småland (Tables S5&6).

To understand the recent demographic patterns contributing to contemporary genetic diversity, we assessed changes in effective population size (*N_e_*) over the past century using the modern genomes with a linkage disequilibrium (LD) based approach. We find stable and very high effective population sizes in *Po. icarus* and fluctuating or only recently declined sizes in *Pl. argus*, in line with their maintenance of genetic diversity (Figure S5; Table S6). In *Cy. semargus*, we find that all populations in Skåne have gone through population declines within the past 30 generations, with current effective population sizes of 35 (SE Skåne), 202 (W Skåne), and 520 (E Skåne) (Figure S5; Table S6).

### Reduced connectivity may drive loss of genetic diversity

While genetic diversity decline is an expected product of demographic decline, it can also be a product of reductions in gene flow (*9*). Modern *Cy. semiargus* populations exhibit stronger population structure than historical ones, based on both genetic differentiation and clustering approaches. For population pairs represented in both modern and historical samplings, modern estimates of genetic differentiation (*F*_ST_) in *Cy. semiargus* have increased from an average of 0.041 to 0.065, with the increase being largest between W Skåne and Småland (Figure 1E; Table S7). Including population pairs in the modern sampling that are not represented in the historical, but that include similarly geographically separated populations, increases the average *F*_ST_ of modern *Cy. semiargus* pairs to 0.086. For *Po. icarus* and *Pl. argus*, average *F*_ST_ between pairs represented in both time periods has changed from 0.018 to 0.011 and 0.034 to 0.038 (Figure 1E; Table S7). When including additional modern pairs in *Pl. argus*, the modern average is slightly lower, 0.035, though this includes pairs at short distances that are not represented in historical pairs. Historical population pairs are sampled at different time points, with pairs including populations sampled up to 20 years apart (Table S7), which could inflate historical *F*_ST_ estimates.

Principal component analyses (PCAs) and admixture analyses also reveal stronger population structure in modern *Cy. semiargus* samples than historical (Figure 2A). Each modern population forms a distinct cluster in the specialist *Cy. semiargus*, with modern Småland samples clustering with their historical counterparts (Figure 2B). Historical samples from Western and Eastern Skåne cluster together in a separate cluster from the respective modern samples, highlighting that the stronger genetic structure found in modern samples is largely driven by isolation of populations in the agricultural landscapes within Sweden’s southernmost province, Skåne (Figure 1D; Figure 2A & B; Figure S6). Populations of both *Po. icarus* and *Pl. argus* from Skåne separate from those sampled elsewhere in southern Sweden along the major axis of variation, with historical samples clustering with geographically proximal modern samples. Admixture analyses support only one genetic cluster in each of these species (Table S8), but the correlation of residuals suggest additional genetic clusters in *Pl. argus* (Figure S7), consistent with the PCA. This highlights that while genetic structure likely exists, the variation is too low for reliable clustering at these sample sizes. We additionally find that historical and modern samples separate out on PC2 in *Po. icarus*, potentially an artifact of residual reference bias in historical samples of this species (Figure S2). The general pattern of historical populations clustering in the PCA with at least one geographically proximal modern population suggests that historical samples are representative of ancestral populations to the modern ones.

**Figure 2.**
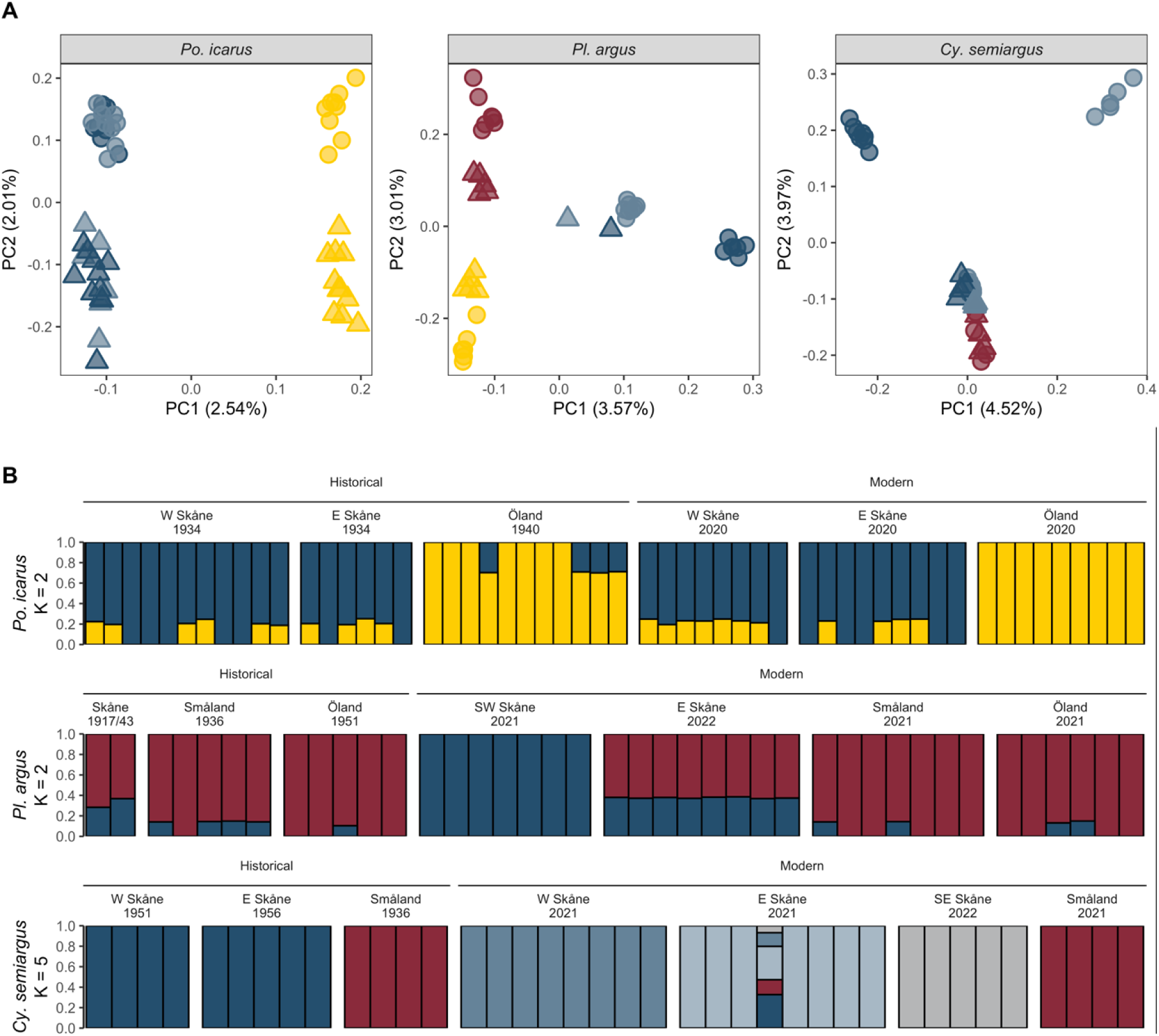
Population structure of the focal species. (A) Principal Component Analysis (PCA) of the three focal species. Shapes refer to sampling period, with each circle representing a modern and each triangle a historical individual, with colors corresponding to the sampling locations in Figure 1B. Percentages on each axis refer to the proportion of total variance explained by the component. (B) Admixture analyses for the three focal species. Each color represents one of K genetic clusters, with bars representing individuals, colored by their probability of belonging to a given cluster. For Po. icarus and Pl. argus, the best fit was at K = 1, but K = 2 is shown for illustrative purposes. For Cy. semiargus, the best fi K = 5 is shown.

### Increased inbreeding in specialist Cy. semiargus

To investigate changes in rates of inbreeding, we estimated the inbreeding coefficient *F_RoH_*, which is the proportion of the autosomal genome in runs of homozygosity (*RoH*) greater than a certain length (*47*). We quantified *F_RoH_* for *RoH* longer than 100kb, corresponding roughly to *RoH* with shared ancestry in the past 200 generations (*48*). We found a significant increase in *F_RoH_* from historical to modern *Cy. semiargus* samples (difference: 0.065, LMM: *P* < 0.01; Figure 1D; Table S9). Across species, the highest inbreeding coefficient for a historical sample is 0.031, with historical species averages all below 0.01 (Tables S2&9). In *Cy. semiargus*, modern samples have a maximum coefficient of 0.329 and an average coefficient of 0.073, an order of magnitude higher than the mean for historical samples. Individual inbreeding coefficients were largely determined by runs of homozygosity >= 1.5 Mb in length (Figure 3A), corresponding to low effective population sizes in the past ∼12 generations, in line with estimates of recently low effective population size in our demographic analyses. In *Po. icarus* and *Pl. argus*, both modern and historical estimates of *F_RoH_* are low, with averages <= 0.01 and no evidence of significant increases (Table S9).

**Figure 3.**
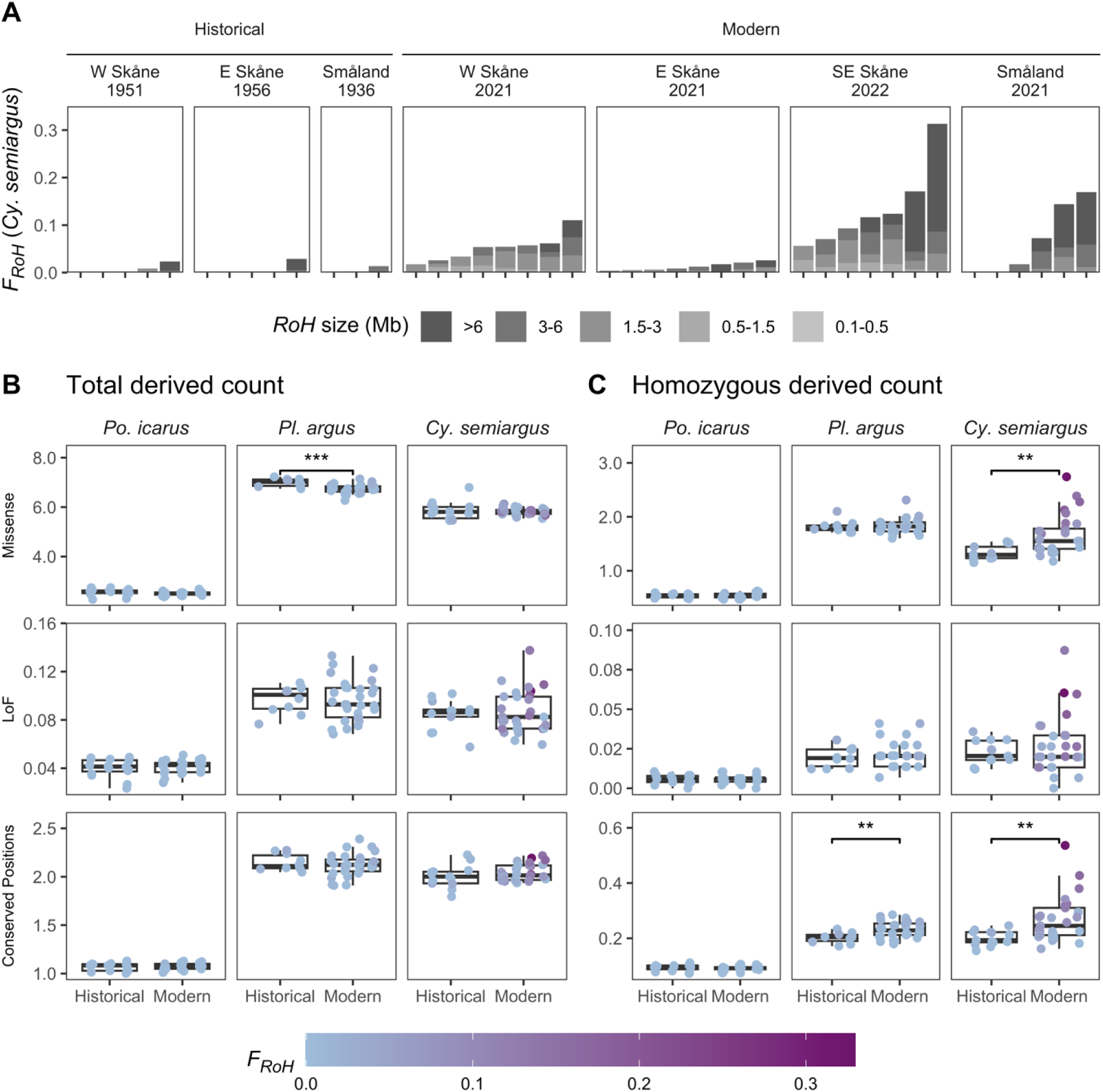
Temporal changes in derived count of putatively deleterious mutations in the focal species. (A) Individual inbreeding coefficients (F_RoH_) partitioned by RoH size class. Longer RoH indicate more recent common ancestry of the identical haplotypes. (B) Total count of derived alleles per 1000 called genotypes for individuals of the three focal species at positions classifying three categories of putatively deleterious mutations: I) missense (i.e. weakly deleterious; upper), II) loss of function (LoF, i.e. strongly deleterious; middle), and III) conserved positions (presumed strongly deleterious; lower). C) Count of derived alleles in homozygous state per 1000 called genotypes for the same mutation categories as in B. In B & C, results of linear mixed models with burden as a response to time period are shown in each panel, where panels without * are not significant (P > 0.05): * P <= 0.05, ** P <= 0.01, *** P <= 0.001.

To assess if reductions in heterozygosity explained by increases in *F_RoH_* match theoretical expectations, we performed coalescent simulations parameterized on the recent demographic histories inferred from the modern genomic data, estimating heterozygosity and *F_RoH_*for individuals sampled at time points corresponding to our modern and historical samplings. In all simulations, genome-wide heterozygosity decline in this time frame was primarily due to the presence of long runs of homozygosity, rather than an even reduction in heterozygosity across the genome (Table S10). To assess if we observed this same pattern in the empirical genomic data, we adjusted estimates of individual heterozygosity by the inbreeding coefficient *F_RoH_* per individual. Individual heterozygosity outside *RoH* is not different between time periods (Figure S8), agreeing with the simulation predictions. Additionally, for all simulated populations except for Småland *Cy. semiargus*, the estimated ratios of modern/historical heterozygosity from the genomic data fell within ranges predicted by the simulations, as did average estimates of *F_RoH_* (Figure S9). In the case of Småland *Cy. semiargus*, we estimated a reduction in heterozygosity and increase in *F_RoH_* from the genomic data, but no change was predicted by the simulations. As the *RoH* estimated from the modern sample suggest some level of recent demographic decline that is absent from the inferred demographic history, this may reflect a limitation of the inferred history for this population used to parameterize the simulations.

### Increased homozygosity of weakly deleterious mutations in Cy. semiargus

Population decline and subdivision can increase the exposure of partially recessive, deleterious mutations to selection due to increased homozygosity, facilitating purging of strongly deleterious mutations (*15*, *49*, *50*). However, as populations decline and subdivide, selection weakens relative to drift, allowing weakly deleterious mutations to drift to higher frequencies and even fix, increasing genetic load (*14*, *51*, *52*).

Here, as a proxy for genetic load, we estimated deleterious mutation burden per individual using derived counts of putatively deleterious mutations (as in e.g. *53*–*57*). Deleterious mutations were identified based off the functional annotation or the sequence conservation score of the position and divided into three categories: (I) missense, i.e. weakly deleterious; (II) loss of function (LoF), i.e. strongly deleterious; and (III) conserved, i.e. with high sequence conservation across Lycaenidae, presumed to be strongly deleterious. We partitioned these counts in to total derived counts, approximating load if mutations are primarily additive, and homozygous derived counts, reflecting load if mutations are primarily recessive.

We find no evidence of changes in the total derived count for any of the three mutation categories in *Po. icarus* or *Cy. semiargus*. However, in *Pl. argus*, total derived count at missense positions is reduced (LMM: *P* < 0.001; Figure 3B; Table S10), but there is no significant change in any other mutation category or at either intergenic or synonymous genic positions (Table S11). However, homozygous derived count has increased at missense and conserved positions in *Cy. semiargus* (LMM: *P* = 0.002 and 0.006; Figure 3C; Table S10), and at conserved positions in *Pl. argus* (LMM: *P* = 0.006; Figure 3B; Table S10), but not changed over time in *Po. icarus. Cyaniris semiargus* individuals with higher inbreeding coefficients also had higher homozygous derived counts across all categories of deleterious mutations (LM: *R^2^* = 0.83, 0.40, 0.76 for missense, LoF, and conserved positions, respectively; Figure S10). There is also a weaker increase of total derived count with inbreeding coefficient at LoF and conserved positions (LM: *R^2^* = 0.12, 0.12, respectively). There is no change in homozygous derived count at LoF positions over time in any species.

### Genetic diversity indicators

We utilized the genetic diversity indicator thresholds outlined by Andersson et al. (*21*) to assess if genomic erosion is reflected in conservation status in the three focal species as well as historical-modern pairs of five additional species of Polyommatinae butterfly (Table S1). The thresholds suggest categorizations of ‘Acceptable’, ‘Warning’, and ‘Alarm’ rates of genetic erosion, reflecting >=95%, 75-94%, and <75% retention of genetic diversity over 100 years, respectively (*21*). Based on changes in mean population heterozygosity and nucleotide diversity, populations of all three species largely fall in the ‘Acceptable’ category, except for *Cy. semiargus* in Southeast and Western Skåne, which fall into ‘Warning’ category (Figure 4; Tables S5&6). The same indicator framework in Sweden categorizes effective population sizes >= 500 as ‘Acceptable’, 51 - 499 as ‘Warning’, and <= 50 as ‘Alarm’ (*21*, *58*).

**Figure 4.**
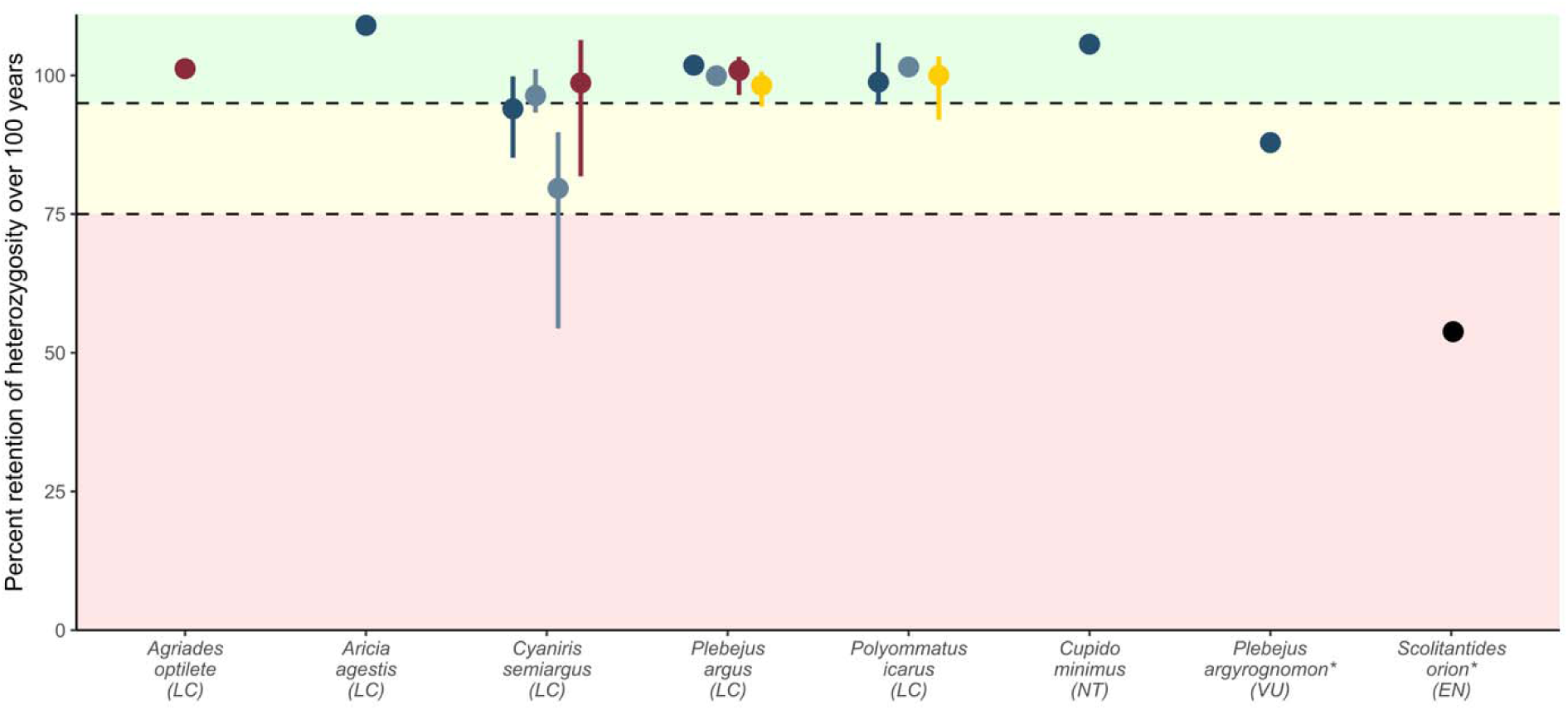
Changes in heterozygosity projected over 100 years based on the rate of change estimated from modern and historical samples for each of 8 species. Species are grouped by conservation status, which follows the species name on the x-axis. Background coloration refers to the three genetic decline thresholds outlined by Andersson et al (21), where a retention of >=95% over 100 year is considered ‘acceptable’ (green), a retention of 75-94% as ‘warning’ (yellow), and a retention of <75% as ‘alarm’ (red). Points represent median estimates for change in heterozygosity across all historical/modern individual pairs for a species. Species with bars have more than one possible sample pair, with error bars illustrating the spread of estimates across all sample pairs. *Sc. orion and Pl. argyrognomon include pinned museum specimens for both the modern and historical individuals. The modern individuals for Pl. argyrognomon and Sc. orion were sampled in 2003 and 2016, respectively, and the conservation statuses shown here reflect the statuses at those times. Plebejus argyrognomon is additionally the only species aligned to a reference genome for a different species (Table S12).

Under this indicator, all *Po. icarus* and *Pl. argus* populations and Småland *Cy. semiargus* would be classed as ‘Acceptable’. The low effective population sizes in Skåne for *Cy. semiargus* would place Eastern Skåne as ‘Acceptable’, but bordering on ‘Warning’, Western Skåne as ‘Warning’, and Southeastern Skåne as ‘Alarm’ (Table S6).

We also assessed if the conservation status of the five additional species, ranging from ‘Least Concern’ to ‘Endangered’ (*39*) (Table S12) are associated with genetic erosion. As a single pair of individuals were sampled from each additional species, we reassessed the focal three species, comparing all possible combinations of historical and modern individuals for each population. We find that ‘Least Concern’ species largely show ‘Acceptable’ changes in genetic diversity, except for Southeastern and Western Skåne *Cy. semiargus*, which fall into the ‘Warning’ category. Of the Red Listed species, *Cupido minimus* (‘Near Threatened’) falls into the ‘Acceptable’ category, *Plebejus argyrognomon* (‘Vulnerable’ in 2003 when the modern individual in this study was collected; ‘Critically Endangered’ and presumed extinct in Sweden presently (*59*)) into ‘Warning’, and *Scolitantides orion* (‘Endangered’) into ‘Alarm’ (Figure 4; Table S6). Results from our focal species illustrate that estimates from single individuals are challenging to interpret, as estimates can vary considerably depending on the individual pair (Figure 4).

## Discussion

Our analyses of 59 historical and 90 modern genomes uncover a decline in genetic diversity in the specialist grassland butterfly *Cyaniris semiargus* over the past century. The previously connected populations of this species in southern Sweden have become isolated into distinct genetic clusters following agricultural intensification. Increased rates of inbreeding have begun to unmask weakly deleterious mutations in *Cy. semiargus*, potentially reducing individual and mean fitness. This contrasts to the generalist *Polyommatus icarus*, where we detect minimal changes in genetic diversity, inbreeding, and genetic differentiation over the focal period. While the decline in genetic diversity and increased homozygosity of deleterious mutations in *Cy. semiargus* is relatively mild, such genomic erosion lags demographic change (*25*, *28*, *30*). We infer that effective population sizes have decreased dramatically in some *Cy. semiargus* populations and that genetic diversity will continue to decline even if these populations stabilize at their current sizes. All three species share a status of ‘Least Concern’ in Sweden, highlighting that assessing genetic erosion can identify genetic extinction debts in species and populations not captured by census-based conservation assessments.

The low genetic differentiation and absence of significant increases in population subdivision in *Po. icarus* and *Pl. argus* suggest gene flow maintains cohesive metapopulations in these species. Consistently, levels of heterozygosity remain stable, and inbreeding remains low in all populations. In contrast, we uncover a loss of population connectivity in *Cy. semiargus*, evidenced by increased population structure in the clustering analysis and higher genetic differentiation among modern populations. The historically lower levels of genetic differentiation across southern Sweden in *Cy. semiargus* suggest that currently isolated populations in Skåne are subsets of a historically well-connected metapopulation. The similarly high estimates of historical effective population size in West- and Eastern Skåne provide an independent line of evidence supporting this interpretation. The concurrent abrupt declines in effective population sizes may reflect the cessation of gene flow, shifting effective size estimates from those of the metapopulation to estimates for each subpopulation. This pattern is consistent with theoretical expectations of high genetic diversity being maintained by even low levels of gene flow (*60*, *61*), as documented in another Lycaenid butterfly (*62*). Hence, concerns about genetic rescue (*63*) wiping out locally adapted ancestry (*64*, *65*) are not likely to be relevant for *Cy. semiargus* and other grassland associated insects in regions with historically abundant habitat. In general, estimates of historical population structure may prove valuable for distinguishing whether contemporary population structure reflects species ecology or is a consequence of anthropogenic habitat isolation.

In the specialist *Cy. semiargus* we document an on average 6% reduction in heterozygosity. This change is connected to increases in the rate of inbreeding, with differences between modern and historical individual heterozygosity largely explained by changes in the abundance of long runs of homozygosity.

Thus, *F_RoH_* can provide a powerful metric for detecting genomic erosion even from only contemporary samples. Additionally, trajectories of recent effective population sizes, inferred from LD, have the potential to uncover recent demographic declines before changes in genetic diversity begin. For instance, temporal changes in effective population size suggest that *Pl. argus* has experienced very recent declines, though not to sizes as low as *Cy. semiargus*, but with no changes in heterozygosity or *F_RoH_*. Hence, there is a possibility that there have been recent reductions in gene flow also in *Pl. argus*, despite stable estimates of differentiation, as population differentiation lags reductions in gene flow (*26*, *66*). Ultimately, while the observed reductions in genetic diversity in these species are relatively mild, genetic diversity loss lags demographic change and will continue even if population sizes stabilize.

Understanding to what extent genetic diversity decline, resulting from habitat loss and isolation, reduces the fitness of individuals and prospects for maintaining long-term viable populations remains a challenge. As we have no direct fitness estimates available, we quantified counts of putatively deleterious mutations as a proxy for genetic load (*67*). We found little evidence for either purging or accumulation of deleterious mutations in any of the three species, as total derived counts did not differ significantly for most mutation categories. We found a small decrease (3.7%) in the total derived count of missense mutations in *Pl. argus*, which is surprising, as theory predicts weakly deleterious mutations to be purged less efficiently than more strongly deleterious ones (*15*, *49*). As there is no change in the derived count at loss of function or conserved positions, which are expected to be more strongly deleterious, this decrease is potentially an artifact.

In *Cy. semiargus*, increased rates of inbreeding have increased the homozygous derived count at missense and conserved positions. Assuming at least partial recessivity for these mutations, this would increase the realized load in more inbred populations, even if the total count of derived alleles has not changed. However, a small increase in homozygosity at missense positions may not contribute to substantial fitness effects, especially if these alleles tend to be additive. Surprisingly, the increased homozygous derived count at conserved positions, expected to have relatively strong deleterious effects (*12*, *68*), more closely mirrors patterns at missense than loss of function positions. This could reflect the limited ability of sequence conservation scores to distinguish between weakly and strongly deleterious mutations (*69*, *70*). In addition, the metric we utilize to assess sequence conservation, Genomic Evolutionary Rate Profiling (GERP) scores (*68*), depends on estimates of the rate of neutral evolution, approximating the rejected substitution rate. Our GERP score estimates are based on a time tree (similar to *54*, *71*, *72*) and hence differ from scores estimated based on neutral rates (e.g. *73*). Nevertheless, our identification of conserved sites selects for the most conserved variants in each dataset and should be robust for within-species comparisons.

While it is unclear to what extent the genomic erosion we estimate in *Cy. semiargus* populations affects fitness in the current generation, the patterns we observe are in line with a lag between population decline and subsequent genomic erosion. While we measure only an average 6% reduction in individual heterozygosity across the study period, we infer greater than 10-fold declines in effective population sizes in recent decades. Using coalescent simulations, we demonstrate that this disparity is to be expected for such recent declines, implying that genetic erosion will continue to ‘catch up’ to the scale of demographic decline in future generations. Recent empirical and simulation studies (*27–29*, *74*) have highlighted a time lag between disturbance events and genomic erosion referred to as drift debt (*16*, *75*) or genetic extinction debt (*25*). Importantly, such a lag is not only expected in the loss of genome-wide genetic diversity, but also in increases in the rate of inbreeding and accumulation of genetic load (*28*). It can take many generations for drift and inbreeding to bring low frequency deleterious mutations to high enough frequency for purging and accumulation of deleterious variants to be noticeable (*28*), so it is not surprising we do not detect these patterns. Given time, populations will lose a portion of their deleterious burden through drift and purging, but a portion also will drift to higher frequency and fixation in subsequent generations, reducing mean fitness of the population in the long term (*12–14*, *16*, *76*). Simultaneously, erosion of genome-wide genetic diversity will continuously reduce adaptive potential (*77*), further reducing the prospects of long-term persistence. Stabilization of populations at current effective population sizes is insufficient to reduce the genetic extinction debt they now carry. Increasing the levels of gene flow to historic levels to increase effective population sizes is needed to reduce this debt.

We assessed if the reduction in heterozygosity and increases in inbreeding found in *Cy. semiargus* are also found in other species of Polyommatinae blues in this region. We find that genetic diversity is maintained in five of the seven additional species assessed. Under the threat categories used in Sweden for genetic monitoring, we uncover ‘Warning’ and ‘Alarm’ level classifications for two additional species, both endangered grassland specialists, *Pl. argyrognomon* and *Sc. orion*. The estimated level of heterozygosity decline in *Pl. argyrognomon* across a period from 1967-2003 is comparable to the decline in *Cy. semiargus*. This is concerning, as when the modern *Pl. argyrognomon* was sampled, the species already had a ‘Vulnerable’ conservation status and is presumed extinct in Sweden since 2018 (*59*).

None of the three focal species are currently demographically declining at a rate to qualify for the Swedish Red List based on census estimates. Systematic monitoring in Sweden from 2010-2023 reveal declines in both *Po. icarus* and *Cy. semiargus*, while *Pl. argus* appears to have stabilized (*78*). In *Cy. semiargus*, we find that demographic decline has been sufficient to begin reducing genetic diversity, but that most genomic erosion has yet to be realized. Genetic monitoring, especially monitoring incorporating historical samples, can enable early identification of signals of populations experiencing genetic erosion to inform conservation actions to mitigate loss of genetic diversity (*24*, *25*). We have previously shown that individual heterozygosity in these *Cy. semiargus* populations is positively related to grassland area on the landscape level (5-20km from the local patch, *44*). Combined with the findings from the current study, these results show that preservation of current grassland area and configuration will be insufficient for maintaining genetic diversity in *Cy. semiargus* populations in southern Sweden. Rather, restoration of suitable habitat at a scale that facilitates gene flow between populations is vital to prevent further genetic erosion in this species, and potentially other insect taxa that dependent on semi-natural grassland habitat.

In conclusion, we document the early stages of genomic erosion in a specialist grassland butterfly in southern Sweden. While census size-based estimates of demographic declines largely inform current conservation status of these species, the isolation and inbreeding of *Cyaniris semiargus* highlights that even formerly common grassland insects may now carry a substantial genetic extinction debt, forming the basis for an extinction debt at the community level (*25*). Our findings suggest that prompt restoration of grassland connectivity, especially in agricultural regions, is needed to restore gene flow and maintain genetic diversity in specialist grassland butterflies, reducing their risk of extirpation and extinction. This work emphasizes the urgency of semi-natural grassland restoration efforts to meet the goal of maintaining genetic diversity under Goal A of the Kunming-Montreal Global Biodiversity Framework (*20*). Additionally, the inbreeding we observe in a formerly abundant grassland butterfly points to the importance of considering population connectivity during the restoration process if restoration is to successfully halt the erosion of genetic diversity. Broadly, our findings underscore that genetic monitoring is a valuable tool in conservation assessment of insects, as early detection of genomic erosion can highlight populations that may have stabilized but still carry a genetic extinction debt, enabling timely action.

## Materials and Methods

### Study System and Sampling

To assess changes in genetic diversity, differentiation, inbreeding, and deleterious mutation burden, we compared modern and historical genomes from three focal blue butterfly species (Polyommatinae), *Polyommatus icarus*, *Plebejus argus*, and *Cyaniris semiargus* (Figure 1A). These species vary in habitat specialization, being grassland generalist, heathland specialist, and grassland specialist, respectively. We used previously generated population-level genetic data (*44*) for the modern genomes. For the historical genomes, we selected pinned specimens from the entomological collections of the Biological Museum, Lund University (*43*) sampled between 1917-1956 in roughly the same geographic area as the modern samples. For each historical locality, we selected 4-11 specimens sampled in the same year to estimate population-level metrics. The historical and modern population-level pairs were sampled in four broad geographic areas: I) Western and II) Eastern Skåne, III) Småland, and IV) Öland (Figure 1B), with historical localities ranging approximately 12-40km from their modern counterparts. This resulted in three historical-modern population pairs for *Po. icarus* and two for *Pl. argus*. For *Cy. semiargus*, we included three pairs and an additional comparison of a modern Southeastern Skåne population paired with the historical Eastern Skåne population. For *Pl. argus*, only one historical specimen was available from Western and Eastern Skåne, respectively, and these were compared to the closest modern populations in individual-level analyses only.

To more broadly assess the variation in rates of genetic decline in Polyommatinae of differing conservation statuses, we sampled historical and modern individuals of five additional species: *Agriades optilete, Aricia agestis, Cupido minimus, Plebejus argyrognomon* and *Scolitantides orion.* Modern specimens of *Ag. optilete, Ar. agestis,* and *Cu. minimus* were collected across the same geographic regions as the focal species in 2021 (Table S1). A pinned specimen used as the modern sample of *Sc. orion* was collected in 2016 in Södermanland and donated to this study by Håkan Elmqvist. Legs from a pinned speciemen of *Pl. argyrognomon*, collected in 2003 in Western Skåne, were donated by Göran Engqvist and used as the modern sample for this species. Historical samples for these five species were selected to be proximal to the modern specimen sampling locality with collection dates ranging from 1939-1967, all provided as pinned specimens from the entomological collections of the Biological Museum, Lund University (*43*).

### DNA Extraction and Sequencing

DNA from fresh specimens was extracted and sequenced following Nolen et al (*44*). We extracted DNA and prepared sequencing libraries from pinned museum specimens in a facility dedicated to historical DNA processing at Lund University, following the protocol described by Twort et al (*79*). Libraries prepared for pinned specimens were pooled into two equimolar pools containing 37-38 libraries each, sequencing each pool on a single NovaSeq 6000 S4 lane. DNA was extracted from the abdomens or legs of male specimens without homogenization to preserve anatomical structures.

### Sequence Processing

Sequence processing and analyses were largely performed using Snakemake v8.25.5 (*80*), incorporating and extending the PopGLen v0.4.1 (*81*) workflow. We trimmed raw sequencing reads using fastp v0.23.4 (*82*) with default settings, additionally collapsing overlapping read pairs in historical samples, discarding those with less than 30bp overlap. As historical DNA is expected to be fragmented, mapping can be biased towards reads containing the reference allele. We investigated two potential alignment options to reduce this impact: (I) BWA aln (*83*) with relaxed parameters, as is used in most historical DNA studies, and (II) alignment to a variation graph incorporating known alternative variants from the modern samples, which has been shown to reduce reference biases when mapping short fragments (*84*). As we found that mapping to the variation graph reduced the amount of reference bias in the historical samples (Figure S2&3), we performed all analyses with both modern and historical samples aligned to the species-specific variation graphs. Detailed methods of the processing for this comparison and the final alignment method are available in the Supplementary Methods, with a brief outline of the final method here.

To construct variation graphs for each species, we first aligned the trimmed, paired-end modern reads to the respective reference genome (Table S12) (*85–87*) using BWA mem v0.7.17 (*88*), removed duplicates using Picard v2.27 (*89*), and realigned around indels using GATK (*90*). We called genotypes in these samples using the BCFtools multi-allelic caller v1.21 (*91*) and constructed a variation graph from these variants and the linear reference genome using vg v1.63.1 (*92*). To produce the final alignments, we mapped modern and historical samples to the variant graph using vg with default settings, additionally setting -k 15 for historical samples as recommended by Martiniano et al. (*84*). We removed duplicates with Picard and dedup v0.12.8 (*93*) for modern and historical samples, respectively, and clipped overlapping read pairs using BamUtil v1.0.15 (*94*). We estimated post-mortem DNA damage in historical samples using DamageProfiler v1.1 (*95*).

### Genotype calling and filtering

To account for variation in sequencing depth, we subsampled all individuals to a mean depth of 6x using Samtools v1.20 (*91*). We excluded individuals with a mean coverage of less than 5x from genotype calling. We implemented a sites-based filtering scheme (see Supplementary Methods) and called genotypes for each species jointly across samples using the BCFtools multi-allelic caller at sites passing the species filter sets (Table S13), including reads with minimum mapping and base qualities of 30 and 20, respectively. We filtered to biallelic and invariant positions, excluding those within 5bp of indels or a variant quality less than 30. We set genotypes with a depth less than 6 and heterozygous genotypes with one allele supported by less than 20% of the reads to missing. We removed transition positions across all samples and filtered out positions with more than 40% missing data in either the historical or modern sample subsets.

### Heterozygosity, nucleotide diversity, and genetic differentiation

We estimated individual genome-wide heterozygosity for all samples from the called genotypes by dividing the count of heterozygous calls by the total count of called genotypes, including invariant sites. We estimated nucleotide diversity using Pixy v2.0.0.beta2 (*96*), which utilizes a sample-size weighted calculation. Nucleotide diversity estimates in these populations are robust to low (N = 4) sample sizes (*44*), so we included estimates for all populations with multiple individuals. We estimated this summary statistic in 50kb non-overlapping windows for populations grouped by sampling locality, estimating the genome-wide mean across windows, and confidence intervals from 10,000 bootstraps, resampling windows with replacement. We estimated genetic differentiation using pairwise population *F*_ST_ with Pixy on the same windows with the Hudson estimator (*97*), which is robust to small and variable sample sizes (*98*). We estimated mean *F*_ST_ and confidence intervals with the same approach as nucleotide diversity.

### Relatedness

As analyses of population structure can be sensitive to the presence of close relatives in the dataset, we estimated pairwise relatedness between all samples within each species dataset. We utilized the IBSrelate SFS-based method (*99*), as implemented in NGSrelate (*100*), to estimate R0, R1, and KING-robust kinship coefficients (*101*) for all pairs of individuals in the dataset. For input, we generated a Beagle file for each focal species dataset, using all sampled individuals, but subsampling individuals with higher coverage to 8X to reduce the influence of the large variation in coverage in our modern samples. We estimated genotype likelihoods in ANGSD v0.940 (*102*) using the GATK model from reads with a mapping quality >= 30, base quality >= 20, at positions with a minimum depth of three that passed species-specific site filters. We called SNPs with a p-value < 1e-6, a minor allele frequency >= 0.05, and with data in at least 90% of the individuals.

### Population Structure

We explored population structure using principal component analysis (PCA) as implemented in PCAngsd v1.10 (*103*). To obtain independent sites for population structure analyses, we estimated linkage disequilibrium in the Beagle file from the relatedness analyses using ngsLD v1.2.0 (*104*) and pruned SNPs using prune_graph (https://github.com/fgvieira/prune_graph; v0.3.2), assuming independence of SNPs >50kb apart and with a coefficient of correlation (r^2^) < 0.1. We removed one individual from each pair of third-degree or closer relatives in the dataset (expected KING-kinship coefficient >= 0.0625; Table S14) before running the PCA.

We formally tested for population structure using admixture analyses for 1-8 ancestral clusters (K) using the pruned, relative removed Beagle file in NGSadmix v33 (*105*). We performed at least 20 replicate runs for each value of K, up to 100 total runs, stopping if the three highest likelihood replicates converged within 2 log-likelihood units of each other. The ‘best-fit’ value for K was then determined to be the highest K for which replicates met this convergence criteria, following Pečnerová et al (*106*). We further assessed the model fit for each value of K using evalAdmix v0.961 (*107*).

### Runs of Homozygosity

We assessed rates of recent and background inbreeding by estimating the inbreeding coefficient *F_RoH_*, the proportion of the autosomal genome existing in runs of homozygosity over a specified length, incorporating runs of homozygosity >100kb in our estimates of *F_RoH_*. Per individual runs of homozygosity were estimated using BCFtools/RoH v1.19 (*108*). We set a constant species-specific recombination rate, assuming 50 centimorgans per autosomal chromosome as in Mackintosh et al (*33*) (Table S12), and ran the analysis on genotype calls at variable sites, setting the PL of unseen genotypes set to 30 as suggested by the documentation. To ensure *F_RoH_* estimates were inferred equivalently in species with population and individual level sampling, we did not estimate allele frequencies, instead using a default minor allele frequency of 0.4 as suggested in the documentation and ignored homozygous reference positions, as including these can inflate estimates of *F_RoH_* based on our testing (Figure S11). We calculated *F_RoH_* for each individual by dividing the cumulative length of runs of homozygosity by the total length of the autosomes.

### Recent demography and coalescent simulations of heterozygosity loss

To infer timing and magnitude of demographic declines that may contribute to reduced genetic diversity, we inferred changes in effective population size over the past 100 years from the modern samples using the linkage disequilibrium-based approach implemented in GONE (*109*). We inferred these histories in modern samples by sampling locality, as this method performs best when input samples do not have additional genetic substructure (*110*). We performed this analysis on the called modern variants used to construct the variation graphs, which included transitions. As we observed a frequent sudden drop in the most recent four generations in most of the GONE analyses, we visualized trajectories both including and excluding these generations (Figure S5).

To assess if our empirical estimates of heterozygosity decline and accumulation of runs of homozygosity fell within expected ranges for the characterized demographic changes, we performed coalescent simulations using Slendr (*111*) with the msprime (*112*) backend. For each modern locality with a paired historical locality, we constructed demographic models that followed size changes inferred for the most recent 100 generations in the GONE analyses. As most sampled populations showed a roughly constant historical size by this time point (Figure S5), we set the ancestral size to that of generation 100. For localities where we inferred a constant history of very large size (>100,000), we modeled a simple constant size model with *N* = 100,000. Due to the sharp drop in the four most recent generations, we set these generations in the model to the estimate of generation five. For each sampled locality, we ran 100 replicate simulations of a single, 10 Mb chromosome with a recombination rate corresponding to 50cM (*33*) and a mutation rate of 2.9e-8 (*113*). At time points corresponding to the collection of the historical samples for a given locality and the most recent generation, we sampled 100 individuals from each simulation and calculated mean heterozygosity, *F_RoH_*, and heterozygosity outside of *RoH* to infer a range of expected values to compare our empirical estimates to.

### Deleterious mutation burden

As proxies for genetic load, we estimated the burden of deleterious mutations per individual using the count of derived alleles with putatively deleterious consequences. We inferred these consequences using two complementary methods, one based on gene annotations to infer functional effects (*114*) and the other identifying positions with high sequence conservation (*115*). As variants were called across the combined historical and modern sample set for each species, alleles that are fixed within one time category, but variable or fixed for the opposing allele in the other, are included.

To identify which alleles were likely ancestral, we co-inferred ancestral states and sequence conservation scores using the GenErode pipeline v0.6.2 (*116*), which estimates Genomic Evolutionary Rate Profiling (GERP) scores (*68*) from a set of outgroup species assemblies. We utilized 24 Lycaenidae chromosome-level genomes from distinct genera (Table S15) (*85–87*, *117–120*) as outgroups, modifying the default fragment size for GenErode’s alignment step from 35 to 100bp, and using divergence times estimated with TimeTree 5 (*121*). We then subset the variable sites to include only those with GERP > 0 and where the ancestral allele matched the reference to avoid underestimating derived counts in historical samples.

We identified deleterious mutations based on functional effects using the genome annotations provided by the Darwin Tree of Life Project (https://projects.ensembl.org/darwin-tree-of-life/). We assigned impacts to each variant using Ensembl’s Variant Effect Predictor (VEP) v112.0 (*114*). VEP categorizes variants into four classes: ‘High’, ‘Moderate’, ‘Low’, or ‘Modifier’, which largely correspond to loss of function, missense, synonymous, and intergenic variants. We forced VEP to pick one effect per variant, using the default selection order (*114*). To identify conserved positions, we selected the top 1% of GERP scores as the most conserved positions using the get_gerp_score_percentile.py script included with GenErode. We inferred deleterious burden at positions inferred as loss of function (‘High’ impact), missense (‘Moderate’ impact), or conserved based on GERP scores, using the derived allele count at these positions. Total deleterious burden is the total derived allele count of a variant category per 1000 called genotypes.

Homozygous deleterious burden is the homozygous derived allele count per 1000 called genotypes. We additionally estimated these counts at synonymous (‘low’ impact) and intergenic positions to assess biases in derived counts across sample types.

### Categorizing rates of genetic decline

To assess the changes in genetic diversity in a framework developed for conservation focused genetic monitoring, we used the indicator thresholds suggested by Andersson et al (*21*), which are based upon indicators applied for national use in Sweden (*58*). Using these thresholds, genetic decline within populations is measured as the amount of genetic diversity retained over a period of 100 years, assuming a constant rate of change. Three threat categories are suggested: retention >=95% is considered ‘Acceptable’, 75-94% as ‘Warning’, and <75% as ‘Alarm’. We estimated the predicted retention of genetic diversity over 100 years for each population using individual heterozygosity and nucleotide diversity.

Genetic diversity retention per year was estimated using the following equation:

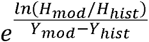

where *H* is the estimate of genetic diversity for a given sample/population and Y is the sampling year for that sample/population. We estimated this metric for nucleotide diversity in the focal species using population mean estimates. For individual heterozygosity, we estimated this metric using population means in the focal species and using all pairwise comparisons of historical and modern individuals in both focal and supplemental species. For the effective population size indicator, we utilized the estimate of effective population size at generation five in the GONE analyses.

## Acknowledgements

We would like to thank Melina Eberhagen, Amanda Görnerup, Kajsa Svensson, Aske Brydegaard Runemark, and Mikkel Brydegaard for assistance with collecting butterflies, Håkan Elmqvist for providing pinned *Sc. orion* specimen, Göran Engqvist for providing tissue from the pinned specimen of *Pl. argyrognomon*, Emma Kärrnäs and Isolde van Riemsdijk for assistance with DNA extractions, Jadranka Rota for assistance with the entomological collections of the Biological Museum, Lund University (https://doi.org/10.15468/dahk2a), Verena Kutschera and the Swedish Bioinformatics Advisory Program for advice regarding the bioinformatic analyses, Anna Penna and Danielle Clake for comments on the manuscript, and Isolde van Riemsdijk, Rachel Steward, Kalle Tunström, Simon Jacobsen Ellerstrand, Colin Olito, Magne Friberg, Mikkel Skovrind, and Carolina Pacheco for helpful discussions regarding the study species, methods and interpretations. We would like to thank Project Psyche and the Darwin Tree of Life project for generating and providing the reference genomes that made these analyses possible.

We acknowledge support from the National Genomics Infrastructure in Genomics Production Stockholm funded by Science for Life Laboratory, the Knut and Alice Wallenberg Foundation and the Swedish Research Council. Computational resources were provided by the National Academic Infrastructure for Supercomputing in Sweden (NAISS) and the Swedish National Infrastructure for Computing (SNIC) at UPPMAX partially funded by the Swedish Research Council through grant agreements no. 2022-06725 and no. 2018-05973. This work was supported by grants from the strategic research area “Biodiversity and Ecosystem services in a Changing Climate” (BECC), the Foundation Oscar och Lili Lamms Minne, and the Swedish Research Council for Sustainable Development (grant 2021-01163) to AR and the Jörgen Lindström’s Foundation to ZJN.

## Author Contributions

**Z.J.N.** - Conceptualization, Methodology, Formal analysis, Investigation (Lead), Data curation, Writing – original draft, Visualization, Funding acquisition. **P.J.** - Investigation (Supporting), Writing – review & editing. **A.S.T.L.** - Investigation (Supporting), Writing - review & editing. **N.W.** - Methodology, Writing – review & editing, Resources, Supervision. **A.R.** - Conceptualization, Methodology, Writing – original draft, Supervision, Funding acquisition.

## Data Availability Statement

The genetic data that support these findings will be made available upon publication at NCBI in BioProjects PRJNA1068054 (focal species, modern) & PRJNA1168917 (focal species, historical; additional species). Bioinformatic workflow configurations, extensions, and resources specific to this study are available at https://github.com/zjnolen/polyommatini-temporal-genomics and archived at https://doi.org/10.5281/zenodo.13902816.

