## Supplementary Materials for "Species-specific loss of genetic diversity and inbreeding following agricultural intensification"

### Supplementary Material

### Supplementary Methods

**Sites based filtering scheme.** We utilized a sites-based filtering scheme similar to that implemented by Pečnerová et al (*106*) which intersects multiple independent sets of filtered sites to create a global list of filtered sites. We performed this filtering for each species separately. We excluded sex-linked chromosomes, mitochondria, and scaffolds less than 1 Mb in length. We excluded repeats based on the softmasked reference genomes provided by Ensembl and the Darwin Tree of Life Project (*122*; https://projects.ensembl.org/darwin-tree-of-life/). For species with no softmasked genome available, we identified repeats using RepeatModeler (*123*) and RepeatMasker (*124*). We estimated positions in the lower and upper 1% of global sequencing depth distributions separately across all samples, historical samples, and modern samples, excluding sites that appeared in any of the three sets. The number and proportion of sites passing each of these filters, as well as passing the final intersected filter set, are reported for each species in Table S13.

**Investigating different alignment options to reduce reference bias**. Higher reference bias is expected in historical DNA as shorter fragments lead to those containing reference alleles to be favored over those with alternates in the alignment process (*125*). These concerns are especially relevant to studies which aim to compare estimates between degraded historical samples and high-quality contemporary ones, where different average sequence length can lead to differences in reference bias that drive the different signals in the historical and contemporary samples. We investigated the amount of reference bias and the impact on downstream analyses under different alignment strategies. The first strategy uses bwa aln (*83*) with relaxed edit distance and gap open parameters (*126*, *127*), the most common technique in historical DNA studies to improve alternative allele mapping (hereafter referred to as 'bwa aln'). The second augments the reference genome into a variation graph with information on known alternative variants identified in modern individuals, improving the ability of shorter fragments to map alternative alleles (*84*) (hereafter referred to as 'vg').

To generate the dataset for alignment with bwa aln, we aligned each sample to the corresponding reference genome (*85*–*87*) using the steps in the PopGLen pipeline v0.4.1. This first trims adapters and poly-G tails using fastp v0.23.4 (*82*), keeping reads as paired in modern samples and collapsing overlapping reads in historical samples with a 30bp overlap, discarding reads < 30 bp from both after trimming and collapsing. We then mapped modern samples to the references with bwa mem v0.7.18 (*88*) using default settings and collapsed reads from historical samples with bwa aln v0.7.18, disabling seeding (-l 16500) decreasing the fraction of missing alignments (-n) from the default of 0.04 to 0.01 and increasing the number of gap opens (-o) form 1 to 2. We additionally disabled seeding (-k 16500) due to the expectation that the first bases of reads may have DNA damage. After alignment, we removed duplicates with Picard v3.2.0 (*89*) in modern samples, setting the optical duplicate pixel distance to 2500, and using dedup v0.12.8 (*93*) in historical samples. We realigned around indels in both modern and historical samples using the GATK IndelRealigner v3.8 (*90*). As ANGSD (*102*) counts overlapping paired reads twice (*128*), we clipped overlapping reads in the modern samples with BamUtil v1.0.15 (*94*).

To generate the dataset for alignment with vg, we joint called genotypes across the modern samples of each species aligned with bwa mem using the BCFtools multi-allelic caller v1.21 (*91*). We called genotypes with a mapping and base quality filter of 30, excluded positions in the upper percentile of the global depth distribution, and grouped samples by sampling locality for the built-in Hardy Weinberg equilibrium assumption. We filtered out genotypes with a genotype quality < 30 or depth < 6 and filtered positions to biallelic SNPs, removing those within 5 bp of indels, a quality < 30, a minor allele frequency < 0.05, or missing data in > 40% of the samples after genotype filtering. We then constructed a variation graph for each species from the linear reference genome and the filtered variants VCF using vg v1.63.1 (*92*). To generate the final modern sample alignments, we re-aligned the trimmed modern reads to variation graph using vg with default settings, re-ordered scaffolds to match the reference and removed duplicates with Picard and clipped overlapping reads with BamUtil. To generate the final historical sample alignments, we aligned the collapsed reads using vg, setting a minimum mem length of 15 (-k 15) and set the band width for long reads to a size greater than the largest fragments included (-w 300) as recommended by Martiniano et al. (*84*).

To estimate reference bias, we utilized the identity by state distance to the reference averaged across sites. We made this estimate using single read sampling at each position as well as from called genotypes. We estimated single read sampling IBS distance per sample using ANGSD using the settings implemented in PopGLen to randomly sample a single read (-doIBS 1) at each position passing the filtered sites set, removing transitions (-rmTrans 1), including only sites with at least three reads (-setMinDepth 3) and only using reads with a minimum base quality of 20 and mapping quality of 30. To estimate the identity by state distance for called genotypes, we called and filtered genotypes from the bwa aln and vg alignments separately, following the methods described in the main manuscript, to produce comparable BCFs. Identity by state similarity to the reference was estimated per individual as the count of called reference alleles divided by 2 times the count of called genotypes.

We found that using vg reduced reference bias for both modern and historical samples compared to bwa aln, though this effect was larger for historical samples (Figure S2). This effect was species-specific, with the largest improvement in *Po. icarus*, followed by *Pl. argus*, then by *Cy. semiargus*. This pattern follows patterns of heterozygosity in these three species (*44*), reflecting that disparities in reference bias between historical and modern samples are likely more pronounced in species with higher heterozygosity. This has implications for other population genomic studies working with historical insect genomes, as higher levels of heterozygosity in insects compared to vertebrates (*36*) may mean that standard alignment methods do not sufficiently reduce reference bias in these organisms to produce reliable downstream results.

To understand better how the mismatch rates of the different aligners changed with fragment size, we plotted average mismatch rates per fragment size for a *Po. icarus* historical sample using AMBER (*125*), including only fragments with mapping qualities >= 30. We did this for alignments with bwa aln and vg with the settings described above. We found that under all three tested bwa aln settings, mismatch rates were generally lower than with vg, and that an even pattern of mismatches across the majority of fragment sizes was not achieved even with the most permissive bwa aln settings we tested (i.e. -n 0.01 -o 2; Figure S3). While the pattern of mismatch rates is more even across fragment sizes for vg, we still see a reduced mismatch rate for smaller fragments of sizes that make up a considerable portion of the fragment distribution (Figure S3), which likely explains why some reference bias in the historical samples still persists when using this aligner.

Additionally, to illustrate the impact of reference bias on downstream analyses, we performed principal component analyses (PCAs) for each of the three species when aligned with bwa aln and aligned with vg (Figure S4). We found that samples clustered primarily by sampling time period when aligned with bwa aln, likely due to reference biases shifting allele frequencies in the historical samples towards the reference allele. When aligned with vg, historical samples clustered instead with modern counterparts sampled in nearby localities, as is expected biologically. In *Po. icarus*, where all population genetic structure is captured in PC1, we still observe some clustering by sampling time period along PC2 when aligned with vg, potentially reflecting the greater residual reference bias in this species.

Given the more even mismatch rate across fragment sizes and generally lower disparity of mismatch rates when using vg compared to bwa aln, we chose to perform analyses with alignments made with vg for the main manuscript. As filtered genotype calls further showed reductions in reference bias compared to randomly sampled reads, we additionally chose to utilize genotype calls for the majority of analyses rather than genotype likelihoods.

### Supplementary Figures and Tables

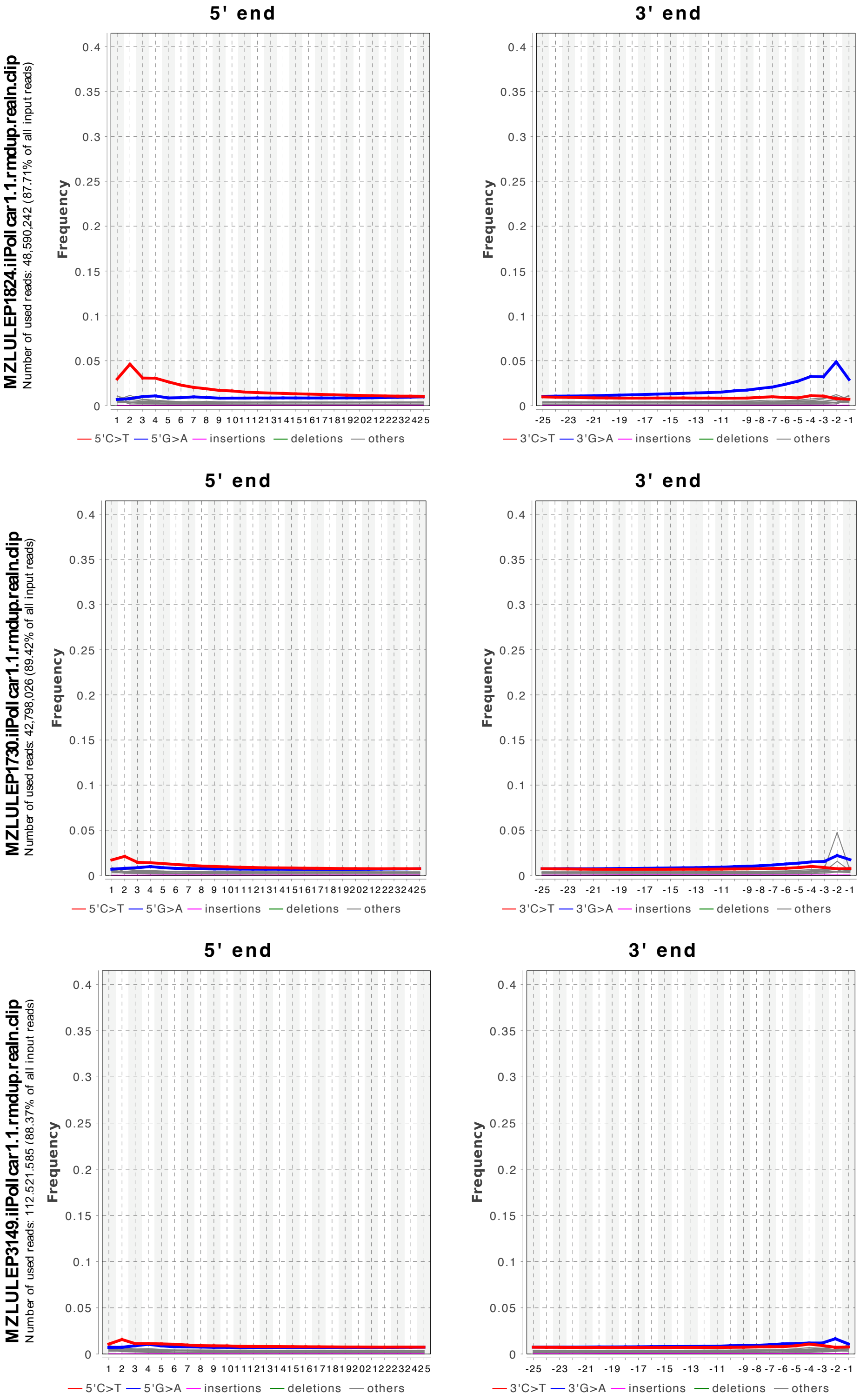


**Figure S1. Examples of DNA damage profiles for historical samples.** Damage profiles produced by DamageProfiler (*95*) for the samples with the greatest (upper), median (middle), and least (lower) DNA damage in the focal species dataset are shown. In all analyses, we accounted for these misincorporations by removing transition variants.

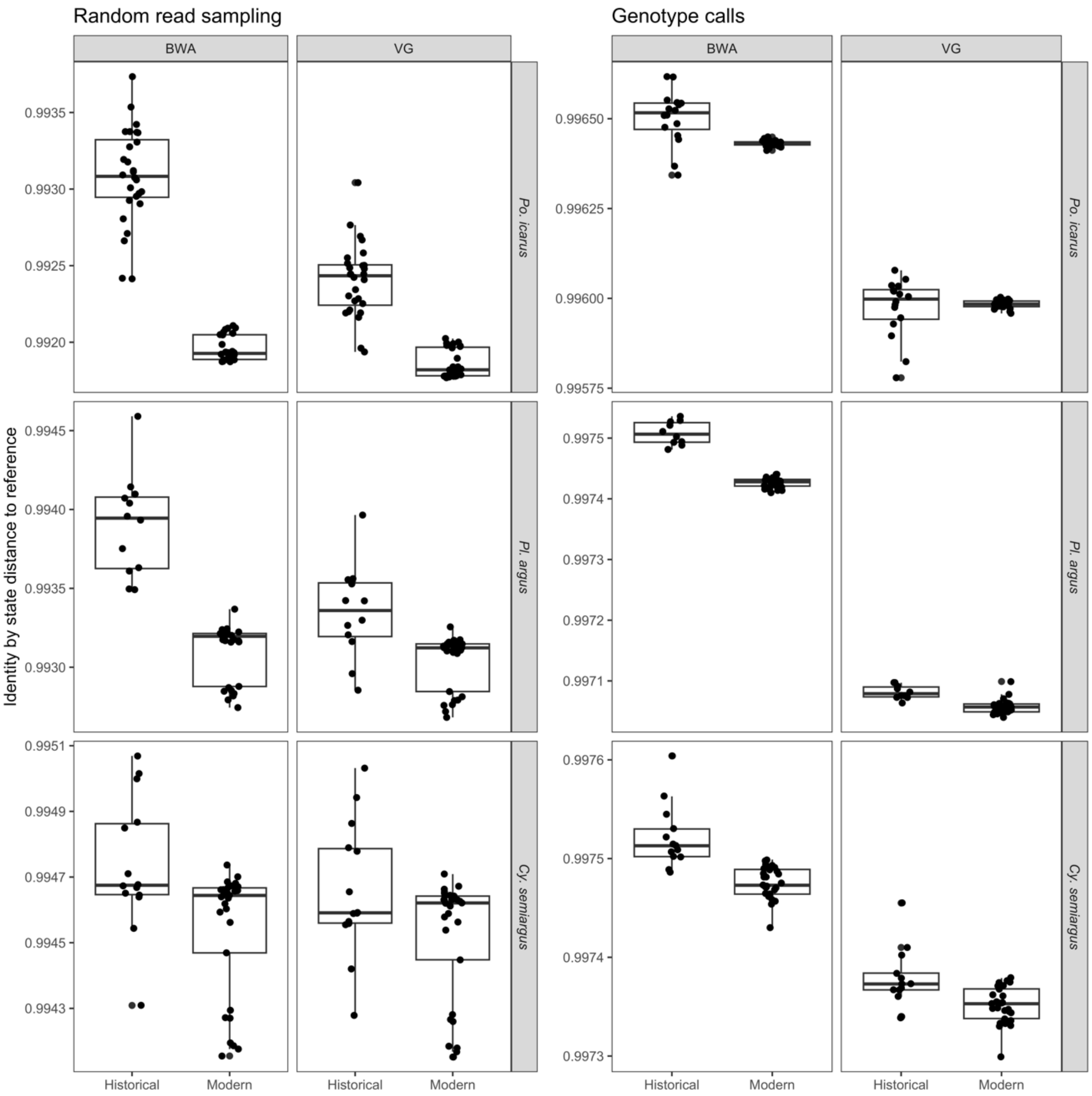


**Figure S2. Examining effects of alignment method on reference bias.** As historical samples are expected to experience reference biases due to shorter fragment size compared to modern samples, we assessed the impact of alignment method on reference bias, comparing alignment to a linear reference with bwa with relaxed parameters to alignment to a variation graph incorporating known variants from modern samples with vg. Reference bias of the two alignment methods is shown here for historical and modern samples of the three focal species, estimated both by sampling a random read at all positions with >= 3x depth (left panels) and from called genotypes (right panels). We find that the variation graph reduced the disparity in reference bias between historical and modern samples, selecting this alignment method to move forward with for downstream analyses.

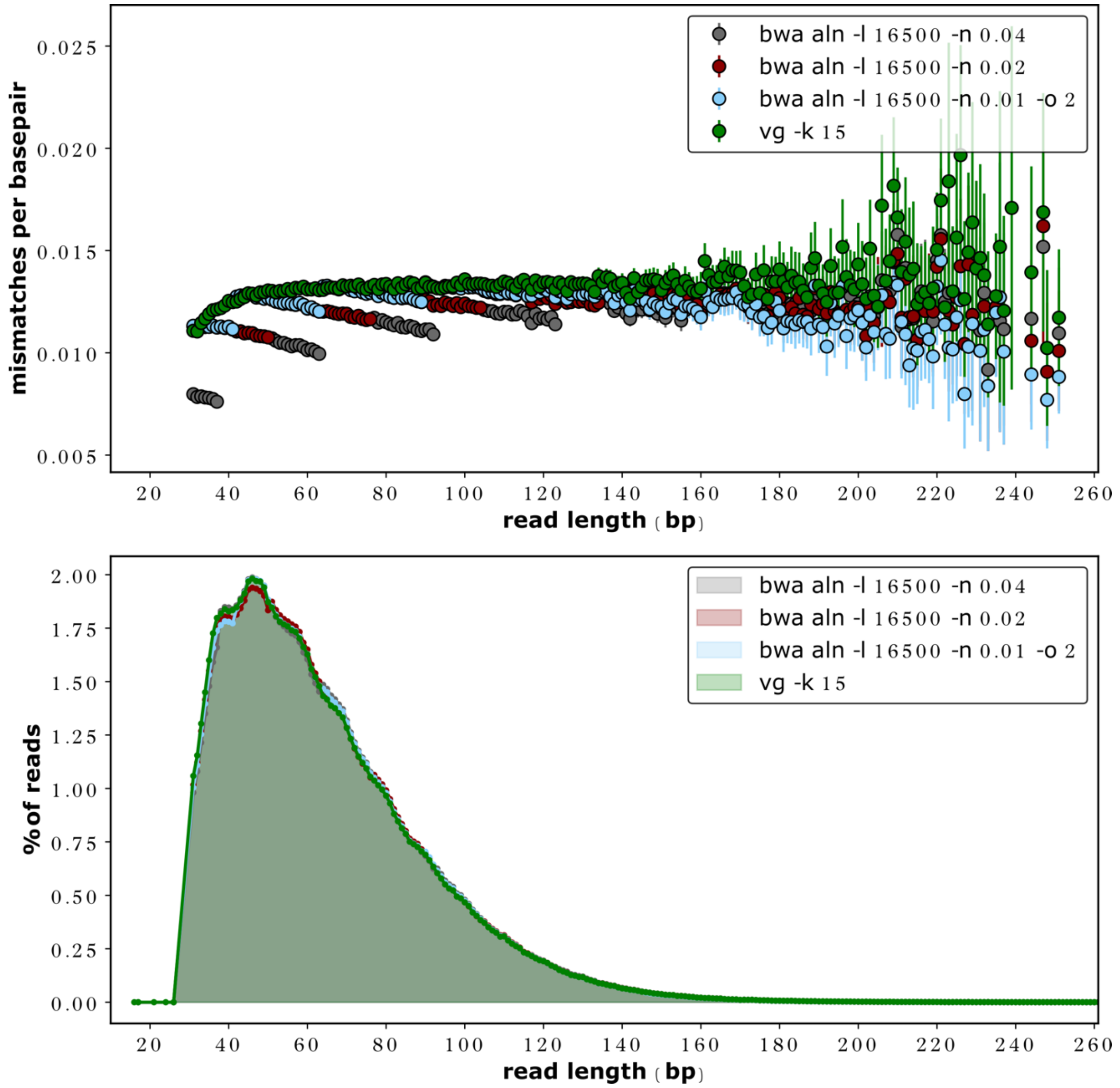


**Figure S3. Mismatches per base pair by fragment size under various alignment settings.** (Upper) Mismatch rates per base pair for different fragment sizes when aligned with different tools (vg vs. bwa aln) and settings in the case of bwa aln. Only reads with mapping quality >= 30 are included. (Lower) Distribution of mapped read lengths, reads below 30bp have been discarded. Figures created using AMBER (*125*).

**Historical samples aligned to linear reference with bwa aln**

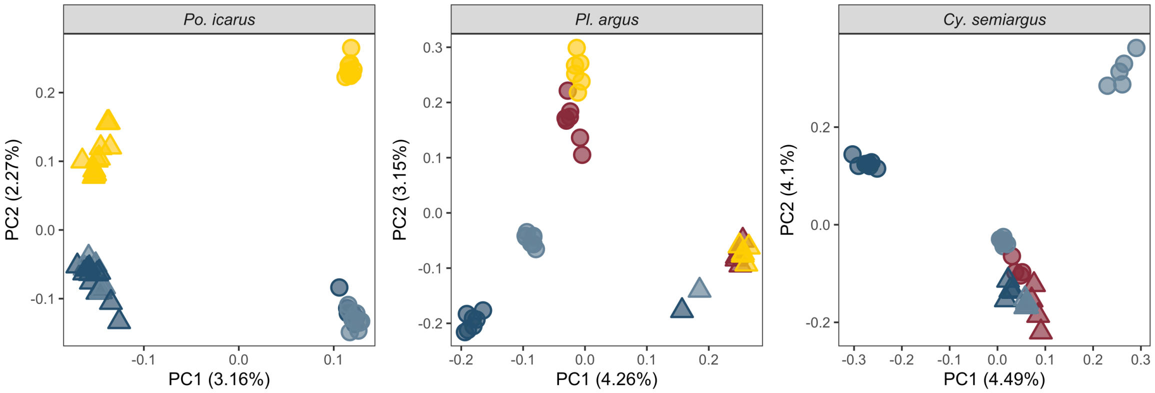


**Historical samples aligned to variation graph with vg (subset from Figure 2)**

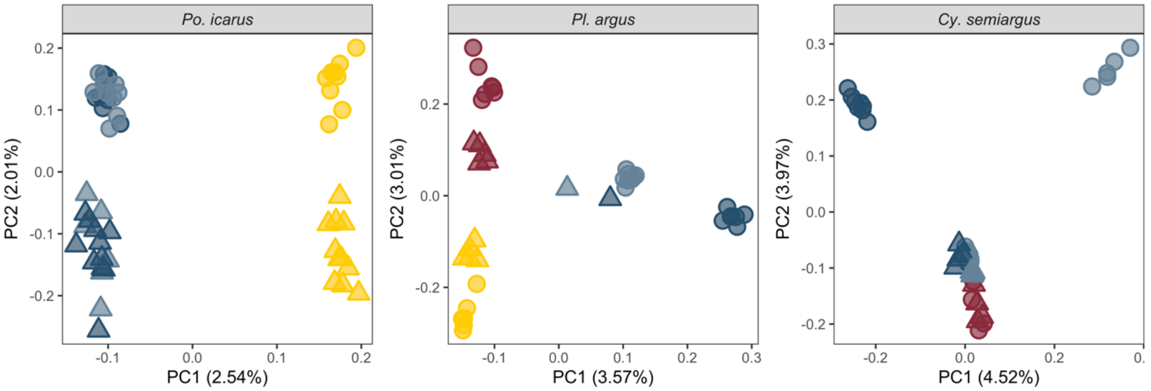


**Figure S4. Comparison of downstream population structure analyses under different alignment methods.** Principal component analyses (PCA) for the three focal species when historical samples are aligned to the linear reference genome with bwa aln (upper) and to the variation graph reference incorporating modern sample variants with vg (lower). Each circle represents a modern and each triangle a historical individual, with colors corresponding to the sampling locations in Figure 1B. Percentages on each axis refer to the proportion of total variance explained by the component. When aligned to a linear reference using bwa aln with relaxed parameters that reduce reference bias (-l 16500 -n 0.01 -o 2; Figure S3), we find that bias is still substantial enough that samples cluster primarily by sample type, as historical allele frequencies are artificially shifted towards the reference allele (upper panel; historical and modern samples separate on PC1 for *Po. icarus* and *Pl. argus* and on PC2 to some extent in *Cy. semiargus*). Utilizing a variation graph that incorporates common variants from the modern samples results in samples clustering by sampling locality (lower panel), as is expected, enabling more accurate comparison of the two different sample types. Clustering by sample type is still visible on PC2 for *Po. icarus* and may reflect the residual reference bias being greater in this species (Figure S2).

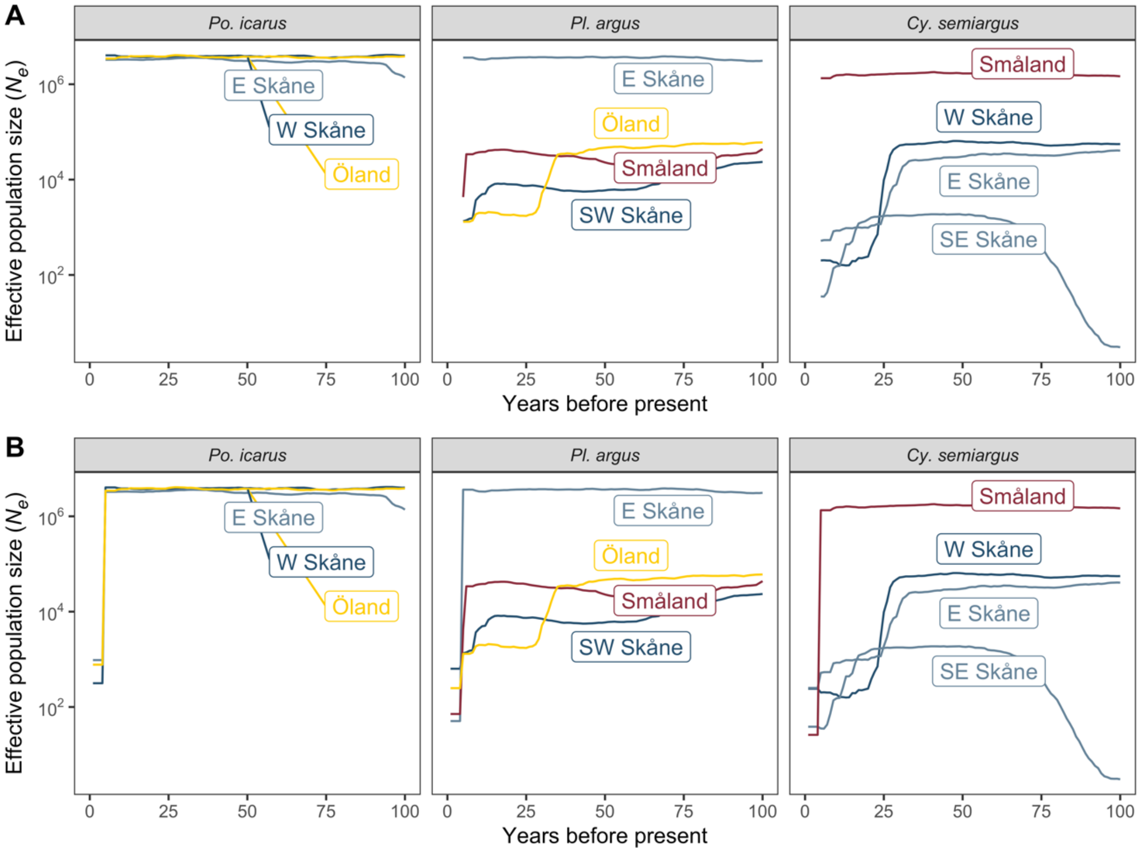


**Figure S5. Effective population size trajectories for the past century estimated with GONE (*109*).** For each sampled modern population, we inferred effective population size changes over the past century using the linkage disequilibrium-based method implemented in GONE. Trajectories are presented here going backward in time and are presented on a log scale. Lines are colored and labeled by population for each species. As the most recent four generations often had a sharp decrease in effective population size, we present the trajectories here excluding (A) and including (B) these generations. As the change is almost always in the downward direction, we utilized the estimates at generation five for estimates of contemporary effective population size, as this provides a more conservative estimate. For parameterizing the coalescent simulations, we utilized the trajectories with the most recent four generations fixed to the effective population size of generation five (i.e. as depicted in A). Low historical (>50 years before present) effective population sizes in SE Skåne *Cy. semiargus* may be an artifact of the exceptionally low recent effective population size, as dramatic recent changes in $N_e$ may erase signals of older demographic events (*109*).

### *Po. icarus*

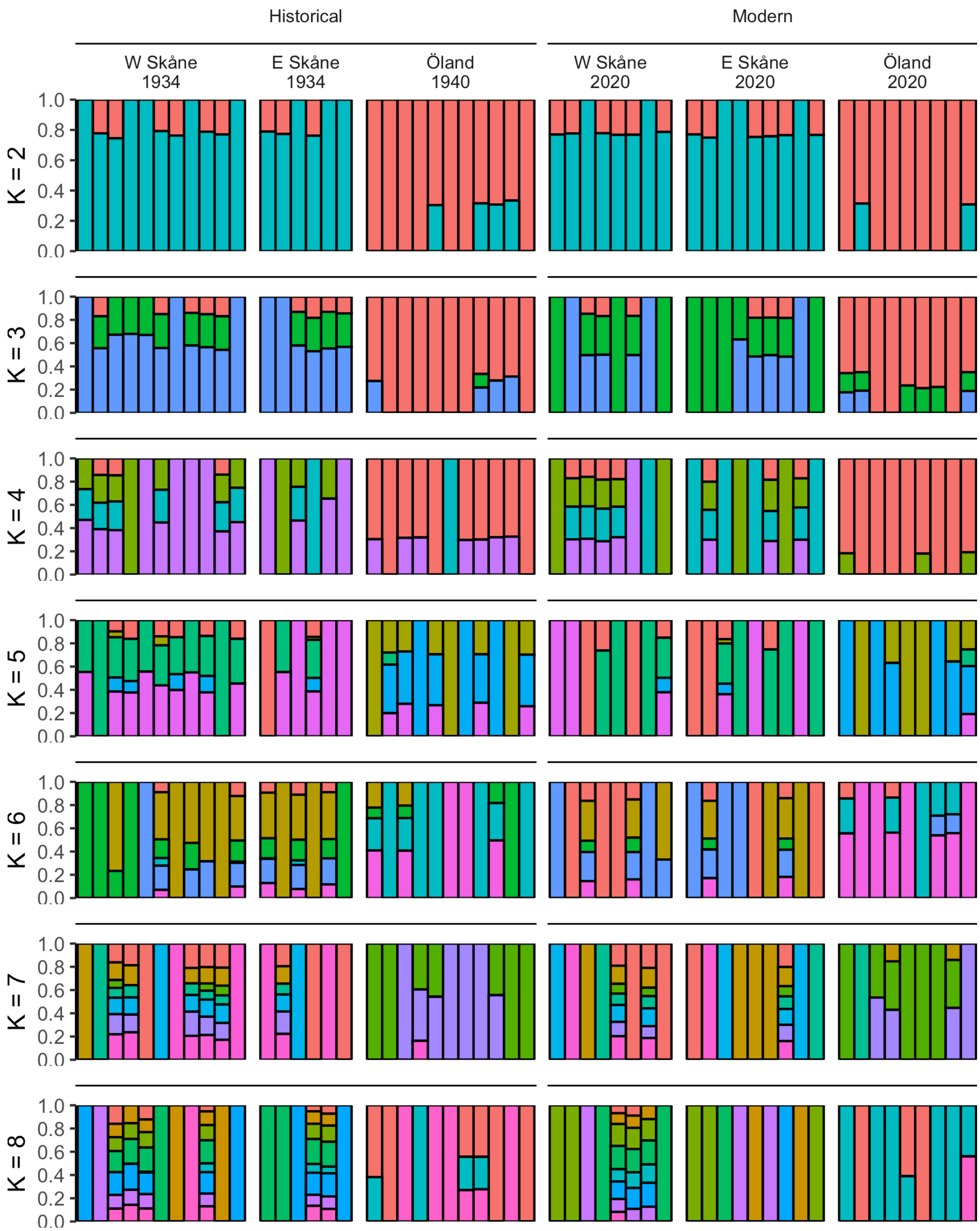


1102

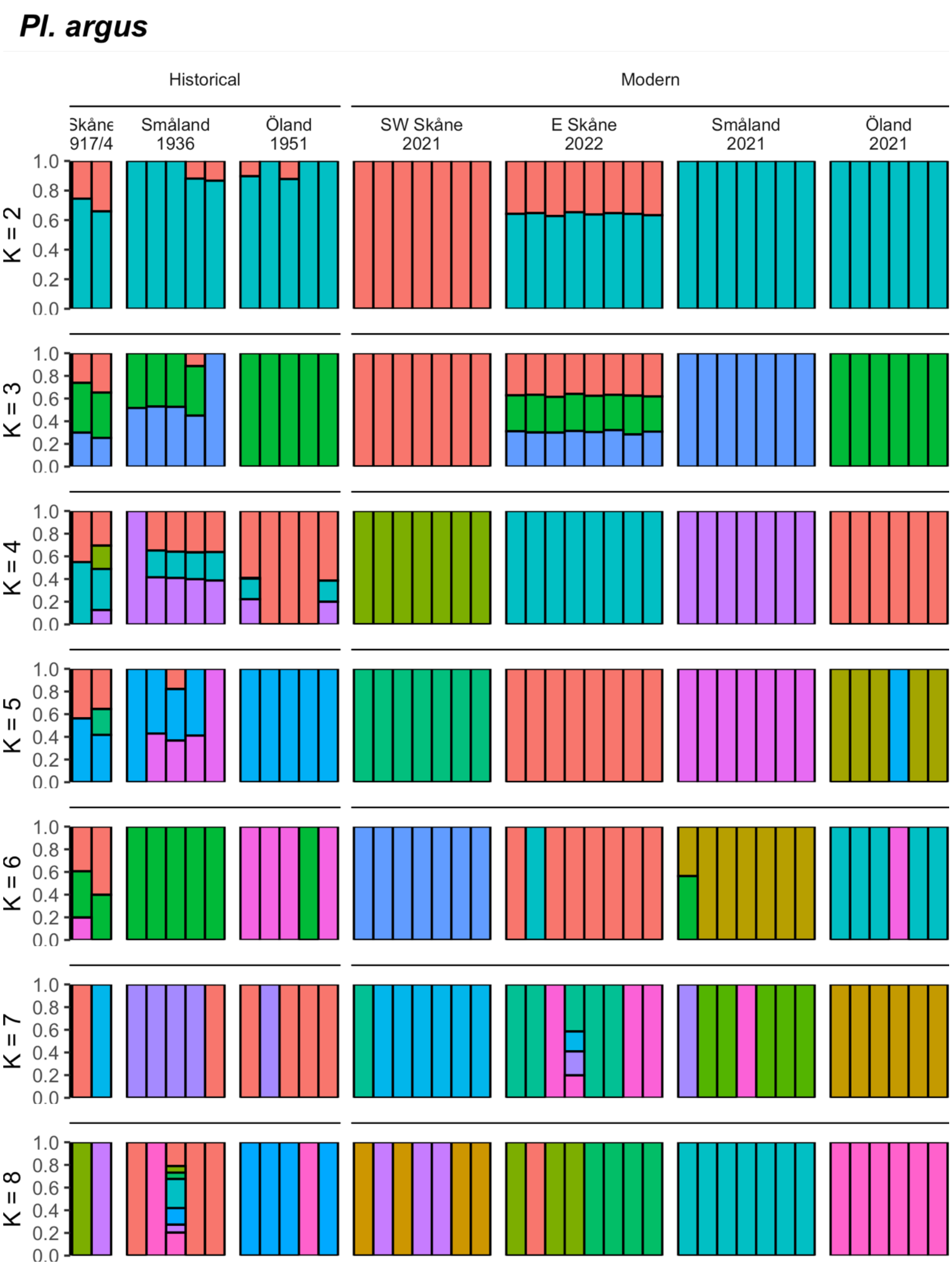

Pl. argus
Historical
Modern
Skåne 1917/4
Småland 1936
Öland 1951
SW Skåne 2021
E Skåne 2022
Småland 2021
Öland 2021
K = 2
K = 3
K = 4
K = 5
K = 6
K = 7
K = 8
1.0
0.8
0.6
0.4
0.2
0.0

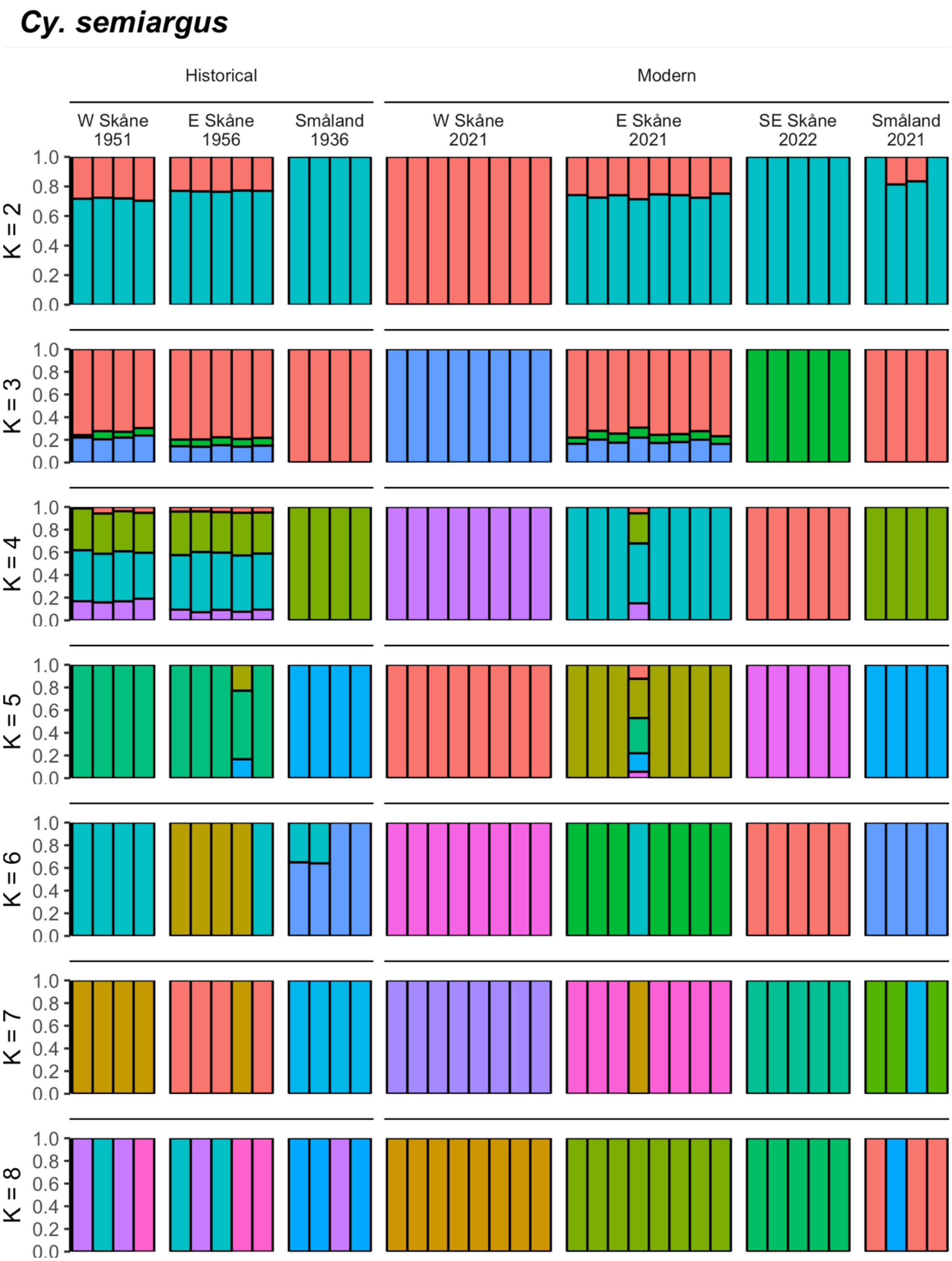

Cy. semiargus
Historical
Modern
W Skåne 1951
E Skåne 1956
Småland 1936
W Skåne 2021
E Skåne 2021
SE Skåne 2022
Småland 2021
K = 2
K = 3
K = 4
K = 5
K = 6
K = 7
K = 8
0.0
0.2
0.4
0.6
0.8
1.0

**Figure S6. Admixture analysis results for K 2-8 for the three focal species.** For each species, an admixture analysis was run for 2-8 clusters (K), with the optimized results for each value of K displayed in the figure above. We selected the best fitting K as the highest value of K that converged within 100 replicate runs of NGSadmix, following Pečnerová et al (*106*). The best fit values of K are 1, 1, and 5 for *Po. icarus*, *Pl. argus*, and *Cy. semiargus*, respectively, with the best fit models depicted in Figure 2 in the main text for *Cy. semiargus* and K = 2 for *Po. icarus* and *Pl. argus*.

***Po. icarus***

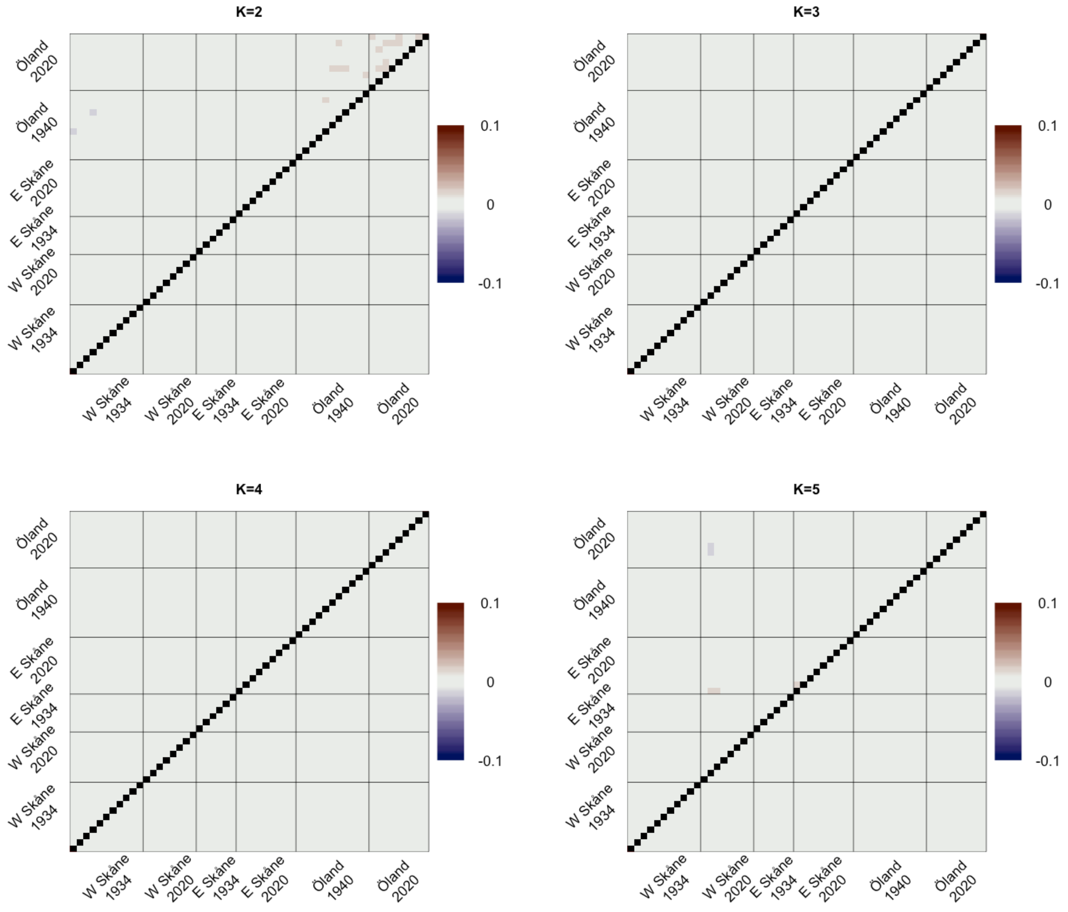
K=2
K=3
K=4
K=5
Öland 2020
Öland 1940
E Skåne 2020
E Skåne 1934
W Skåne 2020
W Skåne 1934
0.1
0
-0.1

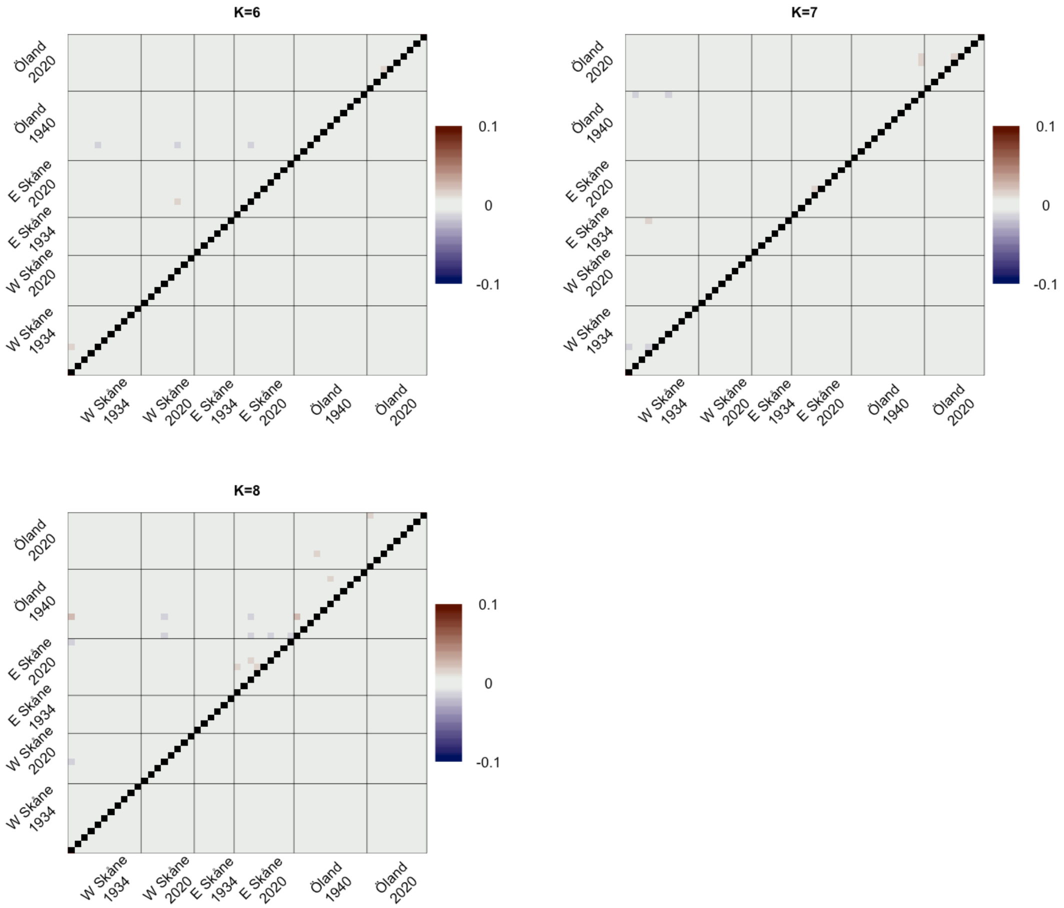

K=6
K=7
K=8
Öland 2020
Öland 1940
E Skåne 2020
E Skåne 1934
W Skåne 2020
W Skåne 1934
0.1
0
-0.1

*Pl. argus*

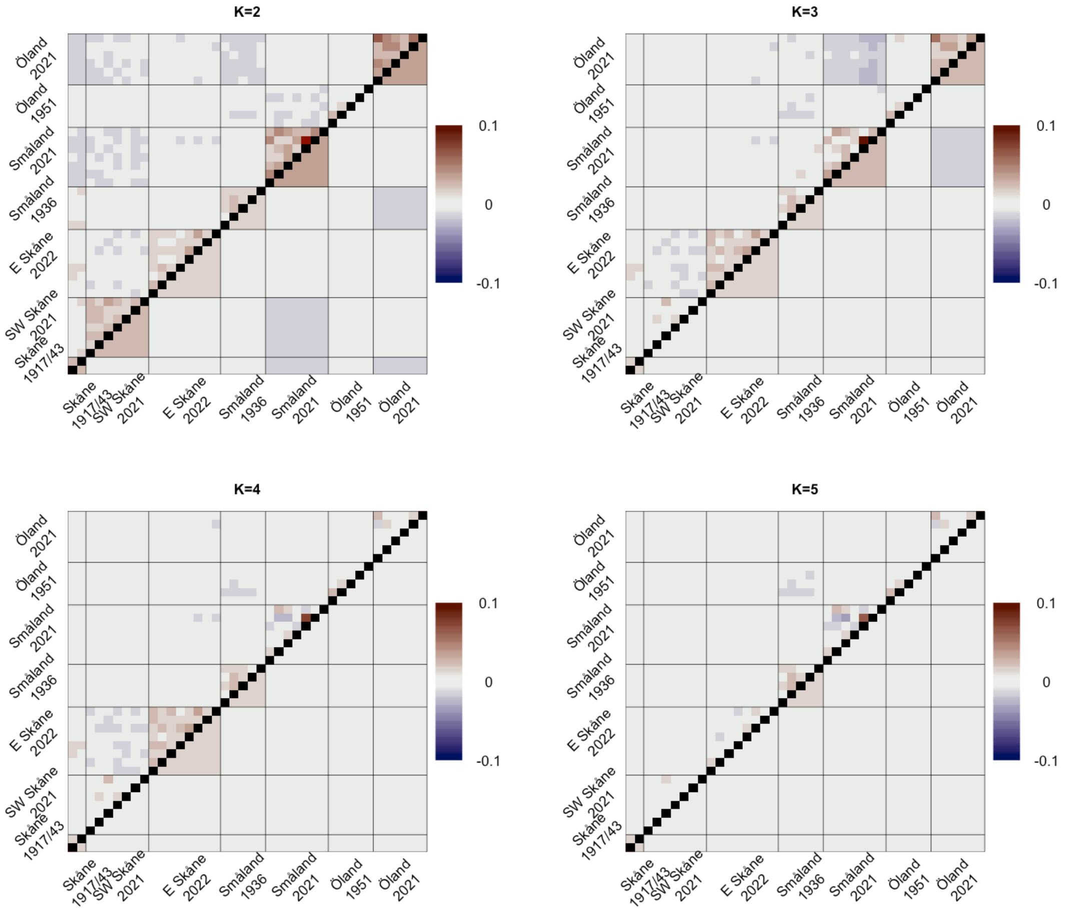
K=2
K=3
K=4
K=5
Skåne 1917/43
SW Skåne 2021
E Skåne 2022
Småland 1936
Småland 2021
Öland 1951
Öland 2021
0.1
0
-0.1

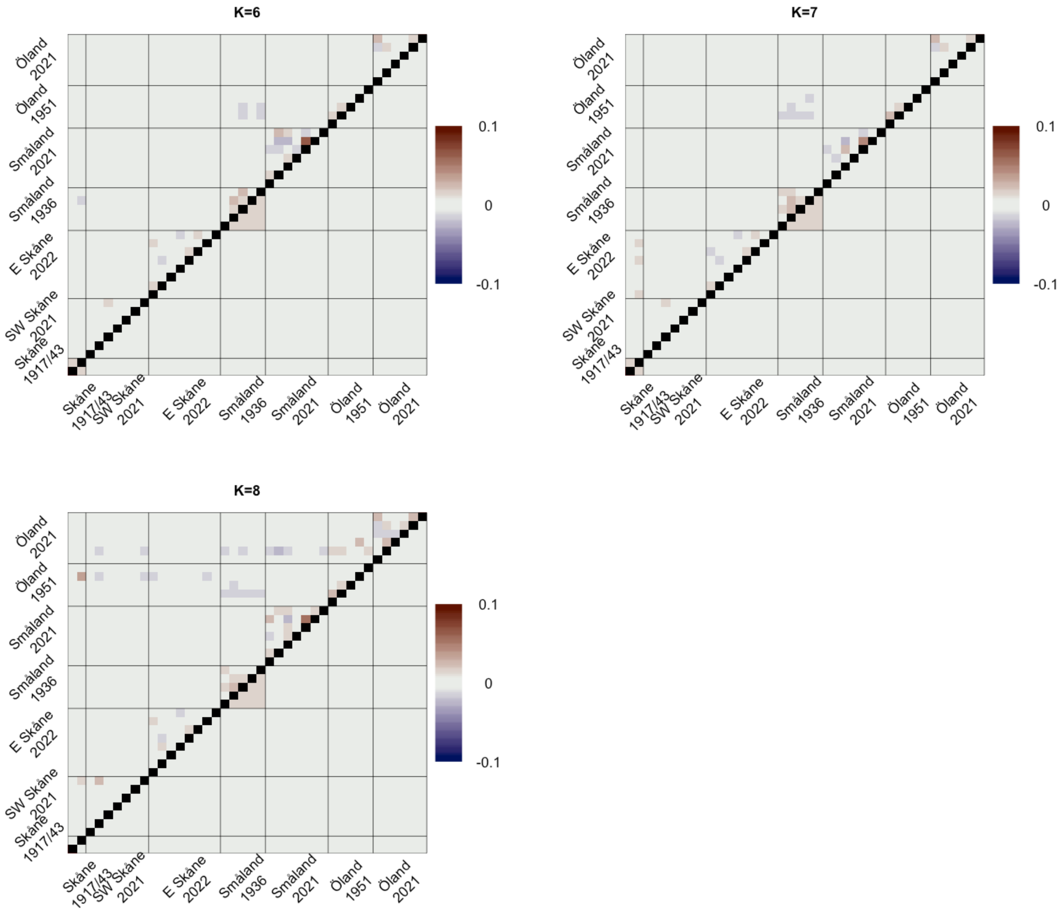
K=6
K=7
K=8
Skåne 1917/43
SW Skåne 2021
E Skåne 2022
Småland 1936
Småland 2021
Öland 1951
Öland 2021
0.1
0
-0.1

***Cy. semiargus***

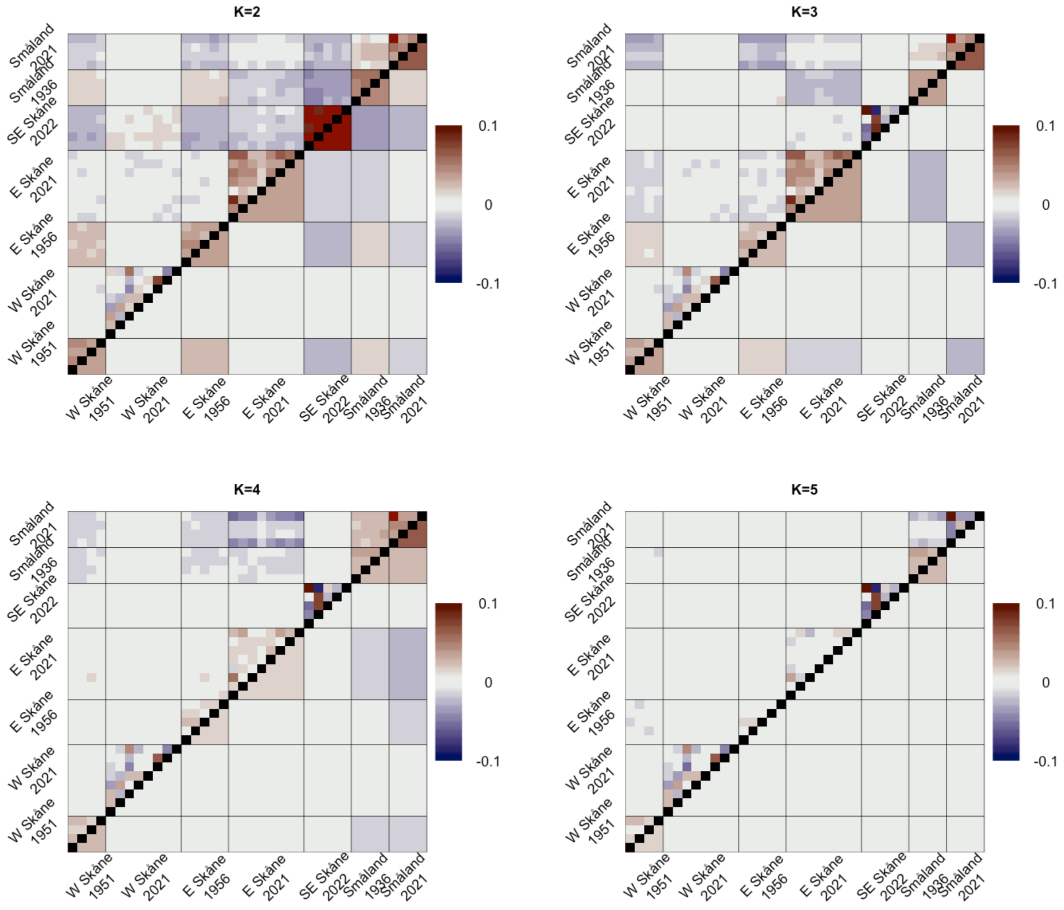

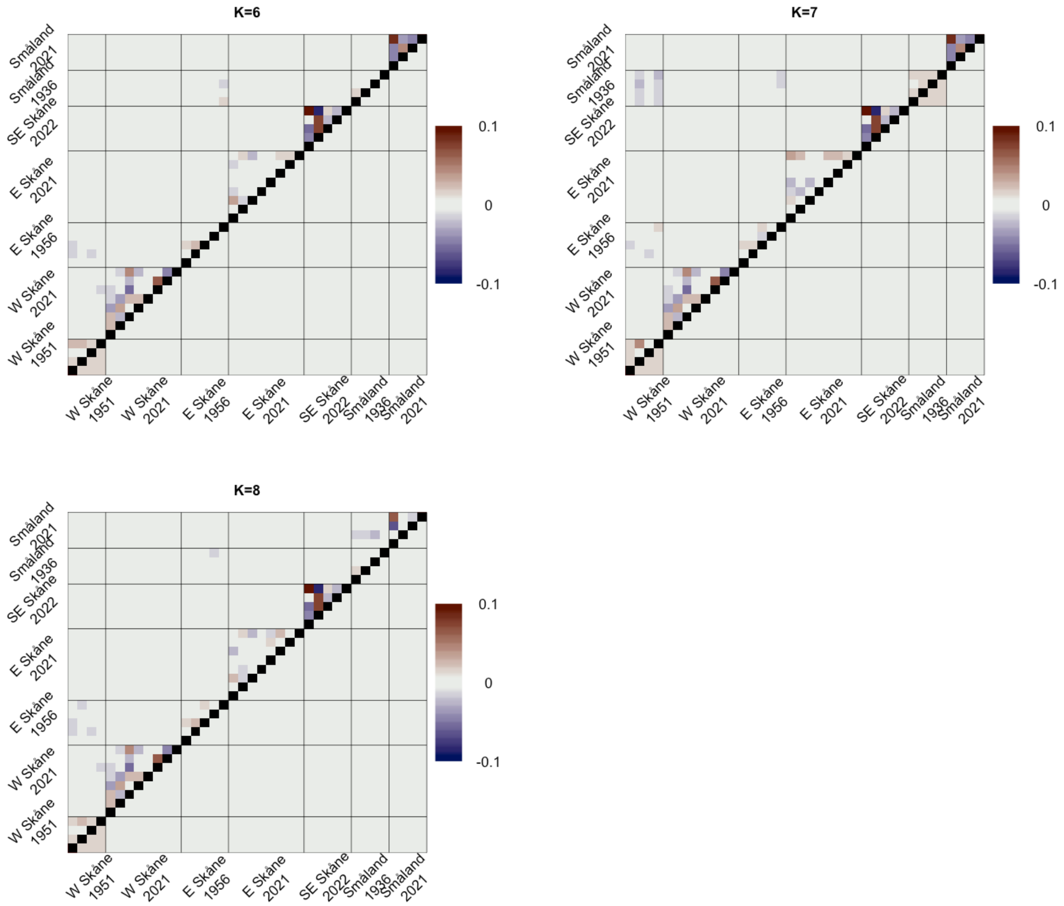


**Figure S7. Correlation of residuals for the focal species' admixture analyses.** For each value of K in the three focal species, we assessed the goodness of fit of our data to that K model of admixture using evalAdmix (*107*), visualized here with the pairwise correlation of residuals between individuals (upper left triangle of the matrix). When individuals are more similar than the model suggests, they have a positive pairwise correlation, when they are more different, they have a negative. Well-fitting sample pairs have a score of 0. The lower right triangle of the matrix shows the average scores of individuals in our *a priori* population groupings, illustrating when it is largely a population's fit that is in need of improvement. While we select K = 1 as our best fit based on our convergence criteria for *Po. icarus* and *Pl. argus*, we see that the data does not necessarily fit the model well in *Pl. argus*, illustrating that there is some genetic structure, as supported by the PCA in Figure 2. Our inability to achieve convergence for K-values above 2 in these species may relate to additional clusters being explained by a relatively small proportion of the genetic variance, requiring higher sample sizes to accurately assign admixture proportions at higher Ks. We do select a higher K, K = 5, for *Cy. semiargus*, which shows relatively good fit.

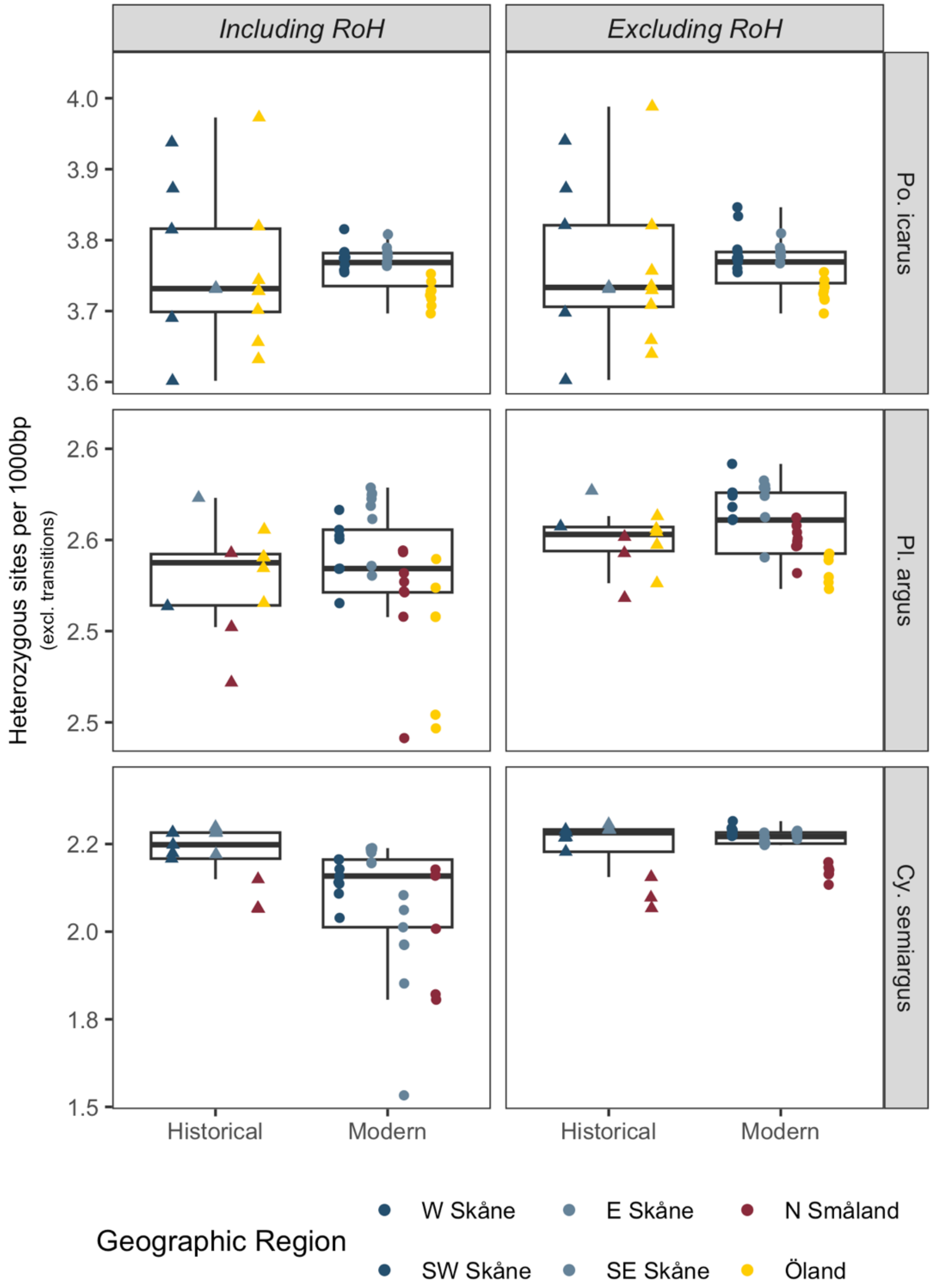


**Figure S8. Heterozygosity estimates including and excluding runs of homozygosity.** To assess the extent by which runs of homozygosity contributed to lower estimates of heterozygosity in modern samples, we calculated heterozygosity adjusted by $F_{RoH}$ as: Heterozygosity / (1 - $F_{RoH}$). This adjusted estimate provides an approximation of the heterozygosity expected outside of, or excluding *RoH*. This figure depicts the whole genome estimates of heterozygosity on the left (including *RoH*) and the estimates adjusted by $F_{RoH}$ on the right (excluding *RoH*). Points represent individuals, colored by sampling population, with boxplots encompassing all individuals for a given time period, as in Figure 1. Much of the difference in heterozygosity in *Cy. semiargus* visible from the whole genome estimates on the left is accounted for by runs of homozygosity, as the differences, even to the population level, are absent once heterozygosity is adjusted for *RoH* on the right.

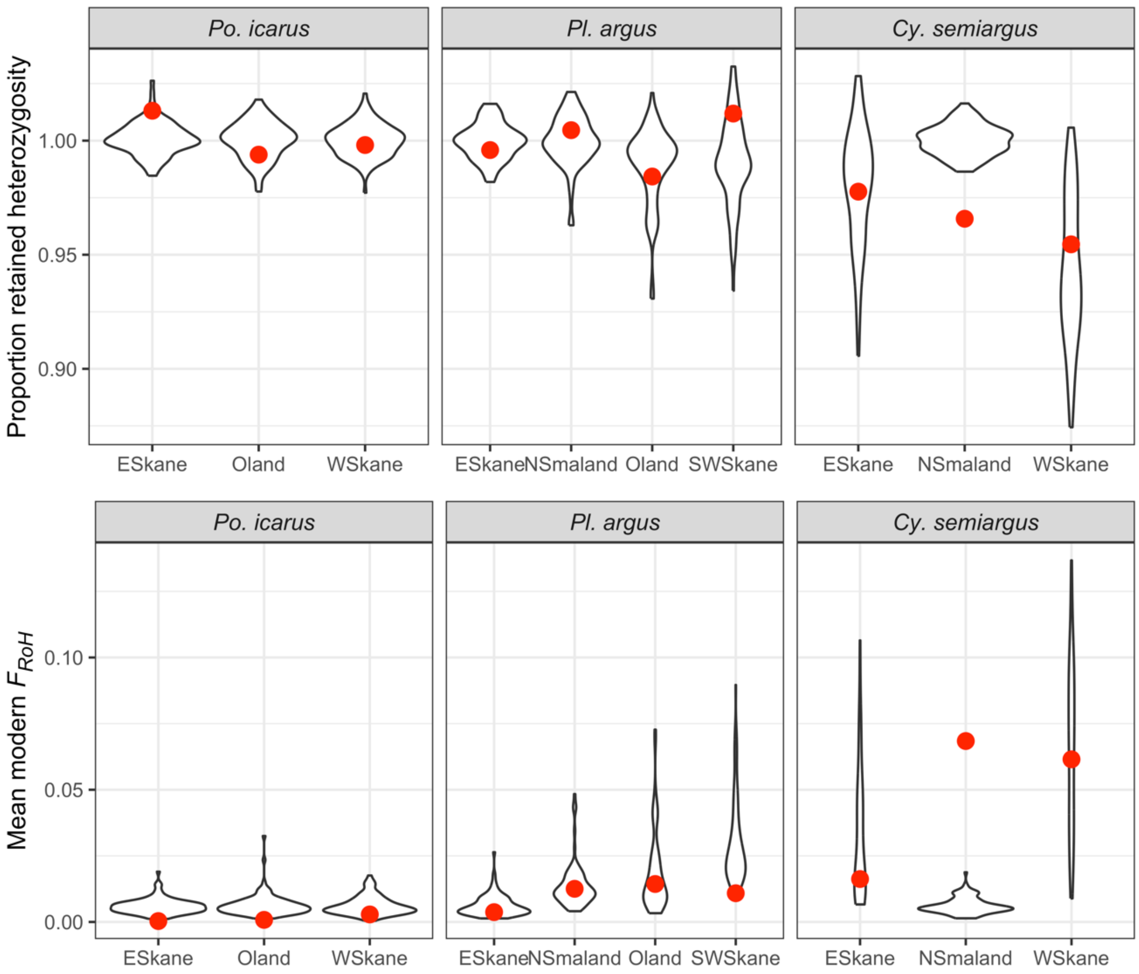


**Figure S9. Simulated expectations for retained heterozygosity and modern $F_{RoH}$.** To assess if the patterns of reduction in heterozygosity and increase in inbreeding coefficients we estimated empirically from historical and modern genomes were reasonable under population genetic expectations, we performed coalescent simulations for each population based on the effective population size trajectories estimated from the modern samples with GONE. Depicted here are violin plots for the 100 replicate runs of each simulated population history describing the distribution of heterozygosity retained in the modern population compared to the historical (upper), and the mean inbreeding coefficient of the simulated modern individuals (lower). Red points represent our empirical estimates of retained heterozygosity and modern inbreeding coefficients from each population. All empirical estimates fall within expected ranges, except for in Småland *C. semiargus* individuals. However, as empirical modern individual heterozygosity in Småland is similar to historical estimates after adjustment for $F_{RoH}$, we suggest that this deviation likely suggests the empirical estimates are reasonable and the modeled history may not reflect the actual history.

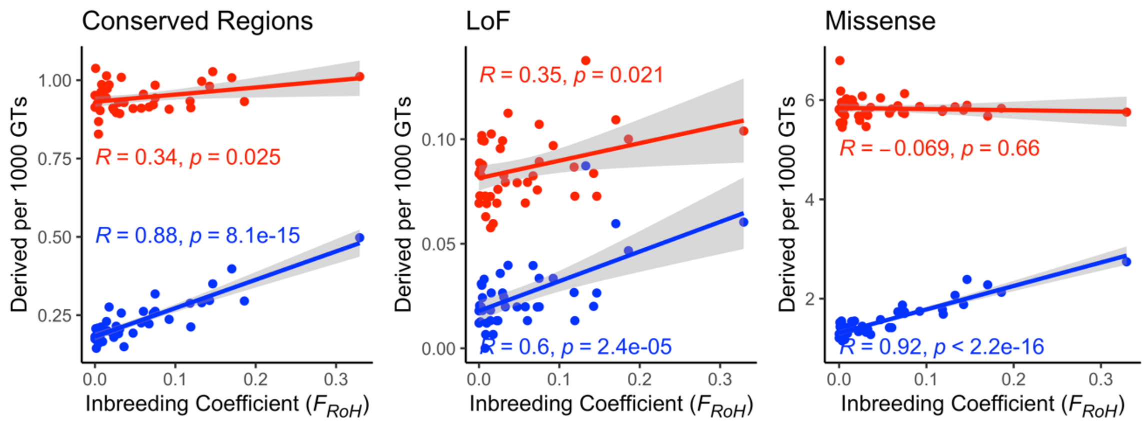


**Figure S10. Linear models examining the relationship between individual inbreeding coefficient and deleterious burden in *Cy. semiargus* across three variant subsets.** Red points and lines depict total estimates of deleterious burden, measured as counts of derived alleles for the given putatively deleterious category per 1000 genotyped positions. Blue points and lines depict homozygous deleterious burden, measured in the same way, but only counting homozygous derived alleles in the numerator. Total deleterious burden has a weak, but significant relationship with inbreeding coefficients for conserved and loss of function variants, whereas homozygous deleterious burden is positively related to inbreeding coefficient across all variant categories.

**(A) Multi-sample BCF with MAF > 0, without --ignore-homref**

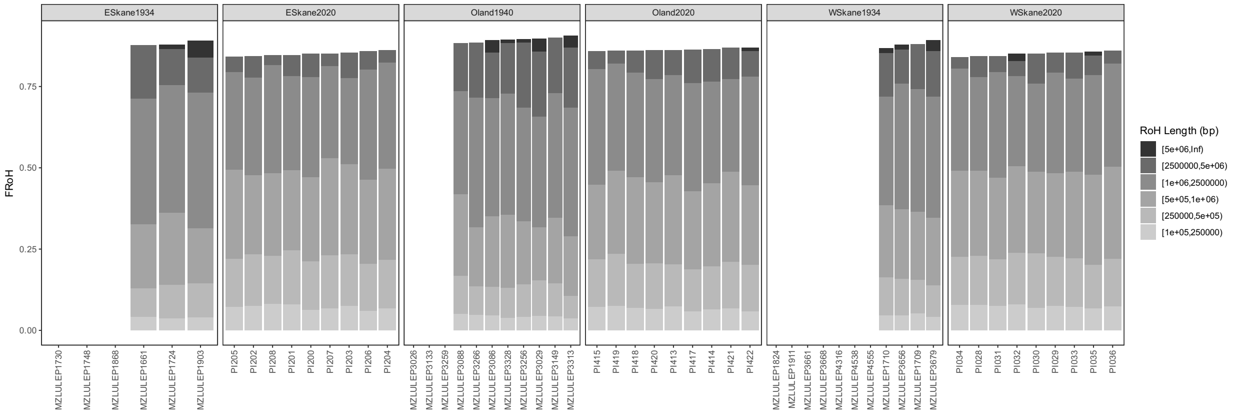


**(B) Multi-sample BCF with MAF > 0, with --ignore-homref (method in manuscript)**

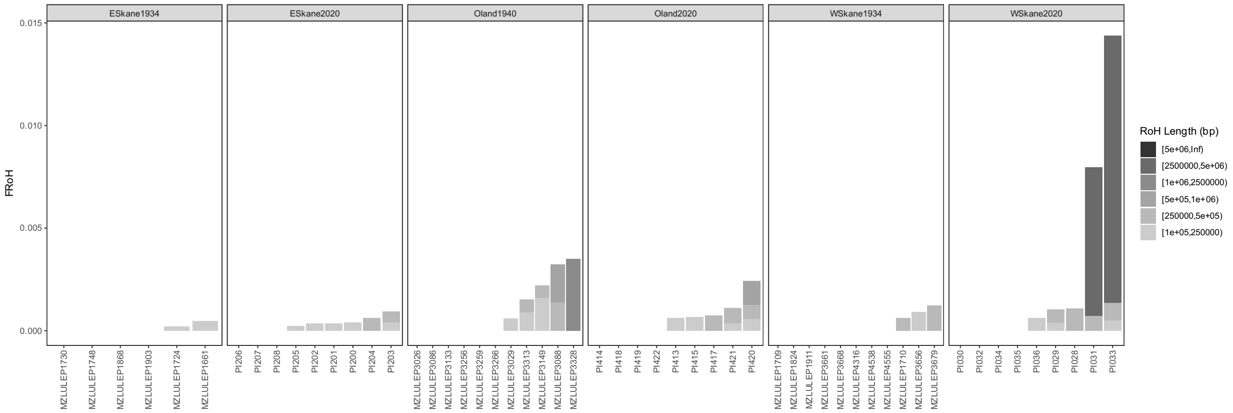


**Figure S11. Impacts of including homozygous reference genotypes when estimating runs of homozygosity.** BCFtools/RoH utilizes deviations from Hardy-Weinberg to detect runs of homozygosity, requiring an allele frequency for each variant to estimate this. In datasets where allele frequencies are unknown, a default can be used, commonly a minor allele frequency of 0.4. However, singleton and other low frequency variants deviate considerably from this default, and can heavily inflate the estimated runs of homozygosity, as in (A) where nearly all individuals of *Po. icarus* in our dataset are described as having almost the entire genome in runs of homozygosity. Two potential mechanisms to address this would be to estimate allele frequencies for the populations and supply them (as we previously did for the modern samples of *Po. icarus* in (*44*)), or filter out low frequency variants with a minor allele frequency filter. We have considerable variation in sample sizes across our species datasets, making a consistent MAF filter or comparable allele frequency estimates challenging, so we did not use either of these approaches. Instead, we utilized the option to ignore homozygous reference variants, with the results shown in (B). This reduces the effect, as the inflation largely comes from homozygous reference variants present in the BCF due to rare, alternate variants present in other individuals. When performed in this way, the analysis is similar to that done for variants called from a single sample the single sample as described in BCFtools online guide (https://samtools.github.io/bcftools/howtos/roh-calling.html). Using this approach, the estimates we get for all individuals show less than 1.5% of the genome in runs of homozygosity for all individuals, in line with the estimates we generated using population specific allele frequencies in (*44*). One main difference to performing the analysis on a single sample is that by using a multi-sample BCF, we filter out alternate variants that are fixed in the study individuals, as the reference allele is unlikely segregating, or segregating in low frequency, in the study populations, and would deviate from our default allele frequency.

**Table S1. Sample collection and sequencing information.** Each sample utilized in this study is listed here, providing its sample ID as referenced in the manuscript text/figures/code. Most specimens were registered into the entomological collections of the Biological Museum at Lund University, and has its museum identifier provided (MZLU ID). Each specimen is assigned a Time Period, which corresponds to whether it is being treated in the Historical or Modern sample grouping. As some samples in the Modern category were pinned specimens, the origin of the tissue used for DNA extraction is provided (Tissue Source). Study Region refers to the population grouping for population level analyses, grouping samples in the same study region and time period together. Sampling Locality refers to the name of the locality in which the sample was collected, either using the authors' designation for a sampling locality for fresh specimens or using the locality listed on the label for pinned specimens. For both, latitude and longitude of the locality is provided. For fresh specimens, these values correspond to the specific location at which butterflies were sampled, for pinned specimens these coordinates are inferred from the locality listed on the label, which is usually the nearest parish from the actual sampled grassland patch. For both fresh and pinned samples, the collection date and collector are provided, with collection dates that could not be fully inferred from the label having missing information replaced with 'xx'. For each sample, we report the total percent of sequencing reads mapping to the reference genome (% Mapped Reads) both when mapped with bwa mem/aln and vg, as well as the mean sequencing depth after filtering for each aligner.

| Sample ID | MZLU ID | Species | Time Period | Tissue Source | Sex | Collect Date | Study Region | Sampling Locality | Latitude | Longitude | Collector | % Mapped Reads (bwa) | % Mapped Reads (vg) | Filtered Depth (bwa aln/mem) | Filtered Depth (vg) |
|---|---|---|---|---|---|---|---|---|---|---|---|---|---|---|---|
| MZLULEP1724 | MZLULEP1724 | *Polyommatus icarus* | Historical | Pinned Specimen (abdomen) | M | 1934-06-03 | E Skåne | Kivik | 55.685 | 14.2266 | P. Benander | 90.25 | 100 | 5.3542 | 5.8738 |
| MZLULEP1868 | MZLULEP1868 | *Polyommatus icarus* | Historical | Pinned Specimen (abdomen) | M | 1934-06-03 | E Skåne | Kivik | 55.685 | 14.2266 | P. Benander | 89.81 | 100 | 2.0647 | 2.2375 |
| MZLULEP1730 | MZLULEP1730 | *Polyommatus icarus* | Historical | Pinned Specimen (abdomen) | M | 1934-06-06 | E Skåne | Kivik | 55.685 | 14.2266 | P. Benander | 90.26 | 100 | 3.2782 | 3.5842 |
| MZLULEP1748 | MZLULEP1748 | *Polyommatus icarus* | Historical | Pinned Specimen (abdomen) | M | 1934-06-06 | E Skåne | Kivik | 55.685 | 14.2266 | P. Benander | 89.23 | 100 | 3.2075 | 3.5499 |
| MZLULEP1661 | MZLULEP1661 | *Polyommatus icarus* | Historical | Pinned Specimen (abdomen) | M | 1934-07-25 | E Skåne | Kivik | 55.685 | 14.2266 | P. Benander | 87.85 | 100 | 6.0072 | 6.6505 |
| MZLULEP1903 | MZLULEP1903 | *Polyommatus icarus* | Historical | Pinned Specimen (abdomen) | M | 1934-07-25 | E Skåne | Kivik | 55.685 | 14.2266 | P. Benander | 89.81 | 100 | 5.6560 | 6.2317 |
| MZLULEP3086 | MZLULEP3086 | *Polyommatus icarus* | Historical | Pinned Specimen (abdomen) | M | 1940-07-28 | Öland | Vickleby | 56.5766 | 16.4604 | Winblad, Gustaf | 89 | 100 | 6.7512 | 7.5855 |
| MZLULEP3149 | MZLULEP3149 | *Polyommatus icarus* | Historical | Pinned Specimen (abdomen) | M | 1940-07-28 | Öland | Vickleby | 56.5766 | 16.4604 | Winblad, Gustaf | 89.19 | 100 | 10.6416 | 11.7809 |
| MZLULEP3328 | MZLULEP3328 | *Polyommatus icarus* | Historical | Pinned Specimen (abdomen) | M | 1940-07-28 | Öland | Vickleby | 56.5766 | 16.4604 | Winblad, Gustaf | 87.93 | 100 | 8.5818 | 9.6501 |
| MZLULEP3026 | MZLULEP3026 | *Polyommatus icarus* | Historical | Pinned Specimen (abdomen) | M | 1940-07-30 | Öland | Vickleby | 56.5766 | 16.4604 | Winblad, Gustaf | 88 | 100 | 3.9545 | 4.4624 |
| MZLULEP3029 | MZLULEP3029 | *Polyommatus icarus* | Historical | Pinned Specimen (abdomen) | M | 1940-07-30 | Öland | Vickleby | 56.5766 | 16.4604 | Winblad, Gustaf | 86.41 | 100 | 14.8130 | 16.7284 |
| MZLULEP3133 | MZLULEP3133 | *Polyommatus icarus* | Historical | Pinned Specimen (abdomen) | M | 1940-07-30 | Öland | Vickleby | 56.5766 | 16.4604 | Winblad, Gustaf | 87.84 | 100 | 4.2509 | 4.8654 |
| MZLULEP3256 | MZLULEP3256 | *Polyommatus icarus* | Historical | Pinned Specimen (abdomen) | M | 1940-07-30 | Öland | Vickleby | 56.5766 | 16.4604 | Winblad, Gustaf | 90.44 | 100 | 6.9215 | 7.5286 |
| MZLULEP3259 | MZLULEP3259 | *Polyommatus icarus* | Historical | Pinned Specimen (abdomen) | M | 1940-07-30 | Öland | Vickleby | 56.5766 | 16.4604 | Winblad, Gustaf | 86.78 | 100 | 4.2605 | 4.8570 |
| MZLULEP3266 | MZLULEP3266 | *Polyommatus icarus* | Historical | Pinned Specimen (abdomen) | M | 1940-07-30 | Öland | Vickleby | 56.5766 | 16.4604 | Winblad, Gustaf | 88.75 | 100 | 6.8630 | 7.7007 |
| MZLULEP3313 | MZLULEP3313 | *Polyommatus icarus* | Historical | Pinned Specimen (abdomen) | M | 1940-07-30 | Öland | Vickleby | 56.5766 | 16.4604 | Winblad, Gustaf | 86.47 | 100 | 4.9087 | 5.5662 |
| MZLULEP3088 | MZLULEP3088 | *Polyommatus icarus* | Historical | Pinned Specimen (abdomen) | M | 1940-08-01 | Öland | Vickleby | 56.5766 | 16.4604 | Winblad, Gustaf | 88.21 | 100 | 6.1489 | 6.9585 |
| MZLULEP4555 | MZLULEP4555 | *Polyommatus icarus* | Historical | Pinned Specimen (abdomen) | M | 1934-06-08 | W Skåne | Södra sandby | 55.7169 | 13.3472 | Ander, Kjell | 87.5 | 100 | 3.0313 | 3.3609 |
| MZLULEP3679 | MZLULEP3679 | *Polyommatus icarus* | Historical | Pinned Specimen (abdomen) | M | 1934-06-12 | W Skåne | Billebjer | 55.6883 | 13.3172 | Ander, Kjell | 90.1 | 100 | 8.6651 | 9.4829 |
| MZLULEP4538 | MZLULEP4538 | *Polyommatus icarus* | Historical | Pinned Specimen (abdomen) | M | 1934-06-24 | W Skåne | Kungsmarken | 55.7138 | 13.2753 | Ander, Kjell | 89.13 | 100 | 4.8134 | 5.2952 |
| MZLULEP3661 | MZLULEP3661 | *Polyommatus icarus* | Historical | Pinned Specimen (abdomen) | M | 1934-07-06 | W Skåne | Kungsmarken | 55.7138 | 13.2753 | Ander, Kjell | 88.41 | 100 | 3.5164 | 3.9514 |
| MZLULEP1710 | MZLULEP1710 | *Polyommatus icarus* | Historical | Pinned Specimen (abdomen) | M | 1934-08-02 | W Skåne | Södra sandby | 55.7169 | 13.3472 | Nordström, FGD | 86.05 | 100 | 5.0558 | 5.5778 |
| MZLULEP1824 | MZLULEP1824 | *Polyommatus icarus* | Historical | Pinned Specimen (abdomen) | M | 1934-08-02 | W Skåne | Södra sandby | 55.7169 | 13.3472 | Nordström, FGD | 90.33 | 100 | 3.3288 | 3.6008 |
| MZLULEP1709 | MZLULEP1709 | *Polyommatus icarus* | Historical | Pinned Specimen (abdomen) | M | 1934-08-20 | W Skåne | Södra sandby | 55.7169 | 13.3472 | Nordström, FGD | 86.77 | 99.99 | 5.6361 | 6.2998 |
| MZLULEP1911 | MZLULEP1911 | *Polyommatus icarus* | Historical | Pinned Specimen (abdomen) | M | 1934-08-20 | W Skåne | Dalby | 55.6648 | 13.3476 | Nordström, FGD | 64.56 | 99.93 | 3.6188 | 3.9413 |
| MZLULEP3656 | MZLULEP3656 | *Polyommatus icarus* | Historical | Pinned Specimen (abdomen) | M | 1934-08-31 | W Skåne | Hällestad | 55.6667 | 13.4167 | Ander, Kjell | 79.12 | 99.99 | 12.8158 | 14.2787 |
| MZLULEP3668 | MZLULEP3668 | *Polyommatus icarus* | Historical | Pinned Specimen (abdomen) | M | 1934-08-31 | W Skåne | Hällestad | 55.6667 | 13.4167 | Ander, Kjell | 88.73 | 99.99 | 4.0034 | 4.3815 |
| MZLULEP4316 | MZLULEP4316 | *Polyommatus icarus* | Historical | Pinned Specimen (abdomen) | M | 1934-08-31 | W Skåne | Torna Hällestad | 55.6781 | 13.4209 | Ander, Kjell | 63.95 | 99.92 | 3.1556 | 3.5317 |
| PI200 | MZLU00126549 | *Polyommatus icarus* | Modern | Field Caught | M | 2020-07-17 | E Skåne | Haväng | 55.7223 | 14.188 | ZJ Nolen | 99.45 | 96.54 | 25.8857 | 26.3887 |
| PI201 | MZLU00126550 | *Polyommatus icarus* | Modern | Field Caught | M | 2020-07-17 | E Skåne | Haväng | 55.7223 | 14.188 | ZJ Nolen | 99.48 | 96.61 | 25.6353 | 26.1393 |
| PI202 | MZLU00126551 | *Polyommatus icarus* | Modern | Field Caught | M | 2020-07-17 | E Skåne | Haväng | 55.7223 | 14.188 | ZJ Nolen | 99.42 | 96.51 | 26.5962 | 27.1144 |
| PI203 | MZLU00126552 | *Polyommatus icarus* | Modern | Field Caught | M | 2020-07-17 | E Skåne | Haväng | 55.7223 | 14.188 | ZJ Nolen | 99.48 | 96.62 | 23.8196 | 24.2916 |
| PI204 | MZLU00126553 | *Polyommatus icarus* | Modern | Field Caught | M | 2020-07-17 | E Skåne | Haväng | 55.7223 | 14.188 | ZJ Nolen | 99.46 | 96.58 | 30.1519 | 30.7385 |
| PI205 | MZLU00126554 | *Polyommatus icarus* | Modern | Field Caught | M | 2020-07-17 | E Skåne | Haväng | 55.7223 | 14.188 | ZJ Nolen | 98.8 | 96.08 | 21.0982 | 21.5142 |
| PI206 | MZLU00126555 | *Polyommatus icarus* | Modern | Field Caught | M | 2020-07-19 | E Skåne | Haväng | 55.7223 | 14.188 | ZJ Nolen | 96.93 | 94.69 | 25.7221 | 26.2343 |
| PI207 | MZLU00126556 | *Polyommatus icarus* | Modern | Field Caught | M | 2020-07-19 | E Skåne | Haväng | 55.7223 | 14.188 | ZJ Nolen | 99.03 | 96.35 | 26.4757 | 26.9919 |
| PI208 | MZLU00126557 | *Polyommatus icarus* | Modern | Field Caught | M | 2020-07-19 | E Skåne | Haväng | 55.7223 | 14.188 | ZJ Nolen | 98.83 | 96.05 | 26.9350 | 27.4674 |
| PI413 | MZLU00126558 | *Polyommatus icarus* | Modern | Field Caught | M | 2020-08-01 | Öland | Sandby Borg | 56.5532 | 16.6396 | ZJ Nolen | 99.4 | 96.02 | 44.1365 | 45.0132 |
| PI414 | MZLU00126559 | *Polyommatus icarus* | Modern | Field Caught | M | 2020-08-01 | Öland | Sandby Borg | 56.5532 | 16.6396 | ZJ Nolen | 99.41 | 95.96 | 47.9536 | 48.8774 |
| PI415 | MZLU00126560 | *Polyommatus icarus* | Modern | Field Caught | M | 2020-08-01 | Öland | Sandby Borg | 56.5532 | 16.6396 | ZJ Nolen | 99.38 | 95.99 | 55.6663 | 56.7365 |
| PI417 | MZLU00126561 | *Polyommatus icarus* | Modern | Field Caught | M | 2020-08-01 | Öland | Sandby Borg | 56.5532 | 16.6396 | ZJ Nolen | 99.39 | 96.05 | 49.5689 | 50.5379 |
| PI418 | MZLU00126562 | *Polyommatus icarus* | Modern | Field Caught | M | 2020-08-01 | Öland | Sandby Borg | 56.5532 | 16.6396 | ZJ Nolen | 99.07 | 95.61 | 45.7502 | 46.6659 |
| PI419 | MZLU00126563 | *Polyommatus icarus* | Modern | Field Caught | M | 2020-08-01 | Öland | Sandby Borg | 56.5532 | 16.6396 | ZJ Nolen | 99.4 | 96.14 | 43.7582 | 44.5946 |
| PI420 | MZLU00126564 | *Polyommatus icarus* | Modern | Field Caught | M | 2020-08-01 | Öland | Sandby Borg | 56.5532 | 16.6396 | ZJ Nolen | 99.39 | 96.02 | 38.3690 | 39.1139 |
| PI421 | MZLU00126565 | *Polyommatus icarus* | Modern | Field Caught | M | 2020-08-01 | Öland | Sandby Borg | 56.5532 | 16.6396 | ZJ Nolen | 99.41 | 95.96 | 45.4122 | 46.3059 |
| PI422 | MZLU00126566 | *Polyommatus icarus* | Modern | Field Caught | M | 2020-08-01 | Öland | Sandby Borg | 56.5532 | 16.6396 | ZJ Nolen | 99.4 | 95.99 | 52.7506 | 53.7786 |
| PI028 | MZLU00126531 | *Polyommatus icarus* | Modern | Field Caught | M | 2020-07-22 | W Skåne | Tvedöra | 55.6951 | 13.4316 | ZJ Nolen | 98.65 | 95.95 | 21.4433 | 21.8670 |
| PI029 | MZLU00126532 | *Polyommatus icarus* | Modern | Field Caught | M | 2020-07-22 | W Skåne | Tvedöra | 55.6951 | 13.4316 | ZJ Nolen | 99.45 | 96.42 | 16.6019 | 16.9345 |
| PI030 | MZLU00126533 | *Polyommatus icarus* | Modern | Field Caught | M | 2020-07-22 | W Skåne | Tvedöra | 55.6951 | 13.4316 | ZJ Nolen | 99.44 | 96.58 | 27.8268 | 28.3772 |
| PI031 | MZLU00126534 | *Polyommatus icarus* | Modern | Field Caught | M | 2020-07-22 | W Skåne | Tvedöra | 55.6951 | 13.4316 | ZJ Nolen | 99.36 | 96.53 | 24.9953 | 25.4878 |
| PI032 | MZLU00126535 | *Polyommatus icarus* | Modern | Field Caught | M | 2020-07-22 | W Skåne | Tvedöra | 55.6951 | 13.4316 | ZJ Nolen | 99.46 | 96.72 | 29.3019 | 29.8777 |

| | | | | | | | | | | | | | | | |
|---|---|---|---|---|---|---|---|---|---|---|---|---|---|---|---|
| PI033 | MZLU00126536 | *Polyommatus icarus* | Modern | Field Caught | M | 2020-07-22 | W Skåne | Tvedöra | 55.6951 | 13.4316 | ZJ Nolen | 99.46 | 96.53 | 23.5182 | 23.9842 |
| PI034 | MZLU00126537 | *Polyommatus icarus* | Modern | Field Caught | M | 2020-07-22 | W Skåne | Tvedöra | 55.6951 | 13.4316 | ZJ Nolen | 99.43 | 96.65 | 24.1715 | 24.6394 |
| PI035 | MZLU00126538 | *Polyommatus icarus* | Modern | Field Caught | M | 2020-07-22 | W Skåne | Tvedöra | 55.6951 | 13.4316 | ZJ Nolen | 99.27 | 96.45 | 26.6241 | 27.1400 |
| PI036 | MZLU00126539 | *Polyommatus icarus* | Modern | Field Caught | M | 2020-07-22 | W Skåne | Tvedöra | 55.6951 | 13.4316 | ZJ Nolen | 99.44 | 96.5 | 21.7631 | 22.1901 |
| MZLU153003 | MZLU153003 | *Plebejus argus* | Historical | Pinned Specimen (abdomen) | M | 1917-06-18 | E Skåne | Åhus | 55.9231 | 14.2942 | Wahlgren, Einar | 86.18 | 99.99 | 5.9404 | 6.5176 |
| MZLU153166 | MZLU153166 | *Plebejus argus* | Historical | Pinned Specimen (abdomen) | M | 1951-07-26 | Öland | Gärdslösa | 56.7842 | 16.7392 | H. Rambring | 88.25 | 100 | 6.7504 | 7.4776 |
| MZLU153167 | MZLU153167 | *Plebejus argus* | Historical | Pinned Specimen (abdomen) | M | 1951-07-26 | Öland | Gärdslösa | 56.7842 | 16.7392 | H. Rambring | 89.11 | 99.99 | 5.1508 | 5.6644 |
| MZLU153168 | MZLU153168 | *Plebejus argus* | Historical | Pinned Specimen (abdomen) | M | 1951-07-26 | Öland | Gärdslösa | 56.7842 | 16.7392 | H. Rambring | 87.21 | 100 | 5.5830 | 6.3410 |
| MZLU153169 | MZLU153169 | *Plebejus argus* | Historical | Pinned Specimen (abdomen) | M | 1951-07-26 | Öland | Gärdslösa | 56.7842 | 16.7392 | H. Rambring | 88.84 | 99.99 | 7.1448 | 7.7978 |
| MZLU153170 | MZLU153170 | *Plebejus argus* | Historical | Pinned Specimen (abdomen) | M | 1951-07-26 | Öland | Gärdslösa | 56.7842 | 16.7392 | H. Rambring | 87.86 | 99.99 | 7.6939 | 8.5635 |
| MZLU153046 | MZLU153046 | *Plebejus argus* | Historical | Pinned Specimen (abdomen) | M | 1936-06-27 | Småland | Hjelmseryd | 57.3112 | 14.5131 | Carlgren G. | 88.52 | 99.99 | 5.7848 | 6.5124 |
| MZLU153040 | MZLU153040 | *Plebejus argus* | Historical | Pinned Specimen (abdomen) | M | 1936-07-01 | Småland | Hjelmseryd | 57.3112 | 14.5131 | Carlgren G. | 88.43 | 99.99 | 4.3350 | 4.7956 |
| MZLU153045 | MZLU153045 | *Plebejus argus* | Historical | Pinned Specimen (abdomen) | M | 1936-07-06 | Småland | Hjelmseryd | 57.3112 | 14.5131 | Carlgren G. | 81.7 | 99.94 | 3.4038 | 3.7513 |
| MZLU153042 | MZLU153042 | *Plebejus argus* | Historical | Pinned Specimen (abdomen) | M | 1936-07-07 | Småland | Hjelmseryd | 57.3112 | 14.5131 | Carlgren G. | 88.85 | 100 | 4.9652 | 5.5416 |
| MZLU153048 | MZLU153048 | *Plebejus argus* | Historical | Pinned Specimen (abdomen) | M | 1936-07-12 | Småland | Hjelmseryd | 57.3112 | 14.5131 | Carlgren G. | 88.98 | 99.99 | 6.4863 | 7.1991 |
| MZLU152940 | MZLU152940 | *Plebejus argus* | Historical | Pinned Specimen (abdomen) | M | 1943-07-03 | W Skåne | Hällestad | 55.6765 | 13.4192 | Ander, Kjell | 89.71 | 100 | 5.4304 | 6.0053 |
| PLAR0200 | MZLU00126622 | *Plebejus argus* | Modern | Field Caught | M | 2022-07-06 | E Skåne | Drakamöllan | 55.7551 | 14.1252 | Melina Eberhagen | 99.24 | 96.28 | 16.3539 | 16.5749 |
| PLAR0202 | MZLU00126623 | *Plebejus argus* | Modern | Field Caught | M | 2022-07-06 | E Skåne | Drakamöllan | 55.7551 | 14.1252 | Melina Eberhagen | 99.23 | 96.33 | 11.5077 | 11.6594 |
| PLAR0203 | MZLU00126624 | *Plebejus argus* | Modern | Field Caught | M | 2022-07-06 | E Skåne | Drakamöllan | 55.7551 | 14.1252 | Melina Eberhagen | 99.23 | 96.15 | 10.1628 | 10.3002 |
| PLAR0208 | MZLU00126625 | *Plebejus argus* | Modern | Field Caught | M | 2022-07-06 | E Skåne | Drakamöllan | 55.7551 | 14.1252 | Melina Eberhagen | 99.2 | 96.06 | 8.8572 | 8.9760 |
| PLAR0211 | MZLU00126626 | *Plebejus argus* | Modern | Field Caught | M | 2022-07-06 | E Skåne | Drakamöllan | 55.7551 | 14.1252 | Melina Eberhagen | 99.21 | 96.31 | 10.1285 | 10.2634 |
| PLAR0217 | MZLU00126627 | *Plebejus argus* | Modern | Field Caught | M | 2022-07-06 | E Skåne | Drakamöllan | 55.7551 | 14.1252 | Melina Eberhagen | 99.22 | 96.2 | 11.8572 | 12.0063 |
| PLAR0222 | MZLU00126628 | *Plebejus argus* | Modern | Field Caught | M | 2022-07-06 | E Skåne | Drakamöllan | 55.7551 | 14.1252 | Melina Eberhagen | 99.21 | 96.14 | 10.6144 | 10.7523 |
| PLAR0216 | MZLU00126629 | *Plebejus argus* | Modern | Field Caught | M | 2022-07-06 | E Skåne | Drakamöllan | 55.7551 | 14.1252 | Melina Eberhagen | 99.23 | 99.23 | 11.1085 | 11.1087 |
| MZLU107434 | MZLU107434 | *Plebejus argus* | Modern | Field Caught | M | 2021-06-24 | Småland | Götafors | 57.492 | 14.1134 | ZJ Nolen | 99.26 | 95.36 | 18.5898 | 18.9344 |
| MZLU107438 | MZLU107438 | *Plebejus argus* | Modern | Field Caught | M | 2021-06-24 | Småland | Götafors | 57.492 | 14.1134 | ZJ Nolen | 99.24 | 94.79 | 24.3515 | 24.8322 |
| MZLU107435 | MZLU107435 | *Plebejus argus* | Modern | Field Caught | M | 2021-07-07 | Småland | Götafors | 57.492 | 14.1134 | ZJ Nolen | 99.24 | 94.76 | 27.5812 | 28.1041 |
| MZLU107436 | MZLU107436 | *Plebejus argus* | Modern | Field Caught | M | 2021-07-07 | Småland | Götafors | 57.492 | 14.1134 | ZJ Nolen | 99.26 | 95.16 | 16.6838 | 17.0067 |
| MZLU107470 | MZLU107470 | *Plebejus argus* | Modern | Field Caught | M | 2021-06-24 | Småland | Götafors | 57.492 | 14.1134 | ZJ Nolen | 99.26 | 95.08 | 20.6732 | 21.0737 |
| MZLU107486 | MZLU107486 | *Plebejus argus* | Modern | Field Caught | M | 2021-06-24 | Småland | Götafors | 57.492 | 14.1134 | ZJ Nolen | 99.25 | 94.65 | 10.6846 | 10.9039 |
| MZLU107469 | MZLU107469 | *Plebejus argus* | Modern | Field Caught | M | 2021-07-07 | Småland | Götafors | 57.492 | 14.1134 | ZJ Nolen | 99.26 | 94.79 | 18.7570 | 19.1297 |
| MZLU107483 | MZLU107483 | *Plebejus argus* | Modern | Field Caught | M | 2021-07-07 | Småland | Götafors | 57.492 | 14.1134 | ZJ Nolen | 98.95 | 94.41 | 6.9052 | 7.0452 |
| MZLU107445 | MZLU107445 | *Plebejus argus* | Modern | Field Caught | M | 2021-07-17 | Öland | Jordtorpåsen | 56.6783 | 16.5511 | ZJ Nolen | 99.08 | 94.52 | 30.0482 | 30.6303 |
| MZLU107449 | MZLU107449 | *Plebejus argus* | Modern | Field Caught | M | 2021-07-17 | Öland | Jordtorpåsen | 56.6783 | 16.5511 | ZJ Nolen | 99.28 | 95.25 | 20.3436 | 20.7343 |
| MZLU107451 | MZLU107451 | *Plebejus argus* | Modern | Field Caught | M | 2021-07-17 | Öland | Jordtorpåsen | 56.6783 | 16.5511 | ZJ Nolen | 99.19 | 95.05 | 19.6817 | 20.0461 |
| MZLU107471 | MZLU107471 | *Plebejus argus* | Modern | Field Caught | M | 2021-07-17 | Öland | Jordtorpåsen | 56.6783 | 16.5511 | ZJ Nolen | 99.27 | 94.9 | 24.9822 | 25.4670 |
| MZLU107477 | MZLU107477 | *Plebejus argus* | Modern | Field Caught | M | 2021-07-17 | Öland | Jordtorpåsen | 56.6783 | 16.5511 | ZJ Nolen | 99.15 | 94.4 | 12.7584 | 13.0254 |
| MZLU107487 | MZLU107487 | *Plebejus argus* | Modern | Field Caught | M | 2021-07-17 | Öland | Jordtorpåsen | 56.6783 | 16.5511 | ZJ Nolen | 99.25 | 94.85 | 11.9836 | 12.2250 |
| MZLU107426 | MZLU107426 | *Plebejus argus* | Modern | Field Caught | M | 2021-07-12 | SW Skåne | Falsterbo | 55.4025 | 12.8853 | ZJ Nolen | 99.26 | 95.29 | 18.9762 | 19.3367 |
| MZLU107439 | MZLU107439 | *Plebejus argus* | Modern | Field Caught | M | 2021-07-12 | SW Skåne | Falsterbo | 55.4025 | 12.8853 | ZJ Nolen | 99.21 | 94.96 | 19.4143 | 19.7905 |
| MZLU107441 | MZLU107441 | *Plebejus argus* | Modern | Field Caught | M | 2021-07-12 | SW Skåne | Falsterbo | 55.4025 | 12.8853 | ZJ Nolen | 99.25 | 95.11 | 16.0225 | 16.3322 |
| MZLU107443 | MZLU107443 | *Plebejus argus* | Modern | Field Caught | M | 2021-07-12 | SW Skåne | Falsterbo | 55.4025 | 12.8853 | ZJ Nolen | 99.26 | 95.07 | 30.3852 | 30.9579 |
| MZLU107468 | MZLU107468 | *Plebejus argus* | Modern | Field Caught | M | 2021-07-12 | SW Skåne | Falsterbo | 55.4025 | 12.8853 | ZJ Nolen | 99.25 | 95.23 | 15.0256 | 15.3127 |
| MZLU107475 | MZLU107475 | *Plebejus argus* | Modern | Field Caught | M | 2021-07-12 | SW Skåne | Falsterbo | 55.4025 | 12.8853 | ZJ Nolen | 99.09 | 95.08 | 32.3835 | 32.9918 |
| MZLU107480 | MZLU107480 | *Plebejus argus* | Modern | Field Caught | M | 2021-07-12 | SW Skåne | Falsterbo | 55.4025 | 12.8853 | ZJ Nolen | 99.24 | 94.45 | 10.2806 | 10.4898 |
| MZLU153246 | MZLU153246 | *Cyaniris semiargus* | Historical | Pinned Specimen (abdomen) | M | 1956-07-21 | E Skåne | Degeberga | 55.838 | 14.0901 | H. Rambring | 87.79 | 99.98 | 7.5046 | 7.9044 |
| MZLU153247 | MZLU153247 | *Cyaniris semiargus* | Historical | Pinned Specimen (abdomen) | M | 1956-07-21 | E Skåne | Degeberga | 55.838 | 14.0901 | H. Rambring | 91.55 | 100 | 8.3716 | 8.9663 |
| MZLU153248 | MZLU153248 | *Cyaniris semiargus* | Historical | Pinned Specimen (abdomen) | M | 1956-07-21 | E Skåne | Degeberga | 55.838 | 14.0901 | H. Rambring | 88.75 | 99.99 | 7.0833 | 7.4967 |
| MZLU153251 | MZLU153251 | *Cyaniris semiargus* | Historical | Pinned Specimen (abdomen) | M | 1956-07-21 | E Skåne | Degeberga | 55.838 | 14.0901 | H. Rambring | 87.62 | 99.99 | 8.2955 | 8.8308 |
| MZLU153250 | MZLU153250 | *Cyaniris semiargus* | Historical | Pinned Specimen (abdomen) | M | 1957-07-21 | E Skåne | Degeberga | 55.838 | 14.0901 | H. Rambring | 92.47 | 99.99 | 9.1029 | 9.5331 |
| MZLU152770 | MZLU152770 | *Cyaniris semiargus* | Historical | Pinned Specimen (abdomen) | M | 1936-06-27 | Småland | Hjelmseryd | 57.3112 | 14.5131 | Carlgren G. | 90.47 | 99.99 | 6.0121 | 6.4092 |
| MZLU152769 | MZLU152769 | *Cyaniris semiargus* | Historical | Pinned Specimen (abdomen) | M | 1936-06-29 | Småland | Hjelmseryd | 57.3112 | 14.5131 | Carlgren G. | 90.34 | 100 | 11.1979 | 12.4081 |
| MZLU152765 | MZLU152765 | *Cyaniris semiargus* | Historical | Pinned Specimen (abdomen) | M | 1936-07-01 | Småland | Hjelmseryd | 57.3112 | 14.5131 | Carlgren G. | 92.27 | 99.99 | 3.2783 | 3.4713 |
| MZLU152766 | MZLU152766 | *Cyaniris semiargus* | Historical | Pinned Specimen (abdomen) | M | 1936-07-07 | Småland | Hjelmseryd | 57.3112 | 14.5131 | Carlgren G. | 92.17 | 99.99 | 4.9600 | 5.2615 |
| MZLU153204 | MZLU153204 | *Cyaniris semiargus* | Historical | Pinned Specimen (abdomen) | M | 1951-06-19 | W Skåne | Torna Hällestad | 55.6781 | 13.4214 | H. Rambring | 92.39 | 100 | 7.6933 | 8.0408 |
| MZLU153205 | MZLU153205 | *Cyaniris semiargus* | Historical | Pinned Specimen (abdomen) | M | 1951-06-19 | W Skåne | Torna Hällestad | 55.6781 | 13.4214 | H. Rambring | 89.85 | 99.99 | 7.4680 | 7.8478 |
| MZLU153206 | MZLU153206 | *Cyaniris semiargus* | Historical | Pinned Specimen (abdomen) | M | 1951-06-19 | W Skåne | Torna Hällestad | 55.6781 | 13.4214 | H. Rambring | 92.53 | 99.99 | 6.8859 | 7.2077 |
| MZLU153216 | MZLU153216 | *Cyaniris semiargus* | Historical | Pinned Specimen (abdomen) | M | 1951-06-19 | W Skåne | Torna Hällestad | 55.6781 | 13.4214 | H. Rambring | 87.08 | 99.97 | 5.8776 | 6.1767 |
| MZLU153221 | MZLU153221 | *Cyaniris semiargus* | Historical | Pinned Specimen (abdomen) | M | 1951-06-19 | W Skåne | Torna Hällestad | 55.6781 | 13.4214 | H. Rambring | 93.52 | 100 | 5.2265 | 5.4991 |
| MZLU107454 | MZLU107454 | *Cyaniris semiargus* | Modern | Field Caught | M | 2021-07-03 | E Skåne | Fästan | 55.776 | 14.1722 | ZJ Nolen; P Jamelska | 99.61 | 97.68 | 18.1545 | 18.3237 |
| MZLU107458 | MZLU107458 | *Cyaniris semiargus* | Modern | Field Caught | M | 2021-07-03 | E Skåne | Fästan | 55.776 | 14.1722 | ZJ Nolen; P Jamelska | 99.64 | 97.53 | 17.6309 | 17.7944 |

| | | | | | | | | | | | | | | | |
|---|---|---|---|---|---|---|---|---|---|---|---|---|---|---|---|
| MZLU107463 | MZLU107463 | *Cyaniris semiargus* | Modern | Field Caught | M | 2021-07-03 | E Skåne | Fästan | 55.776 | 14.1722 | ZJ Nolen; P Jamelska | 99.58 | 97.6 | 18.1059 | 18.2741 |
| MZLU107466 | MZLU107466 | *Cyaniris semiargus* | Modern | Field Caught | M | 2021-07-03 | E Skåne | Fästan | 55.776 | 14.1722 | ZJ Nolen; P Jamelska | 99.64 | 97.58 | 19.5090 | 19.6931 |
| MZLU107503 | MZLU107503 | *Cyaniris semiargus* | Modern | Field Caught | M | 2021-07-03 | E Skåne | Fästan | 55.776 | 14.1722 | ZJ Nolen; P Jamelska | 99.51 | 97.64 | 9.0339 | 9.1190 |
| MZLU107504 | MZLU107504 | *Cyaniris semiargus* | Modern | Field Caught | M | 2021-07-03 | E Skåne | Fästan | 55.776 | 14.1722 | ZJ Nolen; P Jamelska | 99.66 | 97.73 | 14.3537 | 14.4856 |
| MZLU107506 | MZLU107506 | *Cyaniris semiargus* | Modern | Field Caught | M | 2021-07-03 | E Skåne | Fästan | 55.776 | 14.1722 | ZJ Nolen; P Jamelska | 99.53 | 97.55 | 13.2708 | 13.3974 |
| MZLU107507 | MZLU107507 | *Cyaniris semiargus* | Modern | Field Caught | M | 2021-07-03 | E Skåne | Fästan | 55.776 | 14.1722 | ZJ Nolen; P Jamelska | 99.62 | 97.62 | 14.4395 | 14.5770 |
| CYSE0078 | MZLU00107520 | *Cyaniris semiargus* | Modern | Field Caught | M | 2022-06-23 | SE Skåne | SE Skåne | 55.3864 | 14.1014 | Melina Eberhagen | 99.64 | 98.09 | 15.2662 | 15.3476 |
| CYSE0094 | MZLU00107522 | *Cyaniris semiargus* | Modern | Field Caught | M | 2022-06-27 | SE Skåne | SE Skåne | 55.3864 | 14.1014 | Melina Eberhagen | 97.84 | 96.82 | 12.8535 | 12.9280 |
| CYSE0096 | MZLU00107523 | *Cyaniris semiargus* | Modern | Field Caught | M | 2022-06-27 | SE Skåne | SE Skåne | 55.3864 | 14.1014 | Melina Eberhagen | 99.22 | 97.76 | 7.3142 | 7.3457 |
| CYSE0098 | MZLU00107524 | *Cyaniris semiargus* | Modern | Field Caught | M | 2022-06-27 | SE Skåne | SE Skåne | 55.3864 | 14.1014 | Melina Eberhagen | 99.64 | 97.99 | 12.7827 | 12.8535 |
| CYSE0116 | MZLU00107525 | *Cyaniris semiargus* | Modern | Field Caught | M | 2022-06-28 | SE Skåne | SE Skåne | 55.3864 | 14.1014 | Melina Eberhagen | 99.62 | 98.14 | 10.8479 | 10.9059 |
| CYSE0198 | MZLU00107526 | *Cyaniris semiargus* | Modern | Field Caught | F | 2022-07-01 | SE Skåne | SE Skåne | 55.3864 | 14.1014 | Melina Eberhagen | 99.48 | 97.78 | 13.4596 | 13.5280 |
| CYSE0199 | MZLU00107527 | *Cyaniris semiargus* | Modern | Field Caught | F | 2022-07-01 | SE Skåne | SE Skåne | 55.3864 | 14.1014 | Melina Eberhagen | 99.44 | 97.81 | 10.7599 | 10.8094 |
| MZLU107461 | MZLU107461 | *Cyaniris semiargus* | Modern | Field Caught | F | 2021-06-24 | Småland | Götafors | 57.492 | 14.1134 | ZJ Nolen; P Jamelska | 99.5 | 97.27 | 15.5624 | 15.6980 |
| MZLU107462 | MZLU107462 | *Cyaniris semiargus* | Modern | Field Caught | F | 2021-06-24 | Småland | Götafors | 57.492 | 14.1134 | ZJ Nolen; P Jamelska | 99.55 | 97.55 | 20.3960 | 20.5832 |
| MZLU107465 | MZLU107465 | *Cyaniris semiargus* | Modern | Field Caught | M | 2021-06-24 | Småland | Götafors | 57.492 | 14.1134 | ZJ Nolen; P Jamelska | 99.67 | 97.75 | 24.9321 | 25.1610 |
| MZLU107455 | MZLU107455 | *Cyaniris semiargus* | Modern | Field Caught | M | 2021-07-07 | Småland | Götafors | 57.492 | 14.1134 | ZJ Nolen; P Jamelska | 99.66 | 97.61 | 13.2588 | 13.3896 |
| MZLU107456 | MZLU107456 | *Cyaniris semiargus* | Modern | Field Caught | M | 2021-07-07 | Småland | Götafors | 57.492 | 14.1134 | ZJ Nolen; P Jamelska | 99.64 | 97.58 | 11.1263 | 11.2295 |
| MZLU107501 | MZLU107501 | *Cyaniris semiargus* | Modern | Field Caught | F | 2021-07-07 | Småland | Götafors | 57.492 | 14.1134 | ZJ Nolen; P Jamelska | 99.52 | 97.26 | 10.9493 | 11.0538 |
| MZLU107457 | MZLU107457 | *Cyaniris semiargus* | Modern | Field Caught | M | 2021-07-04 | W Skåne | Silvåkra | 55.6695 | 13.4924 | ZJ Nolen; P Jamelska | 99.66 | 97.71 | 19.0891 | 19.2644 |
| MZLU107460 | MZLU107460 | *Cyaniris semiargus* | Modern | Field Caught | M | 2021-07-04 | W Skåne | Silvåkra | 55.6695 | 13.4924 | ZJ Nolen; P Jamelska | 99.64 | 97.71 | 15.4684 | 15.6090 |
| MZLU107464 | MZLU107464 | *Cyaniris semiargus* | Modern | Field Caught | M | 2021-07-04 | W Skåne | Silvåkra | 55.6695 | 13.4924 | ZJ Nolen; P Jamelska | 99.55 | 97.62 | 15.2425 | 15.3885 |
| MZLU107497 | MZLU107497 | *Cyaniris semiargus* | Modern | Field Caught | M | 2021-07-04 | W Skåne | Silvåkra | 55.6695 | 13.4924 | ZJ Nolen; P Jamelska | 94.21 | 93.41 | 11.5859 | 11.6982 |
| MZLU107498 | MZLU107498 | *Cyaniris semiargus* | Modern | Field Caught | M | 2021-07-04 | W Skåne | Silvåkra | 55.6695 | 13.4924 | ZJ Nolen; P Jamelska | 99.37 | 97.37 | 12.1112 | 12.2275 |
| MZLU107499 | MZLU107499 | *Cyaniris semiargus* | Modern | Field Caught | M | 2021-07-04 | W Skåne | Silvåkra | 55.6695 | 13.4924 | ZJ Nolen; P Jamelska | 99.66 | 97.8 | 12.8861 | 13.0066 |
| MZLU107500 | MZLU107500 | *Cyaniris semiargus* | Modern | Field Caught | M | 2021-07-04 | W Skåne | Silvåkra | 55.6695 | 13.4924 | ZJ Nolen; P Jamelska | 99.56 | 97.31 | 9.5916 | 9.6883 |
| MZLU107505 | MZLU107505 | *Cyaniris semiargus* | Modern | Field Caught | M | 2021-07-04 | W Skåne | Silvåkra | 55.6695 | 13.4924 | ZJ Nolen; P Jamelska | 99.13 | 97.1 | 12.6073 | 12.7307 |
| MZLU166853 | MZLU166853 | *Agriades optilete* | Historical | Pinned Specimen (abdomen) | M | 1961-06-29 | Småland | Agunnaryd | 56.7318 | 14.2439 | Nils Burrau | NA | 100 | NA | 7.7865 |
| MZLU107492 | MZLU107492 | *Agriades optilete* | Modern | Field Caught | M | 2021-07-05 | Småland | Agunnaryd | 56.7318 | 14.2439 | ZJ Nolen | NA | 98.23 | NA | 12.211 |
| MZLU166851 | MZLU166851 | *Aricia agestis* | Historical | Pinned Specimen (abdomen) | M | 1939-xx-xx | W Skåne | Vomb | 55.6731 | 13.5503 | Nils Burrau | NA | 100 | NA | 5.8508 |
| MZLU107489 | MZLU107489 | *Aricia agestis* | Modern | Field Caught | M | 2020-08-03 | W Skåne | Tvedöra | 55.6959 | 13.431 | ZJ Nolen | NA | 94.13 | NA | 7.5821 |
| MZLU166852 | MZLU166852 | *Cupido minimus* | Historical | Pinned Specimen (abdomen) | M | 1943-05-30 | W Skåne | Torna Hällestad | 55.6781 | 13.4214 | Nils Burrau | NA | 99.99 | NA | 6.2958 |
| CUMI0012 | NA | *Cupido minimus* | Modern | Field Caught | M | 2022-06-14 | W Skåne | Tvedöra | 55.6959 | 13.431 | ZJ Nolen | NA | 96.21 | NA | 7.7405 |
| EK001 | MZLU161410 | *Plebejus argyrognomon* | Historical | Pinned Specimen (abdomen) | M | 1967-07-18 | W Skåne | Dalhem | 56.0675 | 12.7367 | Harry Rydén | NA | 99.97 | NA | 9.6382 |
| EK006 | private collection | *Plebejus argyrognomon* | Modern | Pinned Specimen (legs) | M | 2003-07-18 | W Skåne | Dalhem | 56.0675 | 12.7367 | Göran Engqvist | NA | 99.99 | NA | 13.8387 |
| MZLU174142 | MZLU174142 | *Scolitantides orion* | Historical | Pinned Specimen (abdomen) | M | 1944-xx-08 | Södermanland | Stockholm | 59.3294 | 18.0686 | E. Norstrand | NA | 100 | NA | 7.779 |
| MZLU174140 | MZLU174140 | *Scolitantides orion* | Modern | Pinned Specimen (abdomen) | M | 2016-xx-xx | Södermanland | Långaedebergen | 59.2154 | 17.1414 | Håkan Elmqvist | NA | 100 | NA | 8.4804 |

**Table S2. Individual estimates of heterozygosity and inbreeding (Froh).** Individual heterozygosity was estimated from the filtered genotypes, called with data subsampled to a uniform 6x coverage. We estimate heterozygosity as a count of heterozygous genotypes per 1000 genotypes, as well as a count per alternate allele as a potential correction for reference biases. These two estimates largely agreed with each other. Froh is the inbreeding coefficient from runs of homozygosity, estimated as the proportion of the autosomal genome in runs of homozygosity longer than 100kb. This analysis was performed using BCFtools RoH on called genotypes. For both analyses, transition sites were removed. For heterozygosity this means that the values are lower than true heterozygosity in the samples, instead representing only transversion heterozygous sites per 1000bp. Froh values should not be affected by transition removal in this same way, as it is based on ranges across multiple variants, not variant counts. Finally, we include an estimate of heterozygosity adjusted by Froh to approximate heterozygosity outside of runs of homozygosity. Much of the variation in heterozygosity between individuals is explained by variation in Froh, as can be seen by the reduced variance in this statistic.

| Sample ID | Species | Time Period | Study Region | Heterozygous sites / 1000bp (excluding transitions; 6x depth) | Heterozygous sites / called alternate allele (excluding transitions; 6x depth) | Froh > 100kb (6x depth) | Heterozygosity / (1-Froh) |
|---|---|---|---|---|---|---|---|
| MZLULEP1724 | *Polyommatus icarus* | Historical | E Skåne | 3.733 | 0.467 | 0.000 | 3.733 |
| MZLULEP1661 | *Polyommatus icarus* | Historical | E Skåne | 3.732 | 0.469 | 0.001 | 3.734 |
| MZLULEP1903 | *Polyommatus icarus* | Historical | E Skåne | 3.731 | 0.469 | 0.000 | 3.731 |
| MZLULEP3086 | *Polyommatus icarus* | Historical | Öland | 3.728 | 0.464 | 0.000 | 3.730 |
| MZLULEP3149 | *Polyommatus icarus* | Historical | Öland | 3.702 | 0.464 | 0.002 | 3.709 |
| MZLULEP3328 | *Polyommatus icarus* | Historical | Öland | 3.744 | 0.465 | 0.004 | 3.757 |
| MZLULEP3029 | *Polyommatus icarus* | Historical | Öland | 3.728 | 0.465 | 0.002 | 3.735 |
| MZLULEP3256 | *Polyommatus icarus* | Historical | Öland | 3.656 | 0.463 | 0.001 | 3.659 |
| MZLULEP3266 | *Polyommatus icarus* | Historical | Öland | 3.819 | 0.469 | 0.000 | 3.821 |
| MZLULEP3313 | *Polyommatus icarus* | Historical | Öland | 3.632 | 0.458 | 0.002 | 3.640 |
| MZLULEP3088 | *Polyommatus icarus* | Historical | Öland | 3.973 | 0.471 | 0.004 | 3.988 |
| MZLULEP3679 | *Polyommatus icarus* | Historical | W Skåne | 3.690 | 0.465 | 0.002 | 3.698 |
| MZLULEP4538 | *Polyommatus icarus* | Historical | W Skåne | 3.602 | 0.459 | 0.000 | 3.603 |
| MZLULEP1710 | *Polyommatus icarus* | Historical | W Skåne | 3.938 | 0.471 | 0.001 | 3.940 |
| MZLULEP1709 | *Polyommatus icarus* | Historical | W Skåne | 3.873 | 0.472 | 0.000 | 3.873 |
| MZLULEP3656 | *Polyommatus icarus* | Historical | W Skåne | 3.815 | 0.471 | 0.002 | 3.821 |
| PI200 | *Polyommatus icarus* | Modern | E Skåne | 3.781 | 0.470 | 0.001 | 3.783 |
| PI201 | *Polyommatus icarus* | Modern | E Skåne | 3.808 | 0.472 | 0.000 | 3.809 |
| PI202 | *Polyommatus icarus* | Modern | E Skåne | 3.782 | 0.471 | 0.000 | 3.784 |
| PI203 | *Polyommatus icarus* | Modern | E Skåne | 3.764 | 0.469 | 0.001 | 3.767 |
| PI204 | *Polyommatus icarus* | Modern | E Skåne | 3.775 | 0.471 | 0.001 | 3.778 |
| PI205 | *Polyommatus icarus* | Modern | E Skåne | 3.782 | 0.471 | 0.000 | 3.782 |
| PI206 | *Polyommatus icarus* | Modern | E Skåne | 3.768 | 0.470 | 0.000 | 3.768 |
| PI207 | *Polyommatus icarus* | Modern | E Skåne | 3.789 | 0.470 | 0.000 | 3.789 |
| PI208 | *Polyommatus icarus* | Modern | E Skåne | 3.777 | 0.470 | 0.000 | 3.777 |
| PI413 | *Polyommatus icarus* | Modern | Öland | 3.741 | 0.465 | 0.001 | 3.743 |
| PI414 | *Polyommatus icarus* | Modern | Öland | 3.718 | 0.463 | 0.000 | 3.718 |
| PI415 | *Polyommatus icarus* | Modern | Öland | 3.753 | 0.466 | 0.001 | 3.755 |
| PI417 | *Polyommatus icarus* | Modern | Öland | 3.729 | 0.465 | 0.001 | 3.734 |
| PI418 | *Polyommatus icarus* | Modern | Öland | 3.728 | 0.465 | 0.000 | 3.728 |
| PI419 | *Polyommatus icarus* | Modern | Öland | 3.697 | 0.462 | 0.000 | 3.697 |
| PI420 | *Polyommatus icarus* | Modern | Öland | 3.707 | 0.463 | 0.002 | 3.716 |
| PI421 | *Polyommatus icarus* | Modern | Öland | 3.728 | 0.463 | 0.002 | 3.735 |
| PI422 | *Polyommatus icarus* | Modern | Öland | 3.722 | 0.465 | 0.001 | 3.724 |
| PI028 | *Polyommatus icarus* | Modern | W Skåne | 3.783 | 0.471 | 0.001 | 3.787 |
| PI029 | *Polyommatus icarus* | Modern | W Skåne | 3.765 | 0.469 | 0.001 | 3.769 |
| PI030 | *Polyommatus icarus* | Modern | W Skåne | 3.776 | 0.472 | 0.000 | 3.776 |
| PI031 | *Polyommatus icarus* | Modern | W Skåne | 3.815 | 0.472 | 0.008 | 3.846 |
| PI032 | *Polyommatus icarus* | Modern | W Skåne | 3.755 | 0.469 | 0.000 | 3.755 |
| PI033 | *Polyommatus icarus* | Modern | W Skåne | 3.777 | 0.469 | 0.015 | 3.834 |
| PI034 | *Polyommatus icarus* | Modern | W Skåne | 3.774 | 0.470 | 0.000 | 3.774 |
| PI035 | *Polyommatus icarus* | Modern | W Skåne | 3.783 | 0.471 | 0.000 | 3.783 |
| PI036 | *Polyommatus icarus* | Modern | W Skåne | 3.758 | 0.468 | 0.001 | 3.760 |

| | | | | | | | |
|---|---|---|---|---|---|---|---|
| MZLU153003 | *Plebejus argus* | Historical | E Skåne | 2.623 | 0.448 | 0.001 | 2.627 |
| MZLU153166 | *Plebejus argus* | Historical | Öland | 2.606 | 0.445 | 0.003 | 2.613 |
| MZLU153167 | *Plebejus argus* | Historical | Öland | 2.566 | 0.442 | 0.004 | 2.576 |
| MZLU153168 | *Plebejus argus* | Historical | Öland | 2.590 | 0.441 | 0.006 | 2.606 |
| MZLU153169 | *Plebejus argus* | Historical | Öland | 2.591 | 0.445 | 0.003 | 2.597 |
| MZLU153170 | *Plebejus argus* | Historical | Öland | 2.585 | 0.443 | 0.008 | 2.604 |
| MZLU153046 | *Plebejus argus* | Historical | Småland | 2.593 | 0.447 | 0.000 | 2.593 |
| MZLU153042 | *Plebejus argus* | Historical | Småland | 2.552 | 0.436 | 0.006 | 2.568 |
| MZLU153048 | *Plebejus argus* | Historical | Småland | 2.522 | 0.431 | 0.031 | 2.602 |
| MZLU152940 | *Plebejus argus* | Historical | W Skåne | 2.564 | 0.440 | 0.017 | 2.607 |
| PLAR0200 | *Plebejus argus* | Modern | E Skåne | 2.619 | 0.444 | 0.005 | 2.633 |
| PLAR0202 | *Plebejus argus* | Modern | E Skåne | 2.629 | 0.447 | 0.000 | 2.629 |
| PLAR0203 | *Plebejus argus* | Modern | E Skåne | 2.623 | 0.444 | 0.001 | 2.624 |
| PLAR0208 | *Plebejus argus* | Modern | E Skåne | 2.626 | 0.445 | 0.002 | 2.630 |
| PLAR0211 | *Plebejus argus* | Modern | E Skåne | 2.580 | 0.439 | 0.012 | 2.613 |
| PLAR0217 | *Plebejus argus* | Modern | E Skåne | 2.625 | 0.445 | 0.002 | 2.630 |
| PLAR0222 | *Plebejus argus* | Modern | E Skåne | 2.612 | 0.443 | 0.006 | 2.628 |
| PLAR0226 | *Plebejus argus* | Modern | E Skåne | 2.586 | 0.446 | 0.002 | 2.590 |
| MZLU107434 | *Plebejus argus* | Modern | Öland | 2.577 | 0.439 | 0.009 | 2.599 |
| MZLU107438 | *Plebejus argus* | Modern | Öland | 2.572 | 0.437 | 0.012 | 2.604 |
| MZLU107435 | *Plebejus argus* | Modern | Öland | 2.593 | 0.440 | 0.007 | 2.612 |
| MZLU107436 | *Plebejus argus* | Modern | Öland | 2.491 | 0.423 | 0.041 | 2.597 |
| MZLU107445 | *Plebejus argus* | Modern | Öland | 2.504 | 0.429 | 0.028 | 2.577 |
| MZLU107449 | *Plebejus argus* | Modern | Öland | 2.558 | 0.435 | 0.013 | 2.591 |
| MZLU107451 | *Plebejus argus* | Modern | Öland | 2.590 | 0.442 | 0.001 | 2.593 |
| MZLU107471 | *Plebejus argus* | Modern | Öland | 2.558 | 0.435 | 0.012 | 2.589 |
| MZLU107477 | *Plebejus argus* | Modern | Öland | 2.574 | 0.438 | 0.002 | 2.580 |
| MZLU107487 | *Plebejus argus* | Modern | Öland | 2.497 | 0.425 | 0.030 | 2.573 |
| MZLU107470 | *Plebejus argus* | Modern | Småland | 2.594 | 0.441 | 0.001 | 2.597 |
| MZLU107486 | *Plebejus argus* | Modern | Småland | 2.558 | 0.435 | 0.019 | 2.608 |
| MZLU107469 | *Plebejus argus* | Modern | Småland | 2.582 | 0.440 | 0.000 | 2.582 |
| MZLU107483 | *Plebejus argus* | Modern | Småland | 2.571 | 0.437 | 0.011 | 2.601 |
| MZLU107426 | *Plebejus argus* | Modern | SW Skåne | 2.600 | 0.440 | 0.010 | 2.626 |
| MZLU107439 | *Plebejus argus* | Modern | SW Skåne | 2.584 | 0.438 | 0.022 | 2.642 |
| MZLU107441 | *Plebejus argus* | Modern | SW Skåne | 2.565 | 0.435 | 0.017 | 2.611 |
| MZLU107443 | *Plebejus argus* | Modern | SW Skåne | 2.584 | 0.438 | 0.010 | 2.611 |
| MZLU107468 | *Plebejus argus* | Modern | SW Skåne | 2.606 | 0.440 | 0.008 | 2.626 |
| MZLU107475 | *Plebejus argus* | Modern | SW Skåne | 2.616 | 0.444 | 0.001 | 2.618 |
| MZLU107480 | *Plebejus argus* | Modern | SW Skåne | 2.602 | 0.441 | 0.008 | 2.624 |
| MZLU153246 | *Cyaniris semiargus* | Historical | E Skåne | 2.283 | 0.434 | 0.004 | 2.293 |
| MZLU153247 | *Cyaniris semiargus* | Historical | E Skåne | 2.299 | 0.432 | 0.003 | 2.307 |
| MZLU153248 | *Cyaniris semiargus* | Historical | E Skåne | 2.282 | 0.432 | 0.004 | 2.292 |
| MZLU153251 | *Cyaniris semiargus* | Historical | E Skåne | 2.221 | 0.417 | 0.031 | 2.291 |
| MZLU153250 | *Cyaniris semiargus* | Historical | E Skåne | 2.291 | 0.435 | 0.000 | 2.291 |
| MZLU152770 | *Cyaniris semiargus* | Historical | Småland | 2.067 | 0.398 | 0.014 | 2.097 |
| MZLU152769 | *Cyaniris semiargus* | Historical | Småland | 2.150 | 0.409 | 0.003 | 2.156 |
| MZLU152766 | *Cyaniris semiargus* | Historical | Småland | 2.065 | 0.406 | 0.001 | 2.067 |
| MZLU153204 | *Cyaniris semiargus* | Historical | W Skåne | 2.208 | 0.421 | 0.026 | 2.268 |
| MZLU153205 | *Cyaniris semiargus* | Historical | W Skåne | 2.283 | 0.434 | 0.004 | 2.291 |
| MZLU153206 | *Cyaniris semiargus* | Historical | W Skåne | 2.248 | 0.430 | 0.009 | 2.270 |
| MZLU153216 | *Cyaniris semiargus* | Historical | W Skåne | 2.282 | 0.434 | 0.000 | 2.282 |
| MZLU153221 | *Cyaniris semiargus* | Historical | W Skåne | 2.224 | 0.429 | 0.002 | 2.228 |
| MZLU107454 | *Cyaniris semiargus* | Modern | E Skåne | 2.234 | 0.425 | 0.008 | 2.251 |
| MZLU107458 | *Cyaniris semiargus* | Modern | E Skåne | 2.197 | 0.418 | 0.022 | 2.247 |
| MZLU107463 | *Cyaniris semiargus* | Modern | E Skåne | 2.195 | 0.417 | 0.033 | 2.270 |
| MZLU107466 | *Cyaniris semiargus* | Modern | E Skåne | 2.238 | 0.426 | 0.006 | 2.253 |

| | | | | | | | |
|---|---|---|---|---|---|---|---|
| MZLU107503 | *Cyaniris semiargus* | Modern | E Skåne | 2.229 | 0.423 | 0.023 | 2.283 |
| MZLU107504 | *Cyaniris semiargus* | Modern | E Skåne | 2.236 | 0.426 | 0.015 | 2.270 |
| MZLU107506 | *Cyaniris semiargus* | Modern | E Skåne | 2.226 | 0.424 | 0.014 | 2.258 |
| MZLU107507 | *Cyaniris semiargus* | Modern | E Skåne | 2.237 | 0.425 | 0.008 | 2.256 |
| CYSE0078 | *Cyaniris semiargus* | Modern | SE Skåne | 1.533 | 0.284 | 0.329 | 2.286 |
| CYSE0094 | *Cyaniris semiargus* | Modern | SE Skåne | 1.852 | 0.347 | 0.186 | 2.275 |
| CYSE0096 | *Cyaniris semiargus* | Modern | SE Skåne | 2.013 | 0.378 | 0.118 | 2.283 |
| CYSE0098 | *Cyaniris semiargus* | Modern | SE Skåne | 1.962 | 0.367 | 0.143 | 2.288 |
| CYSE0116 | *Cyaniris semiargus* | Modern | SE Skåne | 2.062 | 0.389 | 0.092 | 2.271 |
| CYSE0198 | *Cyaniris semiargus* | Modern | SE Skåne | 2.104 | 0.396 | 0.075 | 2.274 |
| CYSE0199 | *Cyaniris semiargus* | Modern | SE Skåne | 1.963 | 0.368 | 0.133 | 2.264 |
| MZLU107461 | *Cyaniris semiargus* | Modern | Småland | 2.160 | 0.408 | 0.018 | 2.198 |
| MZLU107462 | *Cyaniris semiargus* | Modern | Småland | 2.162 | 0.412 | 0.001 | 2.164 |
| MZLU107465 | *Cyaniris semiargus* | Modern | Småland | 1.806 | 0.339 | 0.170 | 2.176 |
| MZLU107455 | *Cyaniris semiargus* | Modern | Småland | 2.177 | 0.412 | 0.003 | 2.183 |
| MZLU107456 | *Cyaniris semiargus* | Modern | Småland | 1.821 | 0.344 | 0.147 | 2.134 |
| MZLU107501 | *Cyaniris semiargus* | Modern | Småland | 2.008 | 0.378 | 0.072 | 2.165 |
| MZLU107457 | *Cyaniris semiargus* | Modern | W Skåne | 2.137 | 0.403 | 0.060 | 2.274 |
| MZLU107460 | *Cyaniris semiargus* | Modern | W Skåne | 2.159 | 0.408 | 0.058 | 2.291 |
| MZLU107464 | *Cyaniris semiargus* | Modern | W Skåne | 2.206 | 0.417 | 0.029 | 2.273 |
| MZLU107497 | *Cyaniris semiargus* | Modern | W Skåne | 2.109 | 0.397 | 0.075 | 2.279 |
| MZLU107498 | *Cyaniris semiargus* | Modern | W Skåne | 2.206 | 0.417 | 0.036 | 2.288 |
| MZLU107499 | *Cyaniris semiargus* | Modern | W Skåne | 2.039 | 0.382 | 0.119 | 2.315 |
| MZLU107500 | *Cyaniris semiargus* | Modern | W Skåne | 2.141 | 0.402 | 0.067 | 2.295 |
| MZLU107505 | *Cyaniris semiargus* | Modern | W Skåne | 2.179 | 0.412 | 0.047 | 2.287 |
| MZLU166853 | *Agriades optilete* | Historical | Småland | 0.790 | 0.342 | 0.005 | 0.794 |
| MZLU107492 | *Agriades optilete* | Modern | Småland | 0.796 | 0.343 | 0.002 | 0.797 |
| MZLU166851 | *Aricia agestis* | Historical | W Skåne | 2.507 | 0.331 | 0.107 | 2.806 |
| MZLU107489 | *Aricia agestis* | Modern | W Skåne | 2.691 | 0.348 | 0.080 | 2.925 |
| MZLU166852 | *Cupido minimus* | Historical | W Skåne | 2.225 | 0.354 | 0.068 | 2.387 |
| CUMI0012 | *Cupido minimus* | Modern | W Skåne | 2.324 | 0.360 | 0.090 | 2.552 |
| EK001 | *Plebejus argyrognomon* | Historical | W Skåne | 1.559 | 0.057 | 0.038 | 1.621 |
| EK006 | *Plebejus argyrognomon* | Modern | W Skåne | 1.461 | 0.053 | 0.082 | 1.591 |
| MZLU174142 | *Scolitantides orion* | Historical | Södermanland | 0.275 | 0.053 | 0.173 | 0.333 |
| MZLU174140 | *Scolitantides orion* | Modern | Södermanland | 0.176 | 0.034 | 0.407 | 0.297 |

**Table S3. Statistical testing of differences in heterozygosity between modern and historical samples.** We statistically assessed differences in heterozygosity between modern and historical samples for each species using a linear mixed model with heterozygosity as a response to the fixed effect of time period and the random effect of sampling region. Outputs from the models are summarized here.

| Species | Mean heterozyg. (hist) | Mean heterozyg. (mod) | Heterozyg. mod/hist | Difference est. | Std.Error | t-statistic | 95% CI (lower) | 95% CI (upper) | p-value |
|---|---|---|---|---|---|---|---|---|---|
| Po. icarus | 3.756 | 3.7606 | 1.00123 | 0.0019 | 0.0209 | 0.0907 | -0.04038 | 0.04417 | 0.9282 |
| Pl. argus | 2.57904 | 2.58191 | 1.00111 | -0.0083 | 0.0109 | -0.76148 | -0.03045 | 0.01385 | 0.45162 |
| Cy. semiargus | 2.22325 | 2.08933 | 0.93976 | -0.13302 | 0.04753 | -2.79842 | -0.22925 | -0.03679 | **0.00802** |

**Table S4. Estimates of pairwise nucleotide diversity for populations of focal species.** For each focal species population sample, we estimated nucleotide diversity (pi) in 50kb non-overlapping sliding windows using the method implemented in Pixy. We estimated mean nucleotide diversity as an average of all windows with confidence intervals estimated by performing 10 000 bootstraps, resampling windows with replacement.

| Species | Region | Time Period | Year | Mean Nucleotide Diversity | 95% CI (Lower) | 95% CI (Upper) |
|---|---|---|---|---|---|---|
| *Po. icarus* | E Skåne | Historical | 1934 | 0.003891 | 0.003868 | 0.003915 |
| *Po. icarus* | E Skåne | Modern | 2020 | 0.003960 | 0.003939 | 0.003982 |
| *Po. icarus* | Öland | Historical | 1940 | 0.003947 | 0.003925 | 0.003969 |
| *Po. icarus* | Öland | Modern | 2020 | 0.003923 | 0.003902 | 0.003945 |
| *Po. icarus* | W Skåne | Historical | 1934 | 0.003999 | 0.003976 | 0.004021 |
| *Po. icarus* | W Skåne | Modern | 2020 | 0.003953 | 0.003930 | 0.003976 |
| *Pl. argus* | E Skåne | Modern | 2022 | 0.002771 | 0.002750 | 0.002793 |
| *Pl. argus* | Öland | Historical | 1951 | 0.002758 | 0.002735 | 0.002781 |
| *Pl. argus* | Öland | Modern | 2021 | 0.002716 | 0.002694 | 0.002738 |
| *Pl. argus* | Småland | Historical | 1936 | 0.002750 | 0.002727 | 0.002773 |
| *Pl. argus* | Småland | Modern | 2021 | 0.002711 | 0.002688 | 0.002733 |
| *Pl. argus* | SW Skåne | Modern | 2021 | 0.002765 | 0.002742 | 0.002787 |
| *Cy. semiargus* | E Skåne | Historical | 1956 | 0.002389 | 0.002373 | 0.002405 |
| *Cy. semiargus* | E Skåne | Modern | 2021 | 0.002345 | 0.002322 | 0.002377 |
| *Cy. semiargus* | SE Skåne | Modern | 2022 | 0.002091 | 0.002069 | 0.002118 |
| *Cy. semiargus* | Småland | Historical | 1936 | 0.002191 | 0.002175 | 0.002208 |
| *Cy. semiargus* | Småland | Modern | 2021 | 0.002175 | 0.002149 | 0.002209 |
| *Cy. semiargus* | W Skåne | Historical | 1951 | 0.002300 | 0.002280 | 0.002324 |
| *Cy. semiargus* | W Skåne | Modern | 2021 | 0.002252 | 0.002236 | 0.002268 |

**Table S5. Change in nucleotide diversity over the sampled time period and projected to 100 years.** Only comparisons between geographically proximal (<40km) sampling sites across time periods are shown. Proportion nucleotide diversity retained represents the modern population nucleotide diversity over the historical population nucleotide diversity. This proportion is extrapolated to a yearly rate of change which is then standardized to 100 years and placed into a threat category using the indicators outlined by Andersson et al. (21).

| Species | Modern Region | Modern Year | Historical Region | Historical Year | Proportion retained | Proportion retained / year | % retained per 100 years | Threat category |
|---|---|---|---|---|---|---|---|---|
| *P. icarus* | E Skåne | 2020 | E Skåne | 1934 | 1.0177 | 1.0002 | 102.06 | Acceptable |
| *P. icarus* | Öland | 2020 | Öland | 1940 | 0.9940 | 0.9999 | 99.25 | Acceptable |
| *P. icarus* | W Skåne | 2020 | W Skåne | 1934 | 0.9885 | 0.9999 | 98.66 | Acceptable |
| *P. argus* | Öland | 2021 | Öland | 1951 | 0.9848 | 0.9998 | 97.84 | Acceptable |
| *P. argus* | Småland | 2021 | Småland | 1936 | 0.9858 | 0.9998 | 98.33 | Acceptable |
| *C. semiargus* | E Skåne | 2021 | E Skåne | 1956 | 0.9817 | 0.9997 | 97.20 | Acceptable |
| *C. semiargus* | SE Skåne | 2022 | E Skåne | 1956 | 0.8752 | 0.9980 | 81.71 | Warning |
| *C. semiargus* | Småland | 2021 | Småland | 1936 | 0.9924 | 0.9999 | 99.10 | Acceptable |
| *C. semiargus* | W Skåne | 2021 | W Skåne | 1951 | 0.9792 | 0.9997 | 97.04 | Acceptable |

**Table S6. Predicted retention of heterozygosity over 100 years per species and estimates of contemporary effective population size.** We estimated the predicted percent of heterozygosity retained over 100 years, following Andersson et al. (21) for each of the study species. For the focal species where we had multiple samples per populations, we first estimated it using the mean population heterozygosity, inferring a yearly rate of heterozygosity retention per population and projecting it out to 100 years. As the additional species only included one sample per time period, we estimated this rate with all possible historical and modern individual pairs for a region. For supplementray species, this results in a single number, for the three focal species, we report the median and range values. We assigned a threat category to each of the median estimates, based on the thresholds described by Andersson et al. For the focal species, we estimated recent trajectories of effective population size with GONE, reporting the size at generation 5 here and assessing this value under the indicator from Andersson et al.

| | | | *Population means* | | *Pairwise between individuals* | | *Contemporary Ne* | |
|---|---|---|---|---|---|---|---|---|
| **Species** | **Study Region** | **2020 Swedish Red List** | **% heterozygosity retained over 100 years** | **Threat Categorization** | **Median % heterozygosity retained over 100 years (range)** | **Threat Categorization** | **Effective population size** | **Threat Categorization** |
| *Aricia agestis* | W Skåne | LC | NA | NA | 109.02 | Acceptable | NA | NA |
| *Agriades optilete* | Småland | LC | NA | NA | 101.22 | Acceptable | NA | NA |
| *Cupido minimus* | W Skåne | NT | NA | NA | 105.64 | Acceptable | NA | NA |
| *Cyaniris semiargus* | E Skåne | LC | 96.58 | Acceptable | 96.36 (93.17-101.22) | Acceptable | 520.557 | Acceptable |
| *Cyaniris semiargus* | E Skåne (SE) | LC | 77.76 | Warning | 79.64 (54.13-92.15) | Warning | 35.4456 | Alarm |
| *Cyaniris semiargus* | Småland | LC | 95.98 | Acceptable | 98.65 (81.46-106.42) | Acceptable | 1338360 | Acceptable |
| *Cyaniris semiargus* | W Skåne | LC | 93.57 | Warning | 93.99 (85.13-99.87) | Warning | 201.655 | Warning |
| *Plebejus argus* | E Skåne | LC | 99.61 | Acceptable | 99.91 (98.45-100.2) | Acceptable | 3621410 | Acceptable |
| *Plebejus argus* | Öland | LC | 97.76 | Acceptable | 98.21 (94.08-101.34) | Acceptable | 1303.89 | Acceptable |
| *Plebejus argus* | Småland | LC | 100.54 | Acceptable | 100.89 (95.41-103.39) | Acceptable | 4297.46 | Acceptable |
| *Plebejus argus* | W Skåne | LC | 101.52 | Acceptable | 101.84 (100.08-102.65) | Acceptable | 1337.37 | Acceptable |
| *Plebejus argyrognomon* | W Skåne | CR | NA | NA | 87.89 | Warning | NA | NA |
| *Polyommatus icarus* | E Skåne | LC | 101.52 | Acceptable | 101.52 (100.97-102.4) | Acceptable | 3259430 | Acceptable |
| *Polyommatus icarus* | Öland | LC | 99.23 | Acceptable | 99.99 (91.38-104.16) | Acceptable | 3451160 | Acceptable |
| *Polyommatus icarus* | W Skåne | LC | 99.78 | Acceptable | 98.8 (94.62-106.94) | Acceptable | 4033310 | Acceptable |
| *Scolitantides orion* | Södermanland | EN | NA | NA | 53.80 | Alarm | NA | NA |

**Table S7. Genetic differentiation between populations of the focal species.** We estimated genetic differentiation using pairwise *F_* ST between populations within each time period, using the Hudson estimator as implemented in Pixy. We estimated *F_* ST in 50kb non-overlapping windows, estimating the mean across all windows and 95% confidence intervals with 10 000 bootstraps, resampling windows with replacement.

| Species | Region 1 | Region 1 Year | Region 2 | Region 2 Year | Time Period | Distance (km) | Mean Hudson Fst | 95% CI (Lower) | 95% CI (Upper) |
|---|---|---|---|---|---|---|---|---|---|
| *P. icarus* | E Skåne | 1934 | W Skåne | 1934 | Historical | 50.01 | 0.01829 | 0.01776 | 0.01887 |
| *P. icarus* | E Skåne | 1934 | Öland | 1940 | Historical | 170.71 | 0.02103 | 0.02047 | 0.02161 |
| *P. icarus* | Öland | 1940 | W Skåne | 1934 | Historical | 212.33 | 0.01547 | 0.01503 | 0.01590 |
| *P. icarus* | E Skåne | 2020 | W Skåne | 2020 | Modern | 47.65 | 0.00818 | 0.00795 | 0.00843 |
| *P. icarus* | E Skåne | 2020 | Öland | 2020 | Modern | 178.28 | 0.01241 | 0.01207 | 0.01275 |
| *P. icarus* | Öland | 2020 | W Skåne | 2020 | Modern | 221.18 | 0.01366 | 0.01330 | 0.01403 |
| *P. argus* | Småland | 1936 | Öland | 1951 | Historical | 147.3 | 0.03497 | 0.03386 | 0.03612 |
| *P. argus* | E Skåne | 2022 | SW Skåne | 2021 | Modern | 87.5 | 0.02665 | 0.02589 | 0.02743 |
| *P. argus* | Småland | 2021 | Öland | 2021 | Modern | 173.36 | 0.03760 | 0.03659 | 0.03863 |
| *P. argus* | E Skåne | 2022 | Öland | 2021 | Modern | 182.25 | 0.03199 | 0.03113 | 0.03290 |
| *P. argus* | E Skåne | 2022 | Småland | 2021 | Modern | 193.41 | 0.03176 | 0.03088 | 0.03269 |
| *P. argus* | Småland | 2021 | SW Skåne | 2021 | Modern | 244.67 | 0.04090 | 0.03989 | 0.04193 |
| *P. argus* | Öland | 2021 | SW Skåne | 2021 | Modern | 268.99 | 0.04129 | 0.04027 | 0.04236 |
| *C. semiargus* | E Skåne | 1956 | W Skåne | 1951 | Historical | 45.61 | 0.02874 | 0.02802 | 0.02948 |
| *C. semiargus* | E Skåne | 1956 | Småland | 1936 | Historical | 166.1 | 0.04307 | 0.04187 | 0.04428 |
| *C. semiargus* | Småland | 1936 | W Skåne | 1951 | Historical | 193.88 | 0.05167 | 0.05040 | 0.05299 |
| *C. semiargus* | E Skåne | 2021 | SE Skåne | 2022 | Modern | 43.61 | 0.08680 | 0.08562 | 0.08800 |
| *C. semiargus* | E Skåne | 2021 | W Skåne | 2021 | Modern | 44.34 | 0.04940 | 0.04859 | 0.05022 |
| *C. semiargus* | SE Skåne | 2022 | W Skåne | 2021 | Modern | 49.72 | 0.10919 | 0.10788 | 0.11048 |
| *C. semiargus* | E Skåne | 2021 | Småland | 2021 | Modern | 191.11 | 0.06224 | 0.06124 | 0.06328 |
| *C. semiargus* | Småland | 2021 | W Skåne | 2021 | Modern | 206.5 | 0.08387 | 0.08267 | 0.08506 |
| *C. semiargus* | Småland | 2021 | SE Skåne | 2022 | Modern | 234.47 | 0.12170 | 0.12022 | 0.12318 |

**Table S8. Convergence results for admixture analyses.** For values of K 1-8, we performed up to 100 replicate admixture analyses per species to assess convergence, performing a minimum of 20 replicates. If the three highest likelihood replicates were within 2 log-likelihood units of each other, the replicates were considered converged and no further replicates were performed. We selected the highest value of K per species that converged as the 'best fit', which are highlighted here in bold, presented with the total number of replicates required to reach convergence, as well as the difference in log-likelihood between the top 3 replicates.

| Species | K | Replicates to convergence (max 100) | Δ log-likelihood of top 3 replicates |
|---|---|---|---|
| ***Polyommatus icarus*** | **1** | **20** | **0.000** |
| *Polyommatus icarus* | 2 | 100 | 85.270 |
| *Polyommatus icarus* | 3 | 100 | 1820.431 |
| *Polyommatus icarus* | 4 | 100 | 1820.550 |
| *Polyommatus icarus* | 5 | 100 | 4206.780 |
| *Polyommatus icarus* | 6 | 100 | 2362.288 |
| *Polyommatus icarus* | 7 | 100 | 2536.728 |
| *Polyommatus icarus* | 8 | 100 | 3810.643 |
| ***Plebejus argus*** | **1** | **20** | **0.000** |
| *Plebejus argus* | 2 | 100 | 88.839 |
| *Plebejus argus* | 3 | 100 | 502.764 |
| *Plebejus argus* | 4 | 100 | 999.728 |
| *Plebejus argus* | 5 | 100 | 646.737 |
| *Plebejus argus* | 6 | 100 | 7439.148 |
| *Plebejus argus* | 7 | 100 | 2735.900 |
| *Plebejus argus* | 8 | 100 | 6233.690 |
| *Cyaniris semiargus* | 1 | 20 | 0.000 |
| *Cyaniris semiargus* | 2 | 20 | 0.013 |
| *Cyaniris semiargus* | 3 | 20 | 0.007 |
| *Cyaniris semiargus* | 4 | 20 | 0.004 |
| ***Cyaniris semiargus*** | **5** | **23** | **0.139** |
| *Cyaniris semiargus* | 6 | 100 | 1287.189 |
| *Cyaniris semiargus* | 7 | 100 | 6647.493 |
| *Cyaniris semiargus* | 8 | 100 | 4275.050 |

**Table S9. Statistical testing of differences in inbreeding between modern and historical samples.** For each species, we tested for differences in the inbreeding coefficient Froh using a linear mixed model with Froh as a response to the fixed effect of time period and the random effect of sampling region. Outputs from the models are summarized here.

| Species | Mean Froh (hist) | Mean Froh (mod) | Difference | Std.Error | t-statistic | 95% CI (lower) | 95% CI (upper) | p-value |
|---|---|---|---|---|---|---|---|---|
| *Po. icarus* | 0.00123 | 0.00133 | 0.00015 | 0.00081 | 0.17988 | -0.00149 | 0.00178 | 0.85818 |
| *Pl. argus* | 0.00784 | 0.01008 | 0.00284 | 0.00358 | 0.79254 | -0.00444 | 0.01012 | 0.43354 |
| *Cy. semiargus* | 0.00785 | 0.0727 | 0.06485 | 0.02054 | 3.15779 | 0.02328 | 0.10643 | **0.00311** |

**Table S10. Predicted retention of heterozygosity, mean modern Froh, and heterozygosity outside of RoH from coalescent simulations.** To assess if our empirical estimates of heterozygosity reduction aligned with expected patterns under inferred population declines, we simulated demographic histories for each population based on size changes inferred with GONE. Here, we document the minimum, quantile, and maximum estimates for three parameters from 100 replicate runs of each simulated history: the proportion of heterozygosity retained in the modern compared to the historical population, the mean estimate of Froh in the modern population, and the proportion of heterozygosity retained adjusted for Froh (i.e. heterozygosity / (1-Froh) estimated in each sampled simulated individual). We find that across all simulations, nearly all changes in heterozygosity over time should be reflected with equivalent changes in Froh.

| | | *Proportion heterozygosity maintained* | | | | | *Mean modern Froh* | | | | | *Proportion heterozygosity maintained outside RoH* | | | | |
|---|---|---|---|---|---|---|---|---|---|---|---|---|---|---|---|---|
| ***Species*** | ***Region*** | ***Min*** | ***25%*** | ***50%*** | ***75%*** | ***Max*** | ***Min*** | ***25%*** | ***50%*** | ***75%*** | ***Max*** | ***Min*** | ***25%*** | ***50%*** | ***75%*** | ***Max*** |
| *Po. icarus* | ESkane | 0.9846271 | 0.9969759 | 1.0004864 | 1.004772 | 1.026243 | 0.00116629 | 0.00397381 | 0.00555022 | 0.00736752 | 0.01902226 | 0.9845965 | 0.997962 | 1.0006891 | 1.003469 | 1.014335 |
| *Po. icarus* | Oland | 0.9776929 | 0.9944727 | 1.0006546 | 1.0054319 | 1.01797 | 0.00132574 | 0.00390375 | 0.00557447 | 0.00763264 | 0.03250569 | 0.9882087 | 0.995937 | 1.0004408 | 1.00357 | 1.010349 |
| *Po. icarus* | WSkane | 0.977188 | 0.9950753 | 1.0000698 | 1.0043247 | 1.020596 | 0.00047012 | 0.00376547 | 0.00523492 | 0.0079526 | 0.01756287 | 0.9883738 | 0.9969518 | 1.0011759 | 1.004197 | 1.013341 |
| *Pl. argus* | ESkane | 0.981891 | 0.994055 | 0.9996801 | 1.0041743 | 1.016116 | 0.00133602 | 0.00355266 | 0.00506823 | 0.00797046 | 0.02636784 | 0.9858546 | 0.9953685 | 0.9994493 | 1.003302 | 1.014422 |
| *Pl. argus* | NSmaland | 0.9629014 | 0.9943889 | 0.9995231 | 1.0072257 | 1.021321 | 0.00406874 | 0.00906731 | 0.01173722 | 0.01601223 | 0.04835541 | 0.9821937 | 0.9971483 | 1.0013107 | 1.00549 | 1.016984 |
| *Pl. argus* | Oland | 0.9307722 | 0.9835353 | 0.9935773 | 0.9994117 | 1.020871 | 0.00331401 | 0.00906678 | 0.01293235 | 0.02450908 | 0.07272099 | 0.9776541 | 0.99505 | 0.9994601 | 1.003431 | 1.017594 |
| *Pl. argus* | SWSkane | 0.9342644 | 0.982784 | 0.9906265 | 1.0034128 | 1.032456 | 0.00844711 | 0.01915366 | 0.0248457 | 0.03672087 | 0.08961475 | 0.9793742 | 0.9948533 | 1.0017425 | 1.0068 | 1.023883 |
| *Cy. semiargu* | ESkane | 0.9057069 | 0.9618579 | 0.9840706 | 0.998742 | 1.028271 | 0.00663231 | 0.01265876 | 0.02513782 | 0.04718959 | 0.10646456 | 0.9806111 | 0.993558 | 1.0006155 | 1.005503 | 1.024795 |
| *Cy. semiargu* | NSmaland | 0.9864114 | 0.9956354 | 1.0009246 | 1.0044686 | 1.016274 | 0.00138933 | 0.00435796 | 0.00549145 | 0.00785324 | 0.01877597 | 0.9878845 | 0.9958627 | 1.0008936 | 1.00398 | 1.013841 |
| *Cy. semiargu* | WSkane | 0.8744006 | 0.920729 | 0.9405299 | 0.9695309 | 1.005677 | 0.00891484 | 0.04040168 | 0.06875496 | 0.09008702 | 0.13671314 | 0.9835457 | 0.9957888 | 1.0006094 | 1.005003 | 1.01724 |

**Table S11. Statistical testing of differences in deleterious burden between modern and historical samples.** We estimated deleterious burden for three categories of variants: conserved sites (those in the top 1% of GERP scores across the genome), high impact variants (e.g. loss of function, signifying strongly deleterious), and moderate impact variants (e.g. missense, signifying weakly deleteriou). We additionally estimated burden at low impact (i.e. genic synonymous) and intergenic variants, both presumed neutral. For each of these categories, we estimated total deleterious burden as the count of derived allelesof the focal category per 1000 called genotypes. We estimated homozygous burden using the count of homozygous derived alleles of the focal category per 1000 called genotypes. We statistically assessed differences in heterozygosity between modern and historical samples for each species using a linear mixed model with burden estimates as a response to the fixed effect of time period and the random effect of sampling region. Outputs from the models are summarized here.

| Species | Mean derived count / 1000 GTs (Hist) | Mean derived count / 1000 GTs (Mod) | Mod/Hist | Difference | Std.Error | t-statistic | 95% CI (lower) | 95% CI (upper) | p-value | Mutation category |
|---|---|---|---|---|---|---|---|---|---|---|
| Po. icarus | 0.09378 | 0.09062 | 0.96631 | -0.00299 | 0.00306 | -0.97664 | -0.00918 | 0.0032 | 0.33477 | Conserved Homozygous |
| Po. icarus | 1.06868 | 1.07351 | 1.00452 | 0.00483 | 0.01097 | 0.44066 | -0.01735 | 0.02702 | 0.66189 | Conserved Total |
| Po. icarus | 0.04043 | 0.04106 | 1.01557 | 0.00063 | 0.00198 | 0.31751 | -0.00338 | 0.00464 | 0.75255 | High Total |
| Po. icarus | 0.00572 | 0.00564 | 0.98539 | -8.00E-05 | 0.00082 | -0.10237 | -0.00174 | 0.00157 | 0.91899 | High Homozygous |
| Po. icarus | 27.94189 | 27.81699 | 0.99553 | -0.11309 | 0.09335 | -1.21144 | -0.3019 | 0.07573 | 0.23302 | Intergenic Total |
| Po. icarus | 5.15418 | 5.0749 | 0.98462 | -0.02557 | 0.04092 | -0.62489 | -0.10835 | 0.0572 | 0.53568 | Intergenic Homozygous |
| Po. icarus | 2.31903 | 2.36566 | 1.02011 | 0.04663 | 0.02664 | 1.75008 | -0.00726 | 0.10052 | 0.08797 | Low Total |
| Po. icarus | 0.52338 | 0.53562 | 1.02338 | 0.01297 | 0.01183 | 1.09648 | -0.01095 | 0.03689 | 0.27959 | Low Homozygous |
| Po. icarus | 2.55575 | 2.50478 | 0.98006 | -0.05097 | 0.03091 | -1.64873 | -0.1135 | 0.01156 | 0.10724 | Moderate Total |
| Po. icarus | 0.53694 | 0.53737 | 1.0008 | 0.0043 | 0.01264 | 0.3402 | -0.02127 | 0.02987 | 0.73553 | Moderate Homozygous |
| Pl. argus | 0.20199 | 0.23058 | 1.14158 | 0.0286 | 0.00984 | 2.90469 | 0.00859 | 0.0486 | **0.00642** | Conserved Homozygous |
| Pl. argus | 2.1471 | 2.12135 | 0.98801 | -0.0189 | 0.03979 | -0.47497 | -0.09977 | 0.06197 | 0.63784 | Conserved Total |
| Pl. argus | 0.097 | 0.09619 | 0.99166 | -0.00025 | 0.00581 | -0.04346 | -0.01205 | 0.01155 | 0.96559 | High Total |
| Pl. argus | 0.01984 | 0.02051 | 1.03376 | 0.00072 | 0.00302 | 0.23704 | -0.00542 | 0.00685 | 0.81405 | High Homozygous |
| Pl. argus | 39.8359 | 39.84086 | 1.00012 | -0.1089 | 0.13944 | -0.781 | -0.39227 | 0.17447 | 0.44021 | Intergenic Total |
| Pl. argus | 9.10907 | 9.66486 | 1.06101 | 0.61549 | 0.16653 | 3.69599 | 0.27706 | 0.95392 | **0.00077** | Intergenic Homozygous |
| Pl. argus | 6.62767 | 6.6161 | 0.99825 | 0.02418 | 0.08958 | 0.26991 | -0.15788 | 0.20624 | 0.78886 | Low Total |
| Pl. argus | 1.93521 | 1.97182 | 1.01892 | 0.06686 | 0.05117 | 1.30673 | -0.03712 | 0.17085 | 0.20008 | Low Homozygous |
| Pl. argus | 6.99366 | 6.73786 | 0.96342 | -0.2398 | 0.06228 | -3.85045 | -0.36637 | -0.11324 | **5.00E-04** | Moderate Total |
| Pl. argus | 1.81401 | 1.83072 | 1.00922 | 0.02756 | 0.04659 | 0.59166 | -0.06711 | 0.12224 | 0.55799 | Moderate Homozygous |
| Cy. semiargus | 0.20064 | 0.26764 | 1.33396 | 0.06612 | 0.02301 | 2.87343 | 0.01954 | 0.11271 | **0.00661** | Conserved Homozygous |
| Cy. semiargus | 2.00528 | 2.04377 | 1.01919 | 0.0382 | 0.02947 | 1.29633 | -0.02146 | 0.09787 | 0.20268 | Conserved Total |
| Cy. semiargus | 0.08421 | 0.08648 | 1.02693 | 0.00196 | 0.00536 | 0.36607 | -0.00889 | 0.01281 | 0.71634 | High Total |
| Cy. semiargus | 0.02213 | 0.02681 | 1.21112 | 0.00467 | 0.00536 | 0.87202 | -0.00618 | 0.01552 | 0.38867 | High Homozygous |
| Cy. semiargus | 52.48658 | 52.95155 | 1.00886 | 0.44782 | 0.29089 | 1.53945 | -0.14107 | 1.0367 | 0.13198 | Intergenic Total |
| Cy. semiargus | 11.10881 | 14.329 | 1.28988 | 3.22019 | 1.05601 | 3.04939 | 1.08241 | 5.35798 | **0.00416** | Intergenic Homozygous |
| Cy. semiargus | 5.14493 | 5.1522 | 1.00141 | 0.04006 | 0.0679 | 0.59001 | -0.0974 | 0.17753 | 0.55867 | Low Total |
| Cy. semiargus | 1.24445 | 1.53844 | 1.23624 | 0.29481 | 0.09787 | 3.01218 | 0.09668 | 0.49294 | **0.0046** | Low Homozygous |
| Cy. semiargus | 5.85531 | 5.81692 | 0.99344 | -0.0236 | 0.07358 | -0.32076 | -0.17256 | 0.12535 | 0.75015 | Moderate Total |
| Cy. semiargus | 1.32953 | 1.66162 | 1.24978 | 0.32898 | 0.10274 | 3.20196 | 0.12099 | 0.53698 | **0.00276** | Moderate Homozygous |

**Table S12. Study species conservation status and reference genome information.** For each study species, the conservation status in Sweden is presented based on the most recent assessment, the 2020 Swedish Red List. Sequencing data for each species was mapped to the closest available chromosome level reference genome. For the three focal species, and all but one of the additional species, this was a species specific genome. For *Plebejus argyrognomon* , the closest reference (*Pl. argus* ) was used. For inbreeding and GONE analyses, we utilized a recombination rate based off the assumption of 50 centimorgans per chromosome, which were used to calculate the autosomal recombination rates for the reference genome species in Morgans per base pair per generation.

| Study Species | Swedish Red List 2020 Status | Reference species | RefSeq Assembly Accession | Assembly Name | Autosomal Chromosomes | Autosomal Length (bp) | Recombination rate (M/bp) |
|---|---|---|---|---|---|---|---|
| *Polyommatus icarus* | LC (Livskraftig; Least Concern) | *Polyommatus icarus* | GCA_937595015.1 | ilPolIcar1.1 | 22 | 472204672 | 2.3295E-08 |
| *Plebejus argus* | LC (Livskraftig; Least Concern) | *Plebejus argus* | GCA_905404155.3 | ilPleArgu1.3 | 22 | 360412310 | 3.05206E-08 |
| *Cyaniris semiargus* | LC (Livskraftig; Least Concern) | *Cyaniris semiargus* | GCA_905187585.1 | ilCyaSemi1.1 | 23 | 416757882 | 2.7594E-08 |
| *Agriades optilete* | LC (Livskraftig; Least Concern) | *Agriades optilete* | GCA_964273475.1 | ilAgrOpti3.hap1.1 | 23 | 607663052 | 1.8925E-08 |
| *Aricia agestis* | LC (Livskraftig; Least Concern) | *Aricia agestis* | GCF_905147365.1 | ilAriAges1.1 | 22 | 394696725 | 2.78695E-08 |
| *Cupido Minimus* | NT (Nära hotad; Near Threatened) | *Cupido minimus* | GCA_965195375.1 | ilCupMini1.1 | 23 | 448207675 | 2.56577E-08 |
| *Plebejus argyrognomon* | CR (Akut hotad; Critically endangered) | ***Plebejus argus**** | GCA_905404155.3 | ilPleArgu1.3 | 22 | 360412310 | 3.05206E-08 |
| *Scolitantides orion* | EN (Starkt hotad; Endangered) | *Scolitantides orion* | GCA_964345685.1 | ilScoOrio1.hap1.1 | 22 | 468110494 | 2.34987E-08 |

**Table S13. Species dataset sequence data filtering.** For each species dataset, we generated a 'filtered sites' file to limit analyses to. This filtered sites file is the intersection of several independent filters that remove scaffold smaller than 1Mb, non-autosomal contigs, repetitive regions, and regions in the top and bottom 1% of global sequencing depth for all sample, museum sample, and fresh sample groupings. Here, the number of base pairs and the proportion of the genome passing each filter is shown for each of the species datasets, as well as these parameters for the final combined filtered sites file. In some cases, only museum samples were utilized for a species, in which case the depth filters are only performed on one sample grouping, which is the same as the filtering across all samples. The reference genome used for each species is listed at the top of the table, and has a * when it is from a different species to the study species.

| | *Polyommatus icarus* (ilPolIcar1.1) | | *Plebejus argus* (ilPleArgu1.3) | | *Cyaniris semiargus* (ilCyaSemi1.1) | | *Aricia agestis* (ilAriAges2.1) | |
|---|---|---|---|---|---|---|---|---|
| **Filter** | **Filtered Length(bp)** | **Filtered percent** | **Filtered Length(bp)** | **Filtered percent** | **Filtered Length(bp)** | **Filtered percent** | **Length(bp)** | **Percent** |
| *Total genome* | 511758028 | 100 | 382108302 | 100 | 441519306 | 100 | 437383100 | 100 |
| *Scaffolds<1000000bp* | 511184185 | 99.8879 | 379254911 | 99.2533 | 441467852 | 99.9883 | 437193358 | 99.9566 |
| *Autosomes* | 472204672 | 92.2711 | 363250311 | 95.0648 | 416794061 | 94.4 | 394871022 | 90.2804 |
| *Repeats* | 230539290 | 45.0485 | 207343925 | 54.2631 | 217090499 | 49.169 | 219807041 | 50.255 |
| *Depth (all)* | 428236719 | 83.6795 | 328918008 | 86.0798 | 388536918 | 88 | 335802494 | 76.7754 |
| *Depth (museum)* | 355099842 | 69.3882 | 328310056 | 85.9207 | 388068388 | 87.8939 | 329841456 | 75.4125 |
| *Depth (fresh)* | 427116071 | 83.4606 | 269759492 | 70.5977 | 347302061 | 78.6607 | 273127265 | 62.4458 |
| **Combined** | 201529665 | 39.3799 | 186409699 | 48.7845 | 195370411 | 44.2496 | 179516799 | 41.0434 |

| | *Agriades optilete* (ilAgrOpti3.hap1.1) | | *Cupido minimus* (ilCupMini1.1) | | *Plebejus argyrognomon* (ilPleArgu1.3*) | | *Scolitantides orion* (ilScoOrio1.hap1.1) | |
|---|---|---|---|---|---|---|---|---|
| **Filter** | **Length(bp)** | **Percent** | **Length(bp)** | **Percent** | **Length(bp)** | **Percent** | **Length(bp)** | **Percent** |
| *Total genome* | 680738285 | 100 | 474403353 | 100 | 382108302 | 100 | 525334176 | 100 |
| *Scaffolds<1000000bp* | 677998561 | 99.5975 | 474293063 | 99.9768 | 379254911 | 99.2533 | 524519661 | 99.845 |
| *Autosomes* | 607663052 | 89.2653 | 448207675 | 94.4782 | 363250311 | 95.0648 | 468110494 | 89.1072 |
| *Repeats* | 231683966 | 34.0342 | 211975703 | 44.6826 | 207343925 | 54.2631 | 210604438 | 40.0896 |
| *Depth (all)* | 520387915 | 76.4446 | 345703778 | 72.8713 | 142493774 | 37.2915 | 381912722 | 72.699 |
| *Depth (museum)* | 517527179 | 76.0244 | 340141899 | 71.6989 | 142493774 | 37.2915 | 381912722 | 72.699 |
| *Depth (fresh)* | 377891541 | 55.512 | 279749861 | 58.9688 | NA | NA | NA | NA |
| **Combined** | 182033741 | 26.7406 | 174264760 | 36.7335 | 124475234 | 32.5759 | 176286414 | 33.557 |

**Table S14. Removal of close-relatives from admixture/PCA.** As close relatives can skew the results of principal component and admixture analyses, we removed third-degree and closer relatives from these analyses. We estimated relatedness using the IBSrelate method, identifying first degree (PO/FS; KING 0.25), second-degree (HS; KING 0.125), and third-degree (C1; KING 0.0625) relatives and removing one of each pair from the analyses. The removed individuals are highlight in bold and red text. These individuals were included in all other analyses.

| Species | Sample 1 | Sample 2 | R0 | R1 | KING | Inferred Relationship |
|---|---|---|---|---|---|---|
| *Po. icarus* | **PI031** | PI033 | 0.2633 | 0.291742 | 0.0917 | C1 |
| *Pl. argus* | **MZLU107435** | MZLU107438 | 0.2510 | 0.2997 | 0.0979 | C1 |
| *Cy. semiargus* | **MZLU107455** | MZLU107461 | 0.025537 | 0.780051 | 0.291401 | FS |
| *Cy. semiargus* | MZLU153216 | **MZLU153221** | 0.073815 | 0.605471 | 0.238239 | FS |
| *Cy. semiargus* | CYSE0078 | **CYSE0098** | 0.212413 | 0.323411 | 0.117873 | HS |
| *Cy. semiargus* | MZLU107456 | **MZLU107501** | 0.237071 | 0.346552 | 0.113124 | HS |
| *Cy. semiargus* | CYSE0198 | **CYSE0199** | 0.271307 | 0.290711 | 0.088494 | C1 |

**Table S15. Reference genomes utilized for estimating GERP scores.** We utilized the method implemented in the GenErode pipeline (116) to estimate GERP scores and ancestral states across the reference genomes for each species dataset. Below is the list of reference genomes, their accessions, and the reference species. We utilized all genomes from this list when estimating GERP scores and ancestral states along each of the focal species reference genomes, excluding the focal species.

| RefSeq Assembly Accession | Assembly Name | Organism_Name | Genome Note |
|---|---|---|---|
| GCA_964273475.1 | ilAgrOpti3.hap1.1 | Agriades optilete | NA |
| GCF_905147365.1 | ilAriAges1.1 | Aricia agestis | NA |
| GCA_965178315.1 | ilCalAvis1.hap1.1 | Callophrys avis | NA |
| GCA_905187575.2 | ilCelArgi3.2 | Celastrina argiolus | (118) |
| GCA_965195375.1 | ilCupMini1.1 | Cupido minimus | NA |
| GCA_905187585.1 | ilCyaSemi1.1 | Cyaniris semiargus | (87) |
| GCA_964396525.1 | ilAriEume1.hap1.1 | Eumedonia eumedon | NA |
| GCA_905404095.1 | ilGlaAlex1.1 | Glaucopsyche alexis | (119) |
| GCA_963853865.1 | ilHelHell1.1 | Helleia helle | NA |
| GCA_964662265.1 | ilQueQuer1.hap2.1 | Hypaurotis quercus | NA |
| GCA_965112825.1 | ilIolDebi1.hap1.1 | Iolana debilitata | NA |
| GCA_964264435.1 | ilKreTrap1.hap1.1 | Kretania trappi | NA |
| GCA_964417165.1 | ilLaeRobo1.hap1.1 | Laeosopis roboris | NA |
| GCA_964656195.1 | ilLamBoet1.hap1.1 | Lampides boeticus | NA |
| GCA_965112865.1 | ilLepPiri1.hap1.1 | Leptotes pirithous | NA |
| GCA_905333005.2 | ilLycPhla1.2 | Lycaena phlaeas | (117) |
| GCA_905333045.1 | ilLysBell1.1 | Lysandra bellargus | (31) |
| GCA_963565745.1 | ilPheArio1.1 | Phengaris arion | NA |
| GCA_905404155.3 | ilPleArgu1.3 | Plebejus argus | (86) |
| GCA_937595015.1 | ilPolIcar1.1 | Polyommatus icarus | (85) |
| GCA_963422495.1 | ilPolIphe1.1 | Polyommatus iphigenia | NA |
| GCA_965178345.1 | ilSatIlic2.hap1.1 | Satyrium ilicis | NA |
| GCA_964345685.1 | ilScoOrio1.hap1.1 | Scolitantides orion | NA |
| GCA_965153325.1 | ilTomBall1.hap1.1 | Tomares ballus | NA |
